# Nuclear quantum effects on ion permeation through the open transient receptor potential melastatin 7 pore: A continuum-electrostatics estimate

**DOI:** 10.64898/2026.11.18.767574

**Authors:** Yizeng Li, Hsiu-Hau Lin

## Abstract

We estimate how nuclear delocalization affects magnesium and calcium release from occupied wells in the naltriben-open pore of transient receptor potential melastatin 7 (TRPM7). We compare the equilibrium distributions of sodium, magnesium, and calcium in the same open structure and fixed continuum field. Thermal quantum states extend over nearby positions, so the ion can sample more than one local environment. For the divalent ions, the model combines a Gaussian test charge in the nonlinear Poisson–Boltzmann potential, spatial dielectric self-energy, and the free-energy cost of restricting a correlated first hydration shell. The hydration shell comprises the water molecules that directly coordinate the ion. In a constriction, the wall can block access to some of these water sites. The classical and quantum calculations use the same insertion energy, pore geometry, and axial diffusivity. The leading Wigner–Kirkwood correction for a smooth ion energy *U*_*i*_ away from the wall is 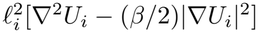, up to an additive constant, with 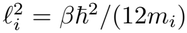 for ion mass *m*_*i*_ and inverse thermal energy *β*. The length *l*_*i*_ is about 0.02 Å at 310 K. At the point-ion magnesium basin, the refined centerline self-energy reaches 430 meV. Steric constraints also restrict water coordination at the 1.43 Å constriction. At 310 K in the primary extended dielectric box, the deepest wells for magnesium and calcium lie at 22.75 and 11.00 Å, respectively. The quantum changes in mean passage time toward the intracellular and extracellular interfaces are 0.68% and −0.66% for magnesium and −0.10% and 16.12% for calcium. Calcium starts on the intracellular side of the constriction and must cross it to reach the extracellular interface. The preferred magnesium site is already on the extracellular side. A change in the dielectric boundary for magnesium or the ligand penalty for calcium can move the deepest well across the constriction and reverse the extracellular response. For comparison, the circular point-ion estimate for magnesium changes from −37.4% and −34.5% to −8.9% and −8.6% with a 0.5 Å wall inset. In the circular reference, the largest local correction occurs at the intracellular widening and depends on near-wall access and axial sampling. Its influence on escape from the occupied well is small. In a separate screened two-ion benchmark, classical repulsion changes escape barriers by tens to hundreds of meV. Isotope substitution tests the mass dependence at fixed charge and chemistry. Together, the results show how pore geometry, dielectric response, and water coordination determine which local quantum corrections affect directional release.

## 1 Introduction

The open pore of transient receptor potential melastatin 7 (TRPM7) provides an aqueous pathway for ions across the cell membrane [1]. Electrophysiology has established that the channel conducts magnesium and calcium, and intestinal deletion of TRPM7 in mice disrupts the balance of several divalent minerals [2–4]. Cytosolic magnesium also regulates opening [5]. Mutagenesis, electrophysiology, and simulations implicate a magnesium-binding site near the lower gate in stabilizing the closed state [5]. Structural studies place glutamate 1047 (E1047), a residue that strongly affects divalent permeation when mutated, at the selectivity filter and place tyrosine 1085 (Y1085) at a constriction nearer the inside of the cell [1, 3, 6]. Classical molecular dynamics simulations have described the hydration of the pore and cation passage in the open N1098Q mutant [1]. We focus on the movement of an ion through the conformation opened by the agonist naltriben. This open-state comparison isolates a step after ion entry. The open pore can hold an ion at a favorable site, and release then depends on the barriers between that site and an exit.

Within an open TRPM7 pore, the chemical environment sets the positions at which a divalent ion can reside and the route it takes to leave. A favorable region can promote ion occupancy and create an energetic cost for escape [7, 8]. The axial free energy measures the statistical weight of the configurations accessible across each section of the pore [7]. A cross-section can favor occupancy by offering more positions to the ion or by making some of those positions especially favorable [7]. In gramicidin, explicit-water simulations show that ions lose part of their hydration shell as they enter the pore [9]. Water within the pore then supplies a substantial part of the stabilization for the permeating ion [9]. The widths of occupied regions and barriers, together with the local rate of diffusion, also influence passage [8, 10].

Nuclear quantum effects reflect the quantum nature of nuclear motion [11]. We consider the equilibrium effect of the direct delocalization of the permeating ion. In classical equilibrium, lower-energy configurations carry greater statistical weight [7]. Quantum states extend in space, so the quantum density at a position also responds to the energies and confining boundaries nearby [11–14]. In water, the motion of light hydrogen nuclei contributes to quantum changes in liquid structure, kinetic energy, and equilibrium isotope fractionation near dissolved ions [15, 16]. The three permeating ions considered here have masses at least twenty times that of a proton [17]. Simulations of lithium and fluoride have also examined how these effects change hydration-shell structure and exchange [18]. Recent many-body simulations included nuclear quantum effects in the hydration free energies of bulk aqueous alkali and halide ions and found modest changes in their first-shell structure [19]. These studies motivate asking how the direct quantum contribution of the ion changes when both its surroundings and the available space vary along a pore.

A quantum model of the KcsA potassium channel has examined the delocalization of a single K^+^ ion in the selectivity filter [20]. Molecular-dynamics analyses have also proposed a connection between correlated ion motion and quantum coherence in transport [21]. We examine the equilibrium redistribution caused by thermal delocalization and test how its location within the pore determines the estimated effect on ion passage.

The thermal de Broglie wavelength gives a scale for the spatial extent of quantum effects. At 310 K, it is 0.21, 0.20, and 0.16 Å for sodium, magnesium, and calcium, respectively. In a circular cross-section with an impenetrable boundary, or hard wall, the leading correction due to confinement is small when this wavelength is short relative to the pore diameter [14]. A second length scale is the distance over which the energy changes. For a smooth potential, the leading local quantum correction depends on its squared gradient and curvature. Both terms are proportional to 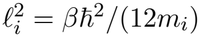, with *l*_*i*_ ≃0.02 Å for the three ions at 310 K [12, 13, 22]. In the TRPM7 model, a change in radius over a short axial interval can expose low-energy positions away from the pore axis. Those positions can carry enough equilibrium weight to produce a steep change in free energy even where the pore is wide. We calculate separate contributions from motion across the pore and motion along its axis to resolve this effect.

We use the structure of the naltriben-open channel in Protein Data Bank (PDB) entry pdb_00008si5 [23]. At fixed pore geometry and field, we isolate the thermal delocalization of the center of mass of a single ion. We pair selective changes in wall geometry with a decomposition of the passage integral to determine how quantum redistribution near the entrance, basin, and exit barriers contributes to escape.

For magnesium and calcium, we evaluate dielectric charging and the free energy of incomplete water coordination on the same ion-center domain used for the quantum trace. This comparison tracks how the same charge and water-coordination costs set the starting well and determine which parts of the pore the ion must traverse to escape. The fixed environment provides a reference for calculations that also quantize water and protein nuclei [11, 15, 18]. We treat the center of mass of a single ion in the primary calculation. Exchange between identical ions is considered separately in assessing the scope of the two-ion benchmark.

## 2 Model and calculations

### 2.1 Accessible pore geometry

The pore radius defines how far the center of an ion can move from the axis at each position along the channel. We use the same axis and accessible domain in both the classical and quantum calculations so that any difference arises from the statistical treatment of ion motion.

We used the four-subunit protein in biological assembly 1 of `pdb_00008si5`, the naltriben-open structure of mouse TRPM7 [1, 23]. We located the pore axis by fitting a line through the centers of rings formed by corresponding *α*-carbon atoms (C*α*) in the four subunits. We used the membrane interfaces from the Orientations of Proteins in Membranes (OPM) database to define the modeled interval. The coordinate *z* increases from *z* = 0 on the intracellular side to *z* = *L* = 30.0 Å (3.0 nm) on the extracellular side [24] (Fig. 1A).

**Figure 1:**
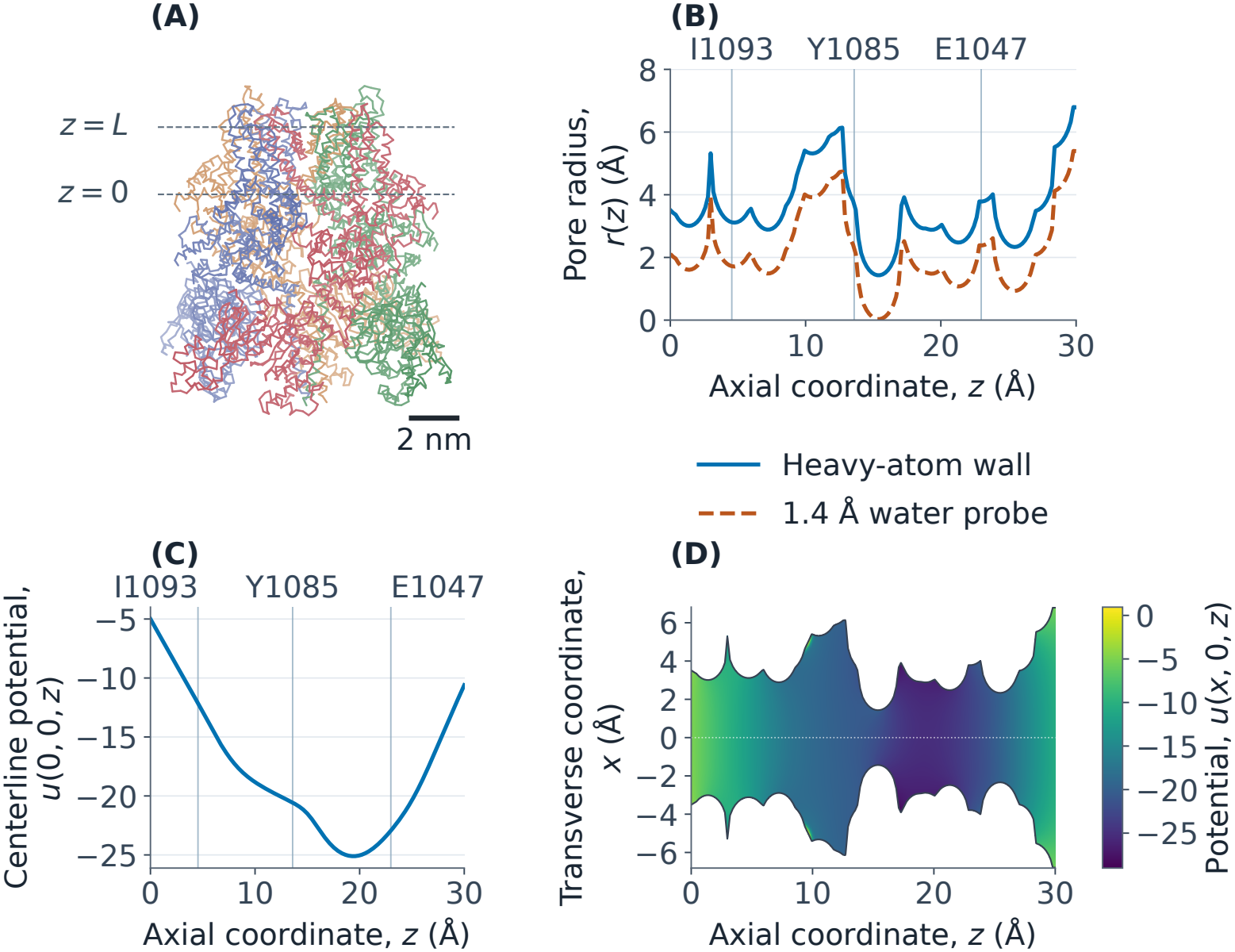
Open TRPM7 pore, accessible radius and electrostatic potential. (A) Backbone traces of the four subunits with projections of the intracellular interface (*z* = 0) and extracellular interface (*z* = *L*). (B) Centered van der Waals radius *r*(*z*) of the structural disk along the axial coordinate *z*. The lower curve is the center clearance max[*r*(*z*) − 1.4 Å, 0] for a 1.4 Å solvent probe. (C) Dimensionless centerline potential *u*(0, 0, *z*) = *eϕ*(0, 0, *z*)*/*(*k*_B_*T*_PB_) at *T*_PB_ = 310 K. (D) Potential *u*(*x*, 0, *z*) in the axial section *y* = 0, clipped at the accessible radius *r*(*z*). Vertical lines mark I1093, Y1085 and E1047.

At each *z*, a structural disk of radius *r*(*z*) sets the outer envelope for the accessible domain Ω(*z*) of the ion center. The circular point-ion reference uses the full disk with area *A*(*z*) = *πr*(*z*)^2^. We set *r*(*z*) to the distance from the axis to the nearest van der Waals surface of a protein heavy atom in that cross-section. We retain the gaps left by unresolved residues in the structural model. The coverage and charge assignments at these gaps are described below. The radius reaches a minimum of 1.43 Å at *z* = 15.50 Å (Fig. 1B). Linear interpolation makes the tabulated wall continuous with slope changes at the radius planes. Some planes lie where different atoms control the clearance, including the sharp widening near *z* = 3 Å. A second rapid change occurs farther into the pore, where the tabulated radius decreases from 6.14 to 4.68 Å between *z* = 12.72 and 12.91 Å. The transverse wave function vanishes at the wall.

Three residues locate structural features along the axis. Starting from the intracellular side, isoleucine 1093 (I1093) lies in the lower gate, Y1085 lines a constriction, and E1047 contributes to the selectivity filter [1, 3]. Each reference plane passes through the mean axial position of the four corresponding C*α* atoms. The planes are at 4.54, 13.59, and 22.98 Å, respectively (Fig. 1B). The radius calculation uses the surrounding heavy atoms, so side chains can affect the accessible radius at positions beyond their reference planes.

For the magnesium geometry control, we construct a centered circular wall that clears every positive-radius PARSE sphere in the electrostatic model. For each interval between adjacent knots, a sphere has its largest cross-section on the plane closest to its center. We subtract the radius of that cross-section from the distance between the center and the pore axis. The smallest of these clearances across the intersecting spheres sets the upper bound on a constant radius that avoids sphere overlap throughout the interval. We cap both endpoints by that radius, with a numerical clearance, so linear interpolation also clears the spheres between knots. A further reduction where needed bounds the magnitude of the slope by that of the steepest segment of the structural profile. The resulting disk can be smaller than the full irregular sphere-free cross-section. The irregular-domain calculation below retains the remaining accessible space around the axis.

We counted 1,097 modeled *α*-carbon positions per subunit across numbered residues 3–1230, equivalent to 89.3% of that numbered span. The internal gaps are 541–613, 790–838, and 1148–1156 in each subunit. The smallest radial distance from any gap-flanking *α* carbon to the pore axis is 43.5 Å. The prepared charge is +10*e* per chain. We screened the flanks for atom names associated with terminal groups, including OXT and additional N-terminal hydrogens, and found no such markers. This coordinate and charge audit locates the missing segments relative to the pore and documents the fixed-charge model that enters the electrostatic solve. The archived PDB2PQR log flags all 12 internal breaks. At each of the 24 flanking residues, the atom names and the individual charge assignments match those of an internal residue of the same type in the same subunit. They also match between the exploratory and reference PQR files. Thus the final model retains internal-residue charge assignments at these gaps. The closest flank lies outside a 20 Å radius of the pore axis. The residue-level charge assignments and break detections accompany the structural model.

For a center at **r**, write *c*(**r**) for the minimum distance to the surfaces of the prepared atomic spheres, with a negative value for an interior point. At each plane the maximum over the structural disk is *c*_max_(*z*). We write *A*(*z*) for the accessible area, which equals *πr*(*z*)^2^ for the full structural disk.

The shortest clearance from a center to the atomic-sphere surfaces is a 1-Lipschitz function of position. We bounded its maximum over each structural disk using a transverse sampling mesh. The maximum at the nodes inside the disk is a lower bound. The maximum at all nodes within one half-diagonal of the disk plus one half-diagonal is an upper bound.

### 2.2 Electrostatic background and insertion energy

The geometry defines the positions an ion can reach. Of the positions the ion center can reach, some leave room for more coordinating waters than others. To assign an energy to each position, we use a fixed electrostatic field for the open channel. The field arises from the fixed protein charges in the dielectric environment of the protein, membrane, and water, with screening by salt in the surrounding solutions. The imposed voltage across the membrane is zero. Both thermal calculations use this same position-dependent energy.

We prepared the protein’s atomic charges and radii with PDB2PQR and estimated the initial protonation states with PROPKA [25–27]. We used PARSE atomic parameters and set the pH to 7.4 for the initial protonation estimate. The Adaptive Poisson–Boltzmann Solver (APBS) supplied the solution of the nonlinear Poisson–Boltzmann (PB) equation for the protein in a dielectric membrane with 150 mM sodium chloride in each reservoir [28, 29]. In the reference protonation state, all four E1047 side chains are deprotonated. This gives the same carboxylate charge in each subunit at the selectivity filter. The relative permittivities of protein, membrane, and water are 4, 2, and 74.2433, respectively, with the water value at 310 K. A full-protein field supplies the boundary values for the nested pore calculations (Supplementary Materials). The pore has water-like permittivity with smooth transitions at its boundary. The membrane reduces mobile-ion accessibility, and the hydrated pore excludes mobile ions from the screening term. The bare point-ion reference uses the energy *U*_*i*_ = *Z*_*i*_*eϕ*(**r**), where *Z*_*i*_ is its signed valence, *e* is the elementary charge, and *ϕ*(**r**) is the molecular electrostatic potential at position **r**, calculated at *T*_PB_ = 310 K. We hold *U*_*i*_ fixed as we vary the ion’s thermal temperature *T* to isolate the effect of temperature on its distribution. For the dimensionless field *u* = *eϕ/*(*k*_B_*T*_PB_), the interaction is *U*_*i*_ = *Z*_*i*_*k*_B_*T*_PB_*u*. The divalent test ions thus have twice the monovalent interaction energy at each point of the common field.

Along the axis, the potential is −5.00 *k*_B_*T*_PB_*/e* at the intracellular interface, reaches a minimum of − 25.11 *k*_B_*T*_PB_*/e* near *z* = 19.37 Å, and rises to −10.61 *k*_B_*T*_PB_*/e* at the extracellular interface (Fig. 1C). A plane through the pore shows lateral variations in the potential, especially near the entrances (Fig. 1D).

This test-charge energy describes the ion’s interaction with the fixed environment. The reference uses the electrostatic potential before the ion is inserted. The test ion therefore enters only through its interaction energy, and the same background potential serves all test-ion charges. The inserted ion polarizes the dielectric medium, and rearrangement of its hydration shell changes its insertion free energy [30, 31]. The primary insertion energy for magnesium and calcium is

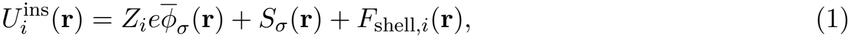

where 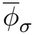 averages the fixed molecular potential over a normalized Gaussian charge distribution of standard deviation *σ* = 0.38 Å with spherical cutoff 3*σ*. The dielectric cost *S*_*σ*_ measures the electrostatic charging energy relative to that of the same ion charge in water. At fixed charge shape and in linear dielectric response, this cost scales with 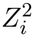. The hydration cost depends on the geometry and accessibility of the ligand sites. We interpolate the reaction energy from axial and off-axis Gaussian-probe solves, with separate numerical checks on mesh spacing and boundaries. The shell term *F*_shell,*i*_ averages steric access for a discrete water geometry over common rotations of its ligands. The reference counts are six for magnesium and eight for calcium, and the ion–oxygen distances are 2.09 and 2.44 Å, respectively [32–34]. The finite-ion domain retains positions inside the structural disk with at least 1.15 Å clearance from the prepared atomic spheres. This choice permits the entire truncated Gaussian support. The shell calculation uses a separate 1.40 Å water probe and a prescribed free-energy cost for an obstructed ligand site. The hydration model below defines the shell orientation integral and its coordination and steric parameters.

Let ε_*ε*_ denote the charging energy of a divalent Gaussian charge distribution of standard deviation 0.38 Å in the heterogeneous dielectric, and let ε_w_ denote the same energy in water. The difference *S* = ε_*ε*_ − ε_w_ gives the reaction-field contribution [31]. We first construct two approximations on the archived dielectric maps. Let *D*_w,*j*_ be the component of the homogeneous-water displacement field of the trial charge, with *j* running over Cartesian directions. Evaluation of the electrostatic potential functional at the water solution gives a lower variational energy *S*_L_. Evaluation of the displacement functional at the same water field gives an upper variational energy *S*_U_. These trial-field bounds follow from the complementary electrostatic variational principles [35]. For directional relative permittivities *ε*_*j*_ these expressions are

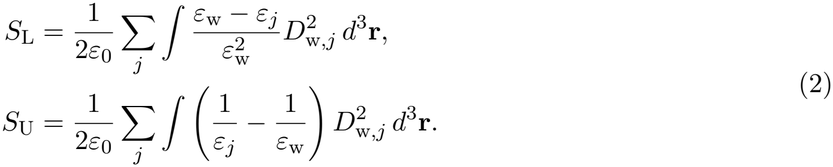

The potential functional reaches its maximum at the electrostatic solution. The complementary displacement functional reaches its minimum under Gauss’s law. Thus the inequalities apply to the continuous calculation for the common unbounded water reference and the chosen Gaussian. We represent the medium outside the coefficient box by water in these integrals. The Gaussian displacement kernel is analytic and the integrals are evaluated by convolution on the staggered 0.25 Å maps. The discretized integrals use the analytic Gaussian. The direct probes use a Gaussian truncated at three standard deviations, on a finite Dirichlet box. The variational fields provide the spatial variation of the interpolation. Direct probe comparisons quantify its accuracy within each finite box.

The axis-calibrated control uses *S*_L_ + *θ*(*z*)(*S*_U_ − *S*_L_). We write 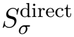 for a direct-probe self-energy and 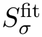 for its axis-calibrated estimate. We obtained *θ* by matching independent on-axis dielectric-only finite-difference solves and interpolated it with a shape-preserving cubic. The direct solves subtract the charging energy of the same discretized source in water, with zero potential at the box faces in both media. The test-ion background energy is averaged over the truncated Gaussian in the Gaussian-background control. We retain the point background in separate comparisons to isolate the effect of the dielectric and shell terms.

We sampled the regularized dielectric Green function with separate lateral and axial domain comparisons. Each source carried charge +2*e* in a Gaussian of standard deviation 0.38 Å, truncated at three standard deviations and normalized on the mesh. We solved the dielectric equation with zero Dirichlet values on all six faces. The same source, mesh, and boundary conditions define the homogeneous-water charging energy. We invert the homogeneous-water finite-difference operator in a separable discrete sine basis. A direct sparse solve on a small three-dimensional box verifies the evaluation. The background molecular potential is averaged over the truncated Gaussian for the thermal calculation. The dielectric charging calculation concerns polarization of the fixed medium. The nonlinear Poisson–Boltzmann background retains reservoir screening.

The calibration uses centerline and off-axis probes. Radial positions are measured relative to the first excluded position at the guarded sphere clearance or the structural disk boundary. The training set uses the fourfold rotations of these positions. Independent probes at distinct azimuths and depths, near low-energy positions, and at rotated counterparts of training sites were reserved for comparison. All Gaussian supports clear the prepared atomic spheres.

We use the variational energies in Eq. 2 to supply the short-range spatial variation. At each calibration probe we form *θ* = (*S*_*σ*_− *S*_L_)*/*(*S*_U_ −*S*_L_) and clip it to [0, 1]. The spatial spline and field-sampling procedure are specified in the Supplementary Materials.

Adding a constant to the potential adds the same energy at every position for a given ion. In either thermal calculation, this multiplies all equilibrium weights by the same factor, which cancels when the weights are normalized. The relative free energies, exit barriers, and passage-time ratios are thus invariant to a uniform potential shift. Each passage route ends at a membrane interface with a potential distinct from the bulk reference. The barriers and times describe escape from the modeled membrane slab.

At the intracellular and extracellular interfaces the dimensionless potential in the fine field is −5.00 and −10.61, respectively. At the two longitudinal faces of the fine box (*z* = −5 and 35 Å) it is +0.18 and −3.60. At the corresponding faces of the parent box (*z* = −15 and 45 Å) it is +1.05 and −1.11. On the 2.0 Å outer grid, the sampled centerline satisfies |*u*| ≤ 0.1 continuously from *z* ≃−117 and 63 Å outward to the corresponding box faces. This criterion diagnoses recovery of the electrostatic bulk reference along the axis. A reservoir also requires a connected solvent-accessible region. The focused grids and the circular thermal domain define the membrane interval used for the escape calculation. Extending the thermal domain into the reservoirs requires an accessible vestibule cross-section and a bath boundary condition.

We separately computed the straight-axis clearance by subtracting each prepared atomic radius from the distance to its center and taking the minimum. Between the intracellular interface and the |*u*| ≤0.1 region, the sampled clearance reaches 0.51 Å near *z* = −84 Å. Intersections of the line with spheres expanded by 1.15 Å give seven intervals that the finite ion center cannot traverse on the axis. On the extracellular side, the smallest clearance between the interface and the corresponding bulk criterion is 4.34 Å. The tabulated centerline electrostatics therefore reach the bulk reference on both sides. A finite-ion path to the intracellular bath must allow lateral motion outside the slab. The clearance table describes the existing structure along the axis, while the escape estimates retain the membrane interfaces as endpoints.

### 2.3 Water coordination and shell access

We represent the magnesium shell by six oxygen positions on an octahedron and the calcium shell by eight positions on a square antiprism. Bulk structural data motivate these coordination geometries and the representative metal–oxygen distances of 2.09 and 2.44 Å [32–34]. Constrained first-principles calculations find a stable sixfold minimum for magnesium and a broader coordination distribution for calcium [34]. We also test a seven-ligand calcium shell with pentagonal bipyramidal geometry. All sites share the same radius within each geometry.

For ion center **r** and shell rotation ℛ, the *l*th oxygen center is at **r** + ℛ **a**_*il*_. Let *d* be its minimum clearance over the prepared atomic spheres. We define *a*_*il*_ = [1 + exp(− (*d* −*a*_w_)*/w*)]^*−*1^, with water probe radius *a*_w_ = 1.40 Å and smoothing length *w* = 0.12 Å. The ion center uses a separate clearance criterion. A smaller water radius of 0.70 Å supplies a steric sensitivity case. For a dimensionless penalty *η* associated with an obstructed site, the orientational free energy is

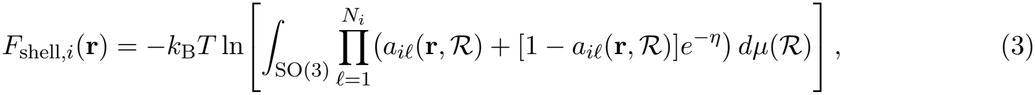

where *dµ* is the normalized uniform rotation measure. The product uses a single rotation for the entire shell. The shell free energy approaches zero as all site-access factors approach unity. The Boltzmann factor can also be written as *e*^*−η*^ + (1− *e*^*−η*^)*a*_*il*_, with the first term representing an unoccupied site. In the fully accessible limit this prior gives *P* (*N*_*i*_ −1)*/P* (*N*_*i*_) = *N*_*i*_*/*(*e*^*η*^− 1). The primary calculation uses *η* = 4, with comparisons at 2 and 6. These prescribed penalties control the cost of incomplete water coordination. Site-specific water exchange and replacement by protein ligands require additional free-energy information. The model keeps the continuum dielectric field fixed as the water configuration varies.

In a matched independent-ligand control, we first average the single-site access factor over all rotations and then take its *N*_*i*_th power. This keeps the same steric criterion, single-site access, penalty, and dielectric field.

For the separate control with independent ligand directions, let *f*_*i*_(**r**) be the fraction of uniformly sampled antipodal directions for which an oxygen center, placed at the representative first-shell distance from **r**, lies outside the prepared atomic spheres. We interpolate this fraction to the thermal positions. The bulk distances are 2.09 Å for magnesium and 2.44 Å for calcium. We use six and eight sites, respectively [32]. For independent ligand directions and an energy increment *ηk*_B_*T* for an obstructed direction, the insertion free energy relative to bulk is

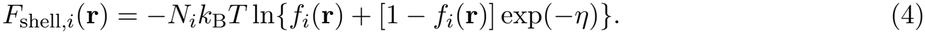

We use *η* = 2 at 310 K and compare *η* = 0 and 4. Each obstruction changes an effective ligand free energy. This independent-site control assigns angular access at a fixed coordination radius, with residue-specific exchange energies left for an explicit ligand model. These cases test the consequences of angular access around the ion. They retain the first-shell distance as a sampling length even in partially obstructed configurations.

### 2.4 Equilibrium free energy and exit barriers

We calculate the axial free energy in two steps. First, at each *z*, we combine the equilibrium weights of the configurations across the pore. Their sum gives the transverse partition function. A wider slice admits more positions, and low energies give those positions more weight. The resulting transverse free energy sets the energy landscape for the second step, in which the ion moves along the axis.

For an ion of species *i* and mass *m*_*i*_ at a fixed axial position *z*, the transverse Hamiltonian and quantum partition function are

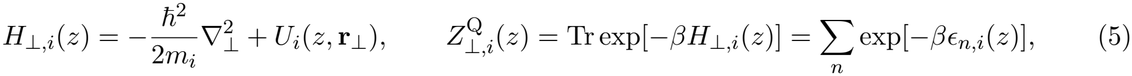

where **r**_*⊥*_ = (*x, y*) specifies position across the pore, 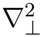 is the corresponding Laplacian, *ϵ*_*n,i*_ are the transverse energy eigenvalues with the hard-wall condition *ψ*| _*∂*Ω_ = 0, and *β* = 1*/*(*k*_B_*T* ) is the inverse thermal energy. For the finite-ion calculation we substitute 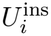 for *U*_*i*_ in the same statistical treatment. Here *k*_B_ is the Boltzmann constant. The trace Tr sums the Boltzmann weights exp(− *βϵ*_*n,i*_) of the quantum states. The classical counterpart integrates over transverse positions and momenta,

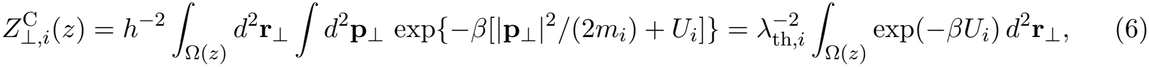

where **p** is the transverse momentum, 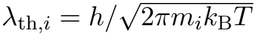 is the thermal de Broglie wavelength, *h* is Planck’s constant, and ℏ = *h/*(2*π*). Integration over momentum gives the prefactor 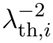, which is constant along the pore at a given mass and temperature. We convert each partition function to a transverse free energy, 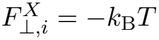 In 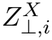, with *X* = C for the classical calculation and *X* = Q for the quantum calculation. A slice with greater partition weight has lower free energy.

For the second step, we use the quantum transverse free energy 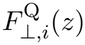 as the effective potential for motion along the axis in 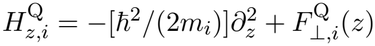. The diagonal matrix element of the thermal operator exp 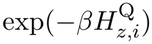 gives the unnormalized quantum density at *z*. The classical density uses the Boltzmann weight of 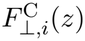. After normalization over *I* = [0, *L*], the two densities are

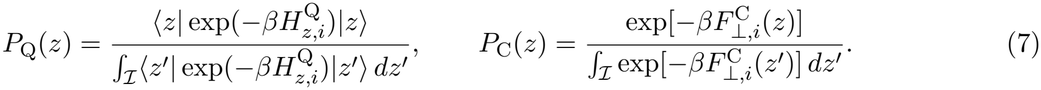

This reduction replaces the local transverse spectrum with a temperature-dependent scalar potential before axial quantization. It assumes local transverse thermal equilibrium in each slice. A three-dimensional treatment resolves the separate transverse mode energies and the kinetic couplings due to axial variation of the transverse states [36]. This reduction defines a matched classical–quantum comparison at fixed geometry and field, with the sharp entrance widening as the region most sensitive to the treatment of transverse and axial motion. We used masses of 22.9898, 24.305, and 40.078 u for sodium, magnesium, and calcium, respectively, with u denoting the unified atomic mass unit.

To place both axial free energies on the same scale, we choose a reference position *z*_*b*_ at a classical minimum. We use the global minimum within the modeled slab for the primary finite-ion calculation. The circular reference and the atomic-boundary and axis-calibrated sensitivity controls use the minimum between Y1085 and E1047. The companion finite-ion window calculation uses the same residue interval.

We call the selected starting point the basin position. We define *W*_*X*_(*z*) = −*k*_B_*T* ln[*P*_*X*_(*z*)*/P*_*X*_(*z*_*b*_)], so *W*_*X*_(*z*_*b*_) = 0 in both calculations. A negative correction Δ*W* = *W*_Q_− *W*_C_ means that the ratio of the density at that position to the density at the basin is greater in the quantum calculation.

We separate the contributions from motion across and along the pore by introducing an intermediate density 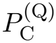. It is the classical axial Boltzmann density calculated with the quantum transverse free energy 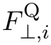 as its potential. Then Δ*W* = Δ*F*_*⊥*_ + Δ*W*_*z*_. The transverse term is the change in 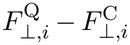 from *z*_*b*_ to *z*. The axial term is the corresponding change in −*k*_B_*T* ln 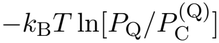.

For either exit direction *s*, we define the barrier as the largest free-energy rise from the basin, 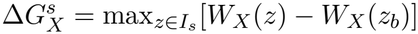, where *I*_*s*_ extends from the basin to the target membrane interface.

We find the maximum of each profile independently. The difference between the barrier heights is 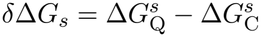. When both profiles have their directional maximum at the same position *z*_*s*_, the difference simplifies to *δ*Δ*G*_*s*_ = Δ*W* (*z*_*s*_) −Δ*W* (*z*_*b*_). This includes the case of a shared maximum at the target interface.

For the atomic-union and finite-ion calculations, each thermal plane retains the part of the structural disk outside the union of all positive-radius spheres of the prepared charge-and-radius model (Fig. 2A, B). An ion-center clearance *a*_c_ is implemented by expanding each three-dimensional sphere radius by *a*_c_ before intersecting it with the plane. The five-point transverse kinetic stencil sets the wave function to zero at the first encountered boundary along an edge. It severs a link if a sphere intersects the edge between accessible nodes. We use the same boundary-distance cutoff convention as for the original disk. For classical integration we subtract the exact excluded intervals on each azimuthal ray and apply Gauss–Legendre quadrature in squared radius on each remaining interval. The transverse quantum trace uses the deterministic Chebyshev solver. Its local ratio to the classical lattice sum is calibrated by the infinite free square-lattice trace per node before applying the continuum classical integral. This calibration uses the same discrete kinetic operator for all irregular domains and preserves a comparison across meshes.

**Figure 2:**
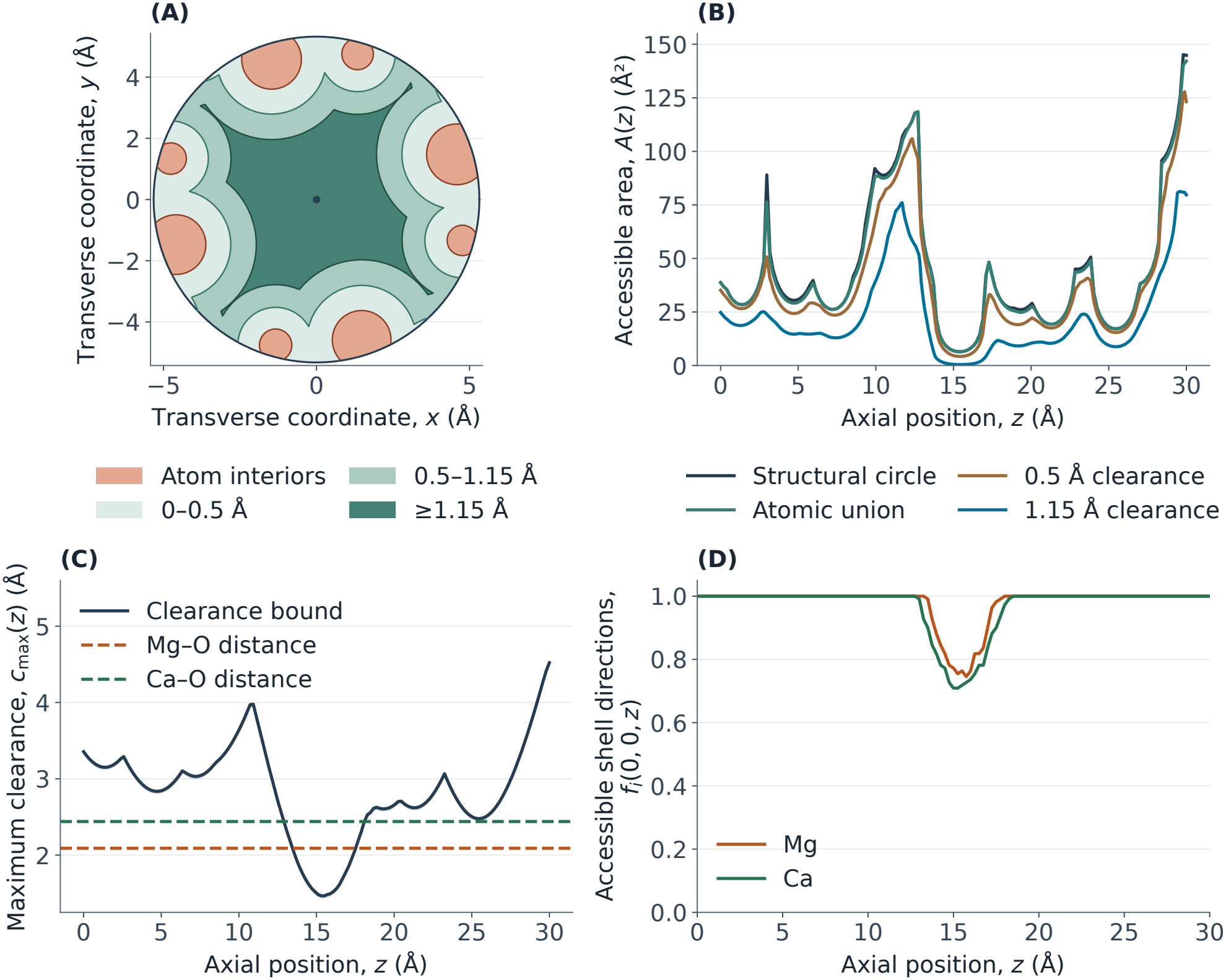
Atomic geometry and independent-direction shell access. (A) Transverse coordinates (*x, y*) at *z* = 2.980 Å, with clearance contours at 0, 0.5, and 1.15 Å. (B) Accessible area *A*(*z*) for each wall. (C) Upper and lower bounds on the maximal center clearance *c*_max_(*z*) and the representative ion–oxygen distances. (D) Fraction *f*_*i*_(0, 0, *z*) of unobstructed oxygen directions for each ion on the centerline. The masks and shell projection use positive-radius atomic spheres in three dimensions.

### 2.5 Diffusive passage to the membrane interfaces

The exit barrier measures the free-energy rise from the basin to the highest point along the route. The passage time also depends on the free energies elsewhere along the route. We model the axial motion as overdamped diffusion, in which the free-energy slope sets the drift and the diffusivity sets the time scale. Both the classical and quantum profiles enter the same diffusion model, which isolates the effect of the equilibrium correction on passage. Within this model, the process represents the exit step of single-ion permeation through the open channel.

We define the mean first-passage time (MFPT) as the average time to reach a chosen membrane interface for the first time. For either profile, it follows from the backward Smoluchowski equation with a common, spatially constant axial diffusivity *D* [37, 38],

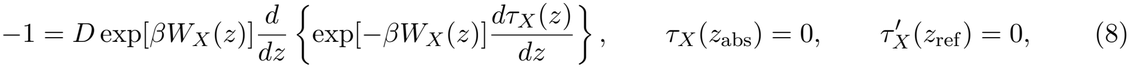

where *z*_abs_ denotes the absorbing interface, at which passage ends, and *z*_ref_ denotes the reflecting interface, which confines the ion at the opposite end. We evaluate the time from *z*_*b*_ in two separate boundary-value problems, one for each choice of absorbing interface. This convention yields directional escape times. A process that can terminate at either interface would use two absorbing boundaries. Integrating the equation gives two nested integrals over positions [37]. The outer position runs from *z*_*b*_ to the target interface and carries the factor exp(*βW*_*X*_). For each outer position, the inner position runs between it and the reflecting interface and carries the factor exp(−*βW*_*X*_). In this way the integral gives more weight to high free energies along the exit route and to low free energies in the regions accessible before passage. The common factor *D*^*−*1^ and any additive constant in *W*_*X*_ cancel from the ratio *R*_*τ*_ = *τ*_Q_*/τ*_C_. We report the relative change Δ*τ*_*s*_ = *R*_*τ,s*_ − 1 as a percentage.

We factored the exact passage integral to quantify the effects of the shape of each profile. For a profile *W*_*X*_, define an effective basin width 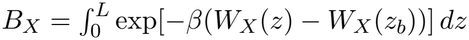 and an effective exit width 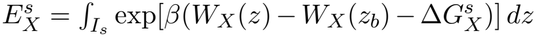. Their product times exp 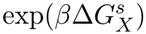 extends the inner integral to the full slab. The dimensionless factor 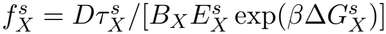 restores the nested limits, with 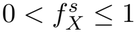 1. Thus

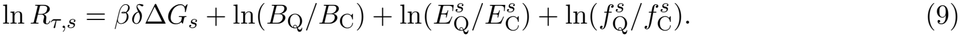

The reported directional times use trapezoidal quadrature on the axial thermal grid. The passage factorization and isotope comparisons integrate each segment of the piecewise-linear profiles analytically. The factorization uses the same integration rule for the time and both effective widths.

### 2.6 Origin of the quantum correction

We vary the wall geometry and the ion charge to identify which features of the free-energy landscape produce the quantum correction. Setting *U*_*i*_ = 0 isolates geometric confinement. Assigning charge +*e* to all three species makes the accessible space and electrostatic energy identical while their masses vary.

Magnesium and calcium already share the same energy at their physical charge of +2*e*. To test how access to positions near the wall affects the result, we smooth the radius near the entrance or along the entire pore, or reduce it by 0.5 Å at every axial position. All geometry tests use the original atomic coordinates and electrostatic field. The uniform circular inset prescribes the same displacement for each ion to test sensitivity to radial access. In a separate calculation we cut the actual union of the prepared atomic spheres out of each structural cross-section. We then expand the excluded spheres by a prescribed clearance for the finite-ion controls. The resulting domains retain their irregular transverse boundaries and any off-axis regions within the structural circle.

The Wigner–Kirkwood (WK) expansion relates thermal delocalization to the slope and curvature of the ion energy. For a smooth three-dimensional energy *U*_*i*_ away from the hard wall, its leading term gives the local free-energy correction [12, 13, 22]

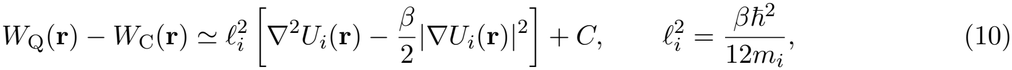

where *C* sets the free-energy reference. Here *W*_Q_(**r**) and *W*_C_(**r**) are local free energies derived from the three-dimensional position densities. The axial profiles include the transverse partition functions. At 310 K, the length *l*_*i*_ is 0.024, 0.023, and 0.018 Å for sodium, magnesium, and calcium, respectively. Both terms scale as 1*/m*_*i*_ at fixed energy, and the curvature and squared-gradient terms in energy units scale as *β* and *β*^2^, respectively. For the point-ion energy *U*_*i*_ = *Z*_*i*_*eϕ*, they scale as *Z*_*i*_ and 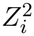, respectively, in the same field. The potential obeys Laplace’s equation wherever the permittivity is uniform and the region is free of fixed and mobile charges. In such a region the Laplacian vanishes, and the squared-gradient term contributes a nonpositive correction before a reference value is subtracted. The reduced axial profile depends on the pore radius and the energies away from the axis. It can therefore be sharply curved even where the potential on the axis is smooth.

We evaluate the one-dimensional Wigner–Kirkwood approximation from

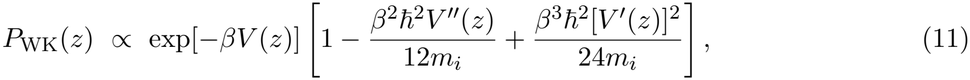

where 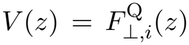 is the effective axial potential. We estimate its derivatives at interior axial positions from local quartic fits. Where the bracket is positive, we take the logarithm of the bracket, subtract its value at the basin position, and multiply the difference by −*k*_B_*T* . This diagnostic estimates the axial component in the smooth-potential regime. The transverse traces and axial thermal calculation give the total quantum correction and the changes in barriers and passage times. The Supplementary Materials describe the grid, sampling, and interpolation tests.

We estimated the smooth electrostatic contribution from the squared-gradient term of the local Wigner–Kirkwood expansion at centerline positions inside the two interfaces [12, 13, 22].

The smooth harmonic limit gives a separate energy scale for the basin. At the minimum of 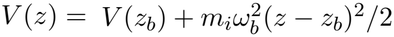, the leading local curvature term is 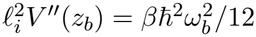. For ℏ*ω*_*b*_ = *k*_B_*T*, its value at 310 K is about 2.23 meV before normalization. The pore basin has changing transverse access and the normalized shift also includes the reference, so we use the integrated basin-width contribution in Eq. (9) to interpret the passage time.

### 2.7 Screened pair and isotope calculations

To estimate how a second ion of the same species affects escape, we combine two copies of the classical transverse free-energy profile with a screened Coulomb repulsion 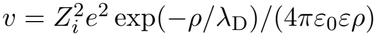. The effective separation is *ρ* = [(*z*_1_ − *z*_2_ )^2^ + *a* ^2^]^1*/*2^, with transverse offset *a* = 2 Å and screening length *λ*_D_ = 8.01 Å. We evaluate this prescribed interaction at 310 K with relative permittivities *ε* = 20, 40, and 78.5. The screening length is the bulk Debye value for 150 mM electrolyte at *ε* = 78.5 and remains fixed across the three permittivities. This comparison varies the strength of the prescribed pair repulsion at a common screening range. The classical partition function for two indistinguishable ions is 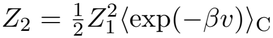. Here *Z*_1_ is the single-ion partition function, and the average uses two independent draws from the classical single-ion Boltzmann distribution. The interaction raises the pair free energy by Δ*F*_pair_ = −*k*_B_*T* ln ⟨exp(− *βv*) ⟩ _C_ relative to the same two-ion system with *v* = 0. Integrating over one ion’s position gives the marginal free-energy profile whose barriers describe escape for its partner. The reported pair barriers use the local marginal minimum within the Y1085–E1047 reference window, consistent with the circular single-ion comparison.

This benchmark conditions on the presence of two ions. The equilibrium odds of double versus single occupancy in a one-species grand-canonical model are *P*_2_*/P*_1_ = exp(*βµ*)*Z*_2_*/Z*_1_, with chemical potential *µ* and a common energy reference for pore and bath [7]. The pair free-energy increment characterizes repulsion within the occupied pore. The occupancy odds also depend on the free energy of inserting an ion from the bath.

We repeated the two-ion benchmark on the directly calculated classical transverse profiles. Both ions follow independent axial imaginary-time Brownian bridges in the kinetic reference; their potential along a path is the sum of the two single-ion profiles and the screened repulsion used in the classical two-ion benchmark. For each thermal trace, we drew the path roots from the corresponding classical Boltzmann distribution on a midpoint grid. The quantum-to-classical partition-function ratio, after removal of the free kinetic prefactor, is the classical-root average of the bridge weight exp[− *β*(*Ū* −*U*_root_)], where *Ū* is the discretized imaginary-time average and paths with an axial bead outside the slab receive zero weight [22]. In the ratio of the interacting trace to the square of the single-ion trace, the kinetic prefactors cancel. The bridge integrations, time-slicing comparisons, and root-grid settings are given in the Supplementary Materials.

We sampled the electrostatic cost of adding a divalent charge at four axial positions, using the same directional permittivity and mobile-ion accessibility maps as in the matched-grid calculations. Each probe carried +2*e* in a spherical Gaussian with standard deviation 0.38 Å, truncated at three standard deviations and normalized on the mesh. All four source supports cleared the prepared atomic spheres. We used the linear response of the nonlinear Poisson–Boltzmann equation about the parent potential *u*_0_. The operator retains the dielectric term and a mobile-ion response coefficient proportional to *χ*(**r**) cosh[*u*_0_(**r**)], where *χ* is the dimensionless mobile-ion accessibility and *u*_0_ = *eϕ*_0_*/*(*k*_B_*T*_PB_) is the background potential in thermal units. Owing to the linear response, the screened pair energy is the probe charge times the responding potential averaged over the second probe. Its value is symmetric under exchange of the two source locations. To isolate the dielectric self-energy, we also solved without mobile-ion response in the heterogeneous dielectric and in a homogeneous water reference. One half of the source-weighted potential difference gives the self-energy relative to water.

The four-site calculation provides a cross-check for a larger axial map. We extended the same calculation to 16 on-axis sites from *z* = 0 to 30 Å. At each site, we applied a normalized Gaussian source in turn and averaged the induced potential over all 16 test probes with the same charge shape. With *δϕ*_*i*_ the response to probe *i* and *ρ*_*j*_ the charge density of probe *j*, the screened pair energy is *v*_*ij*_ = ∫*ρ*_*j*_*δϕ*_*i*_ *d*^3^**r**. The factor of one half applies to the diagonal charging energy relative to water. For a pair of different probes, the two equal cross terms remove this factor. The complete response matrices include the screened, dielectric-only, and homogeneous-water calculations. The same grid and boundary conditions recover the four-site values within 10^*−*7^ meV.

We recalculated the axial thermal density at both isotope masses on each common classical transverse free-energy profile. The magnesium and calcium pairs used their respective divalent profiles. The lithium pair used the sodium profile as a fixed monovalent test-charge landscape. This control isolates the effect of isotope mass on axial motion at fixed transverse free energy. We used tabulated isotope masses [17]. The axial-spacing comparison is given in the Supplementary Materials.

### 2.8 Pore fluctuations and coupled thermal motion

The fluctuation and coupled-plane calculations use the point-ion reference field. We denote the root-mean-square amplitude of the prescribed deformation mode by *σ*_def_.

We test an ensemble of smooth axial deformations that advect both the wall and the static electrostatic landscape. Within an interval of width *b*_def_, the deformation is the flow generated by sin^2^(*πz/b*_def_) with a zero-mean Gaussian amplitude. It fixes the endpoints and approaches unit slope there. We use *b*_def_ = 6.1 Å for the entrance mode and *b*_def_ = 30 Å for the whole-pore mode. Inverse mapping of each sampled conformation transfers both transverse partition functions to the deformed laboratory coordinates. This is a prescribed open-state ensemble of correlated axial displacements. Local side-chain reorientation, water exchange, and dielectric reorganization would require a structural dynamics model.

We compare two statistical limits. In the fast-exchange approximation we average the unnormalized position weights with the bare conformer prior before obtaining the free-energy profile and passage estimate. In the slow-exchange approximation we average the passage times over conformers before forming the quantum-to-classical ratio.

The plane correction Δ*W* ^plane^ is the quantum-minus-classical free-energy difference for the chosen diametric plane.

We calculated the axial heat-trace profile of a two-dimensional diametric plane through the steep entrance. The transverse direction was selected along a favorable off-axis position in the slice at 2.980 Å. We also tested its perpendicular plane. The calculation uses the three-dimensional molecular field restricted to that plane. A two-dimensional Cartesian Dirichlet lattice retains kinetic links in both directions wherever adjacent nodes are accessible. A Chebyshev expansion evaluates the diagonal of the coupled heat operator and groups its entries by axial plane. The comparison uses the same nodes and potential for two reductions. One uses a scalar transverse free energy in the axial operator. The other diagonalizes the transverse problem at every plane and propagates the ordered transverse bands independently with axial Dirichlet endpoints. This supplies a mode-resolved reference that omits changes of transverse eigenfunctions in the axial kinetic operator. On a separable test potential all three calculations recover the same heat-trace profile as dense diagonalization.

We solved the coupled problem on a local entrance domain and examined its interior. The plane quantum corrections use a common axial reference within each comparison.

## 3 Results

### 3.1 Water coordination, insertion and occupied wells

At the narrowest tested plane, the maximal ion-center clearance is bounded across the structural disk (Fig. 2C). This clearance lies below both representative bulk ion–oxygen distances. A spherical envelope that extends to the first-shell oxygen radius therefore has no continuous path through the full modeled slab for either divalent ion. A discrete shell can rotate and retain a subset of its waters as it enters a constriction (Fig. 2D).

At the centerline of the narrowest structural section, the primary water probe admits only part of the first-shell coordination geometry. The correlated shell has a free-energy cost there under the primary penalty (Fig. 3A). Average site counts use the sharp distance criterion, and the energy calculation uses the smooth access factor. Both refined orientation samples give zero fully accessible orientations at this site for either geometry. This observation concerns the centerline and the specified coordination geometries.

**Figure 3:**
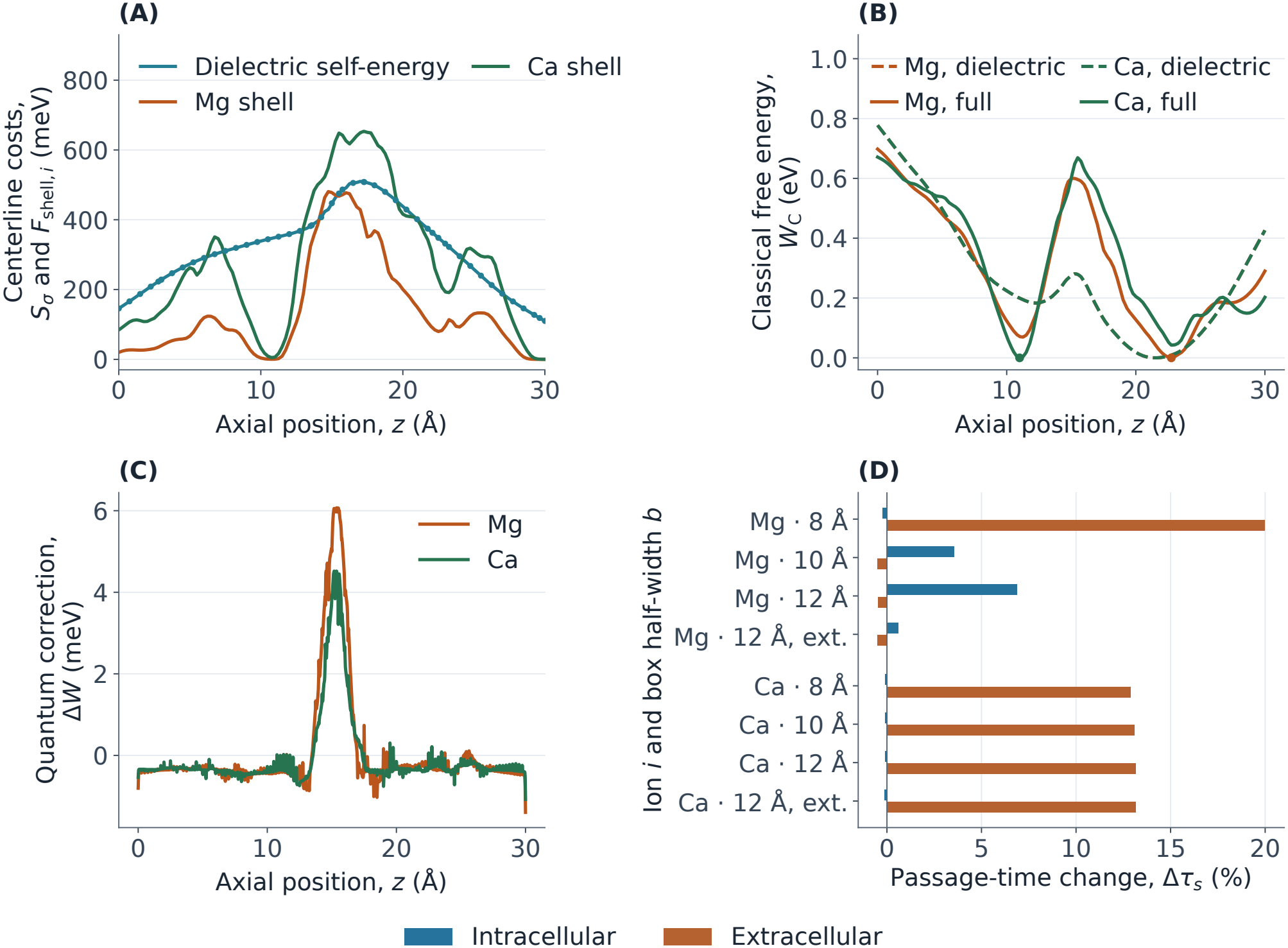
Finite-ion insertion and escape at 310 K. (A) Centerline dielectric self-energy *S*_*σ*_ and correlated hydration costs *F*_shell,*i*_ along *z* for ion species *i*. Markers are direct axial probes. (B) Classical free energies *W*_C_(*z*) with the dielectric term alone and with both insertion terms, each relative to its global well. (C) Full-model quantum correction Δ*W* (*z*) referenced to the same well as the classical profile. Panels A–C use the extended axial box; C shows the quantum correction on the finest transverse grid. (D) Directional percentage changes Δ*τ*_*s*_ by ion and lateral box half-width *b*, at 0.085 Å transverse spacing. The extended box moves the axial faces from −4 and 36 Å to −7.5 and 37.5 Å.

At the point-ion magnesium basin plane, the refined probe benchmark resolves the charging energy on the pore axis and at an interior off-axis position. The largest spatial-calculation box gives a corresponding centerline estimate at the same plane. These charging energies exceed the quantum barrier shifts of the bare magnesium reference. Their spatial variation changes which positions contribute to the transverse Boltzmann weights.

The primary calculation uses the extended axial dielectric domain. Magnesium and calcium occupy different deepest classical wells, with distinct intracellular and extracellular barriers (Fig. 3B; Tables 1 and 2). From these wells, quantization lengthens intracellular passage and shortens extracellular passage for magnesium. For calcium it shortens intracellular passage and lengthens extracellular passage. The extracellular route from the calcium well crosses the constriction, where the quantum free-energy profile raises the exit barrier. The preferred magnesium well lies beyond the constriction on the extracellular side (Fig. 3C).

**Table 1.** Magnesium escape from the global classical basin at 310 K. The Gaussian background and minimum ion-center clearance are common to these cases. Box widths are lateral half-widths. Ligand penalties and oxygen-probe variations use the extended axial box. The −and + directions lead to the intracellular and extracellular interfaces.

| Model | *h*<sub>⊥</sub> (Å) | *z*<sub>*b*</sub> (Å) | Δ*G*<sup>*s*</sup><sub>C</sub> (meV) |  | Δ*τ*<sub>*s*</sub> (%) |  |
|---|---|---|---|---|---|---|
|  |  |  | − | + | − | + |
| Gaussian background | 0.085 | 19.79 | 1070 | 743 | −6.05 | −6.95 |
| Correlated shell only | 0.085 | 22.08 | 901 | 523 | −0.65 | −0.98 |
| Self-energy, extended axial box | 0.085 | 21.46 | 778 | 426 | −0.61 | −1.40 |
| Independent ligands, extended axial box | 0.085 | 22.08 | 711 | 321 | −0.40 | −0.68 |
| Correlated ligands, 8 Å box | 0.085 | 11.25 | 670 | 562 | −0.23 | 19.99 |
| Correlated ligands, 10 Å box | 0.085 | 22.75 | 638 | 290 | 3.54 | −0.48 |
| Correlated ligands, 12 Å box | 0.085 | 22.75 | 630 | 282 | 6.87 | −0.48 |
| Correlated ligands, extended axial box | 0.085 | 22.75 | 698 | 290 | 0.61 | −0.49 |
| Correlated ligands, extended axial box | 0.065 | 22.75 | 698 | 290 | 0.69 | −0.58 |
| Correlated ligands, extended axial box | 0.050 | 22.75 | 698 | 290 | 0.68 | −0.66 |
| Refined shell, extended axial box | 0.085 | 22.75 | 697 | 290 | 0.59 | −0.48 |
| Finer axial sampling, 8 Å box | 0.065 | 11.25 | 670 | 562 | −0.23 | 22.65 |
| Ligand penalty 2*k*<sub>B</sub>*T* | 0.085 | 22.50 | 711 | 315 | −0.35 | −0.58 |
| Ligand penalty 6*k*<sub>B</sub>*T* | 0.085 | 22.75 | 801 | 287 | 21.47 | −0.53 |
| Oxygen probe 0.70 Å | 0.085 | 22.50 | 759 | 390 | −0.28 | −0.54 |

**Table 2.**
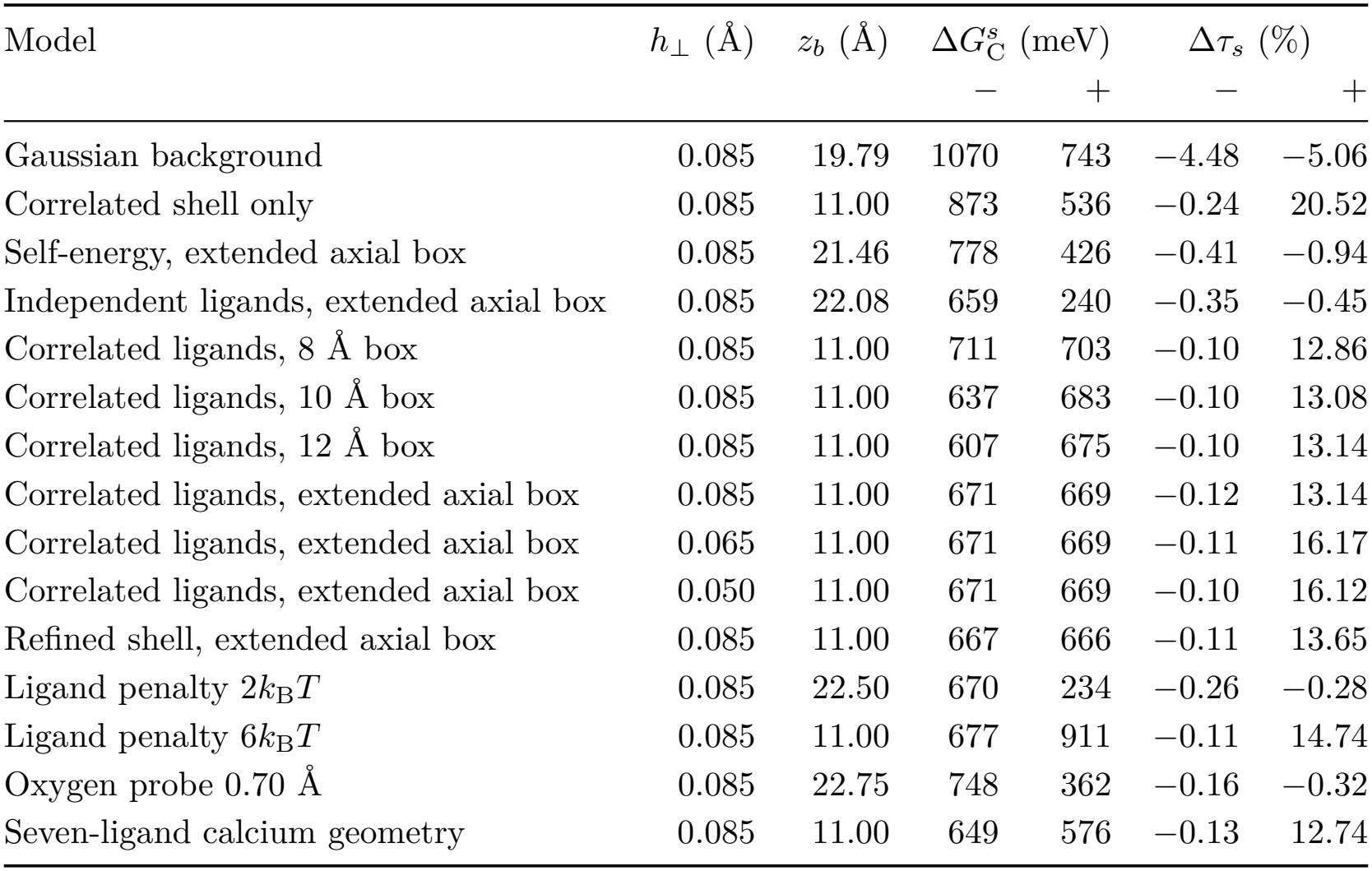
Calcium escape from the global classical basin at 310 K. The Gaussian background and minimum ion-center clearance are common to these cases. Box widths are lateral half-widths. Ligand penalties and oxygen-probe variations use the extended axial box. The −and + directions lead to the intracellular and extracellular interfaces.

The deepest magnesium well in the smaller lateral box lies on the intracellular side of the constriction. With greater lateral extent it lies on the extracellular side. At common transverse spacing, extending the axial domain also changes the directional passage effects for magnesium and calcium (Fig. 3D).

In the extended box, the matched independent-ligand calcium control favors a downstream well and shortens extracellular passage. A common shell rotation places the well upstream and increases the extracellular passage time. The comparison uses the same dielectric field, ligand count, access criterion, and obstruction penalty. Within the correlated model, reducing the ligand penalty moves the calcium well downstream and changes the extracellular response to a decrease (Table 2).

To resolve regional retention, we also start passage from local classical wells in the same insertion landscape. The Y1085–E1047 window minima define companion starting positions and energy gaps relative to the global minima. Passage from the window minimum gives the corresponding local-well comparison for each ion and exit direction. The upstream magnesium well supplies a further extracellular escape estimate. The positions, well-depth differences, and passage responses are retained in the Supplementary Materials.

We also averaged the two pointwise passage profiles over a shared classical equilibrium starting distribution. For magnesium, the upstream part of the pore carries a small fraction of the classical weight and a much larger share of the averaged classical extracellular time. The long extracellular escape time from the upstream side accounts for its large contribution to the average.

For intracellular passage of magnesium, we decompose the logarithm of the time ratio into the barrier change, basin weight, exit width, and nested integration limits (Eq. 9). The positive contributions from the basin and exit weights exceed the contribution of the barrier decrease and leave the overall time ratio above one.

### 3.2 Atomic boundaries and dielectric sensitivity

Independent off-axis sites with the guarded clearance and sites closer to the atomic spheres were withheld from calibration. The largest discrepancies occurred at low-clearance sites (Fig. 4A, B). At the basin plane, the interior and nearby wall probes give distinct charging energies. Part of the wall probe’s Gaussian charge can overlap the atomic spheres. These comparisons quantify the loss of accuracy as the Gaussian approaches the protein boundary. The guarded control restricts centers to the prescribed clearance. Its electrostatic uncertainties still exceed individual quantum free-energy changes at some sites.

**Figure 4:**
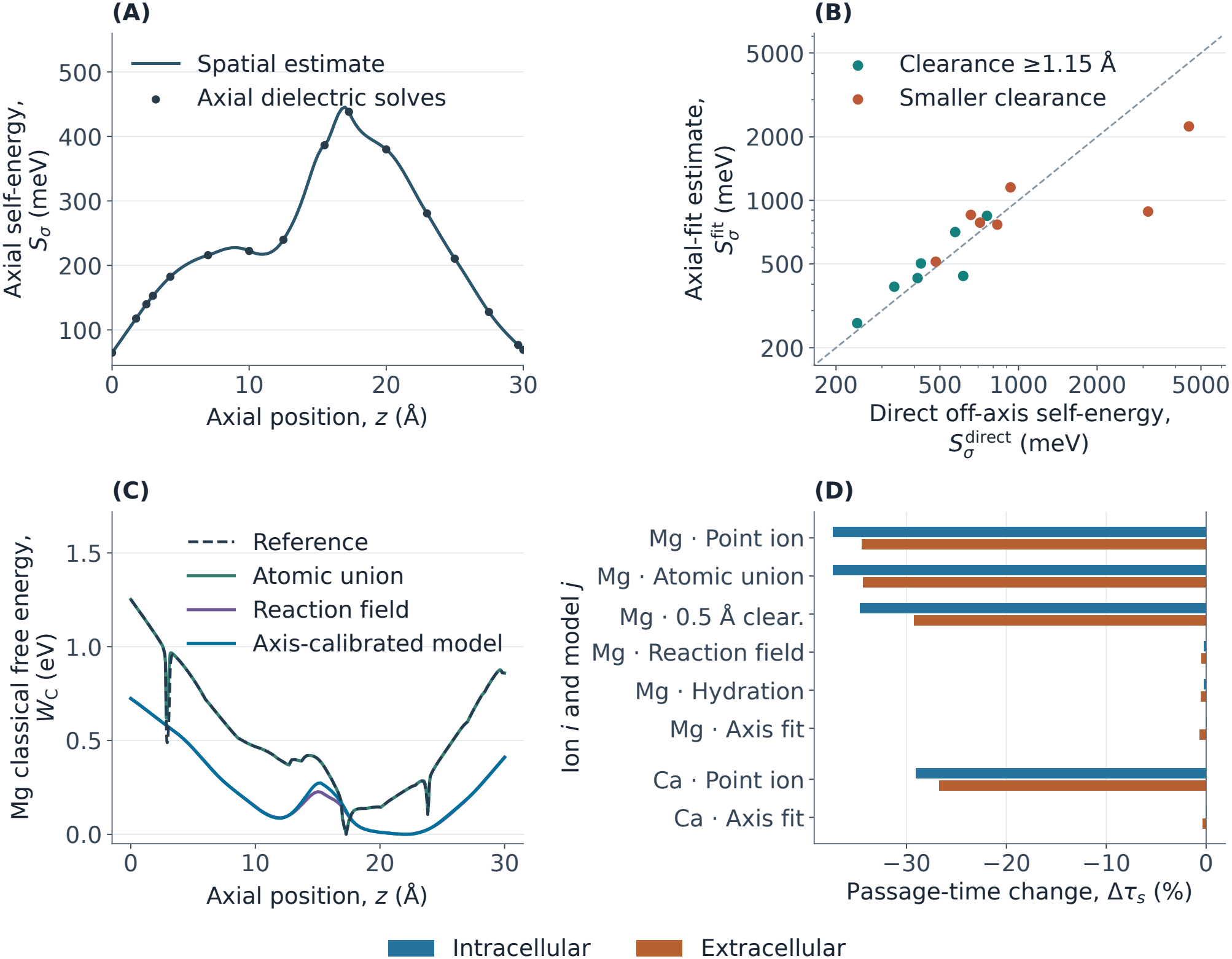
Axis-calibrated dielectric and hydration controls at 310 K. (A) Axial self-energy *S*_*σ*_(*z*) and its direct Gaussian-probe values. (B) Axis-calibrated estimates 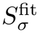 and direct values 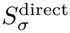 at held-out off-axis sites. The diagonal indicates equality. At smaller clearance, some of the Gaussian charge can overlap the spheres. (C) Classical magnesium free energy *W*_C_(*z*) relative to each model’s basin. (D) Directional passage-time percentage change Δ*τ*_*s*_ at common diffusivity, indexed by ion *i* and model *j*. The axis fit uses 1.15 Å atomic clearance, the Gaussian background, and the independent-direction shell rule in Eq. 4.

Cutting out the atomic spheres reduces the magnitude of the local magnesium peak. Passage times remain shorter toward both interfaces. A clearance around the spheres gives a different perturbation from a uniform inset of the circular wall. Adding the reaction field at fixed atomic clearance moves the classical magnesium basin and reduces the classical intracellular barrier. The independent-shell calcium calculation supplies the corresponding comparison at the guarded clearance (Fig. 4C, D).

The axis-calibrated estimate raises low-energy positions in the original basin and alters the classical barriers. We compare several controls, including the variational energies propagated independently, the Gaussian background average, and the shell-access penalty (Table 3). The lower and upper variational fields give different changes, and the intracellular change can have either sign. The two trial fields give model sensitivity cases for passage because the passage integral depends on energy differences along the entire profile. In the upper-energy case, an additional classical well lies below the minimum in the reference basin window. Starting at this global minimum gives a companion directional-passage comparison. The axis-calibrated comparisons place the global classical minima inside the reference basin window. The passage comparison depends on spatial differences in energy, so the bounds on charging energy define sensitivity cases for escape.

**Table 3.**
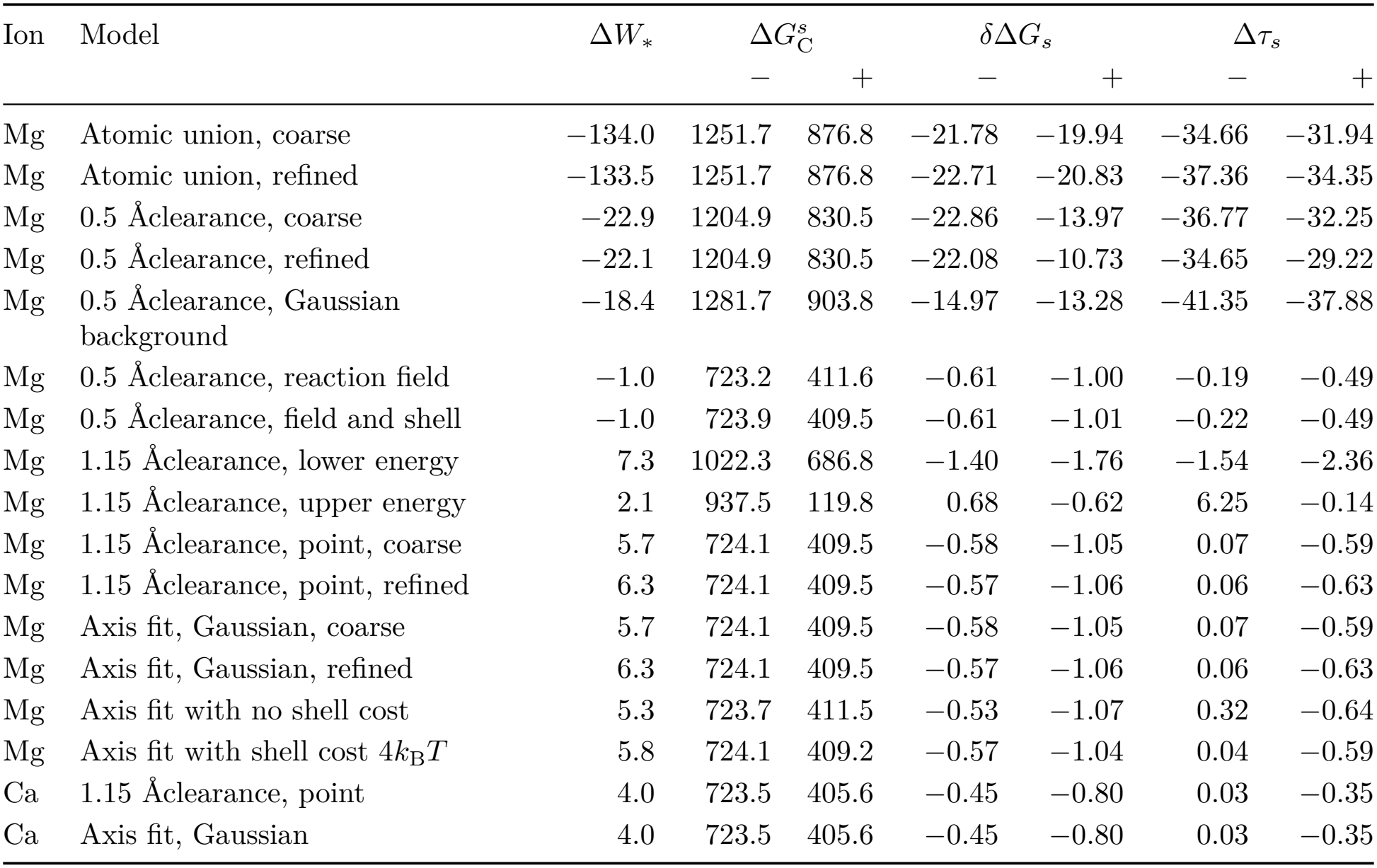
Atomic-union and axis-calibrated sensitivity controls at 310 K. Each row starts at the classical minimum within the Y1085–E1047 window. The extremum Δ*W*_*\**_ is the largest quantum correction by magnitude, with its sign retained and the classical basin as reference. The classical barriers and their quantum changes are in meV, with −denoting the intracellular direction and + the extracellular direction. Passage changes are in percent. Rows marked refined use 0.065 Å transverse spacing; all other rows use 0.085 Å. The independent-direction shell rule is Eq. 4.

| Ion | Model | ΔW<sub>*</sub> | ΔG<sup>s</sup><sub>C</sub> |  | δΔG<sub>s</sub> |  | Δτ<sub>s</sub> |  |
|---|---|---|---|---|---|---|---|---|
|  |  |  | − | + | − | + | − | + |
| Mg | Atomic union, coarse | −134.0 | 1251.7 | 876.8 | −21.78 | −19.94 | −34.66 | −31.94 |
| Mg | Atomic union, refined | −133.5 | 1251.7 | 876.8 | −22.71 | −20.83 | −37.36 | −34.35 |
| Mg | 0.5 Åclearance, coarse | −22.9 | 1204.9 | 830.5 | −22.86 | −13.97 | −36.77 | −32.25 |
| Mg | 0.5 Åclearance, refined | −22.1 | 1204.9 | 830.5 | −22.08 | −10.73 | −34.65 | −29.22 |
| Mg | 0.5 Åclearance, Gaussian background | −18.4 | 1281.7 | 903.8 | −14.97 | −13.28 | −41.35 | −37.88 |
| Mg | 0.5 Åclearance, reaction field | −1.0 | 723.2 | 411.6 | −0.61 | −1.00 | −0.19 | −0.49 |
| Mg | 0.5 Åclearance, field and shell | −1.0 | 723.9 | 409.5 | −0.61 | −1.01 | −0.22 | −0.49 |
| Mg | 1.15 Åclearance, lower energy | 7.3 | 1022.3 | 686.8 | −1.40 | −1.76 | −1.54 | −2.36 |
| Mg | 1.15 Åclearance, upper energy | 2.1 | 937.5 | 119.8 | 0.68 | −0.62 | 6.25 | −0.14 |
| Mg | 1.15 Åclearance, point, coarse | 5.7 | 724.1 | 409.5 | −0.58 | −1.05 | 0.07 | −0.59 |
| Mg | 1.15 Åclearance, point, refined | 6.3 | 724.1 | 409.5 | −0.57 | −1.06 | 0.06 | −0.63 |
| Mg | Axis fit, Gaussian, coarse | 5.7 | 724.1 | 409.5 | −0.58 | −1.05 | 0.07 | −0.59 |
| Mg | Axis fit, Gaussian, refined | 6.3 | 724.1 | 409.5 | −0.57 | −1.06 | 0.06 | −0.63 |
| Mg | Axis fit with no shell cost | 5.3 | 723.7 | 411.5 | −0.53 | −1.07 | 0.32 | −0.64 |
| Mg | Axis fit with shell cost 4*k*<sub>B</sub>*T* | 5.8 | 724.1 | 409.2 | −0.57 | −1.04 | 0.04 | −0.59 |
| Ca | 1.15 Åclearance, point | 4.0 | 723.5 | 405.6 | −0.45 | −0.80 | 0.03 | −0.35 |
| Ca | Axis fit, Gaussian | 4.0 | 723.5 | 405.6 | −0.45 | −0.80 | 0.03 | −0.35 |

Each full spatial field was propagated through the thermal and passage calculations. These interpolation comparisons share the dielectric boundary within each box. The independent comparisons across boxes measure the response to changing that boundary.

Varying the ligand penalty in the extended axial box changes the sign of the magnesium intracellular response and the calcium extracellular response. At the reference penalty, reducing the water probe moves the lowest calcium well downstream and shortens extracellular passage. The seven-ligand geometry with the primary water probe retains the intracellular-side calcium well and lengthens extracellular passage (Tables 1 and 2).

### 3.3 Point-ion reference and regional escape

A widening gives the ion a larger area and can expose new electrostatic environments. Both effects change the transverse partition function, so the preferred axial position depends on the whole cross-section. The classical free-energy minimum lies on the intracellular side of the centerline potential minimum. The positions differ because the transverse Boltzmann integral depends on the area of each slice and the mean Boltzmann factor over that area.

From the interior minimum, an ion encounters a free-energy rise on the way to either membrane interface. The highest point along the intracellular route lies at the interface for all three ions. On the extracellular route it lies at the interface for sodium and inside the slab near the extracellular interface for magnesium and calcium. The classical and quantum profiles place the interior minimum and exit maxima in the same regions of the membrane slab (Fig. 5A). These are confined escape barriers of the fixed test-charge model. A bulk-to-bulk permeation profile would include the energy of ion insertion, the vestibules, and occupancy by mobile ions [8, 30, 31]. The barrier heights represent a large thermal penalty for an ion to leave its preferred site in the bare-charge model. Dielectric charging and water coordination change the landscape on which that retention occurs.

**Figure 5:**
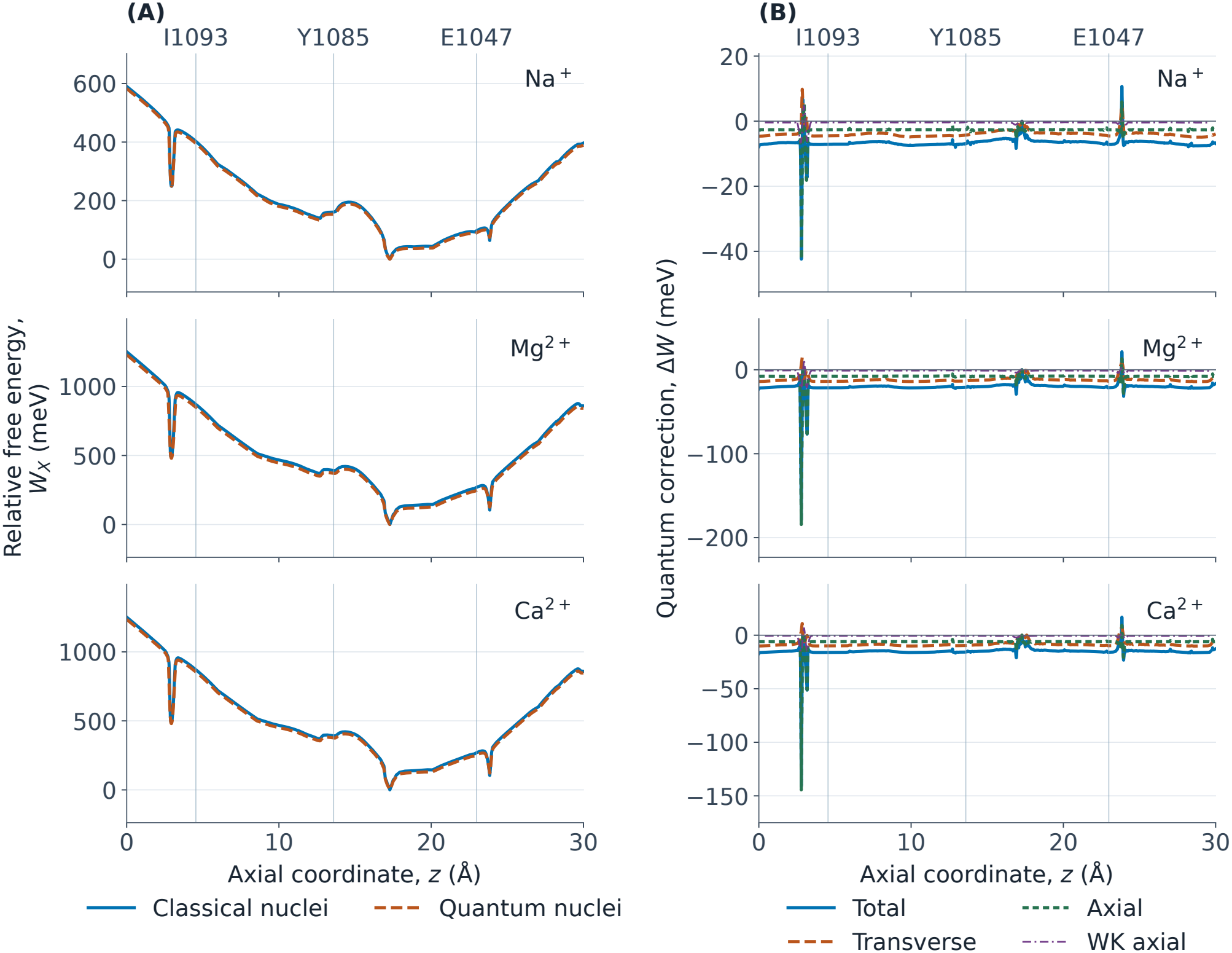
Circular point-ion reference at 310 K, with sodium, magnesium, and calcium from top to bottom. (A) Classical and quantum free energies *W*_C_(*z*) and *W*_Q_(*z*) as functions of axial position *z*, referenced to the classical basin. (B) The total correction Δ*W* = Δ*F*_*⊥*_ + Δ*W*_*z*_, its transverse and axial terms, and the Wigner–Kirkwood estimate of Δ*W*_*z*_. All corrections use the same basin reference. Vertical lines mark I1093, Y1085, and E1047.

The local effect of quantization is greatest near the intracellular entrance, where the free energy changes over a short distance. For magnesium, motion along the pore lowers the local free energy, and motion across the pore offsets part of that change. Calcium likewise gives a negative axial contribution and a positive transverse contribution. For sodium, the axial contribution is negative and the transverse contribution is small. At that entrance peak, the smooth-potential estimate captures a smaller axial response for all three ions (Eq. 11, Fig. 5B). For magnesium, the negative entrance extremum is followed by a positive local maximum. Smaller variations occur farther down the pore. Smoothing the radius near the entrance moves the largest magnesium correction outside the entrance region. The sharp widening therefore controls where the largest local correction occurs, while the slow steps of escape lie elsewhere in the pore (Supplementary Materials).

The entrance widening exposes substantially lower-energy positions across neighboring slices. Across the widening, the radius increases and the classical magnesium free energy falls sharply. The increase in area alone accounts for a smaller part of the decrease. Favorable positions close to the protein account for most of the lowering. Delocalization across a sharp change in the positions accessible to the ion explains the much larger axial correction.

The centerline squared-gradient estimates are small compared with the entrance axial correction. This is a gradient-term diagnostic on the pore axis. The effective axial potential also depends on the area and energy distribution across the section, including regions where the permittivity changes. The centerline diagnostic therefore quantifies a different contribution from the axial Wigner–Kirkwood estimate (Eq. 11). A scale estimate illustrates the difference. In the leading local expression for magnesium, an entrance-sized correction would require a much steeper energy slope or curvature than the centerline diagnostic resolves. These are one-term scales calculated by inserting the size of the entrance axial correction into the smooth expansion, before choosing a reference. The axial correction itself is basin-referenced and resolves an abrupt change in accessible configurations. The one-term estimates give a scale for comparison with a smooth landscape.

In the circular point-ion reference, quantum delocalization shortens the modeled time to reach either membrane interface. Magnesium shows the largest effect on passage (Fig. 6A). Its exit-barrier changes are much smaller than the largest local free-energy change at the entrance (Fig. 6B, Table 4). Along the intracellular route, the free-energy difference derived from the normalized densities, −*k*_B_*T* ln(*P*_Q_*/P*_C_), is positive at the basin minimum and negative at the barrier maximum. The change in the barrier height is the difference between these two values.

**Table 4.**
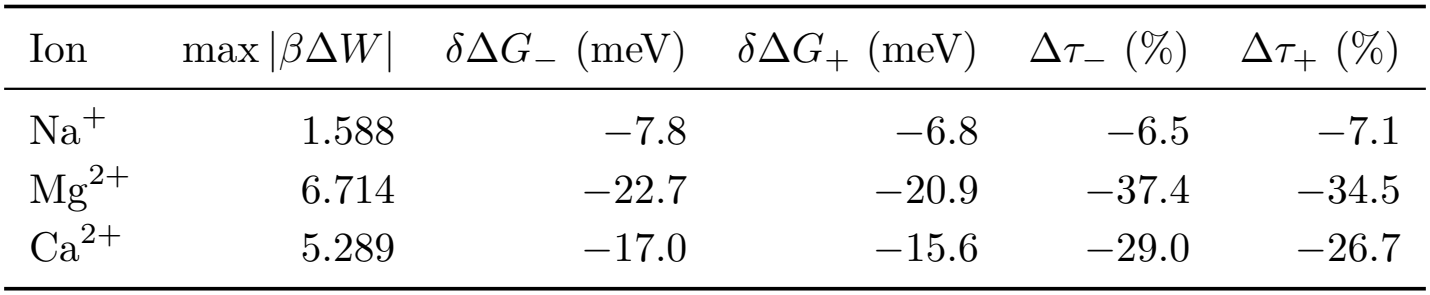
Circular point-ion reference at 310 K: largest local free-energy correction, changes in exit barrier heights, and changes in mean first-passage times. The local peak max |*β*Δ*W*| is an absolute magnitude in thermal units. The barrier corrections and relative time changes are signed. Negative values indicate a reduction from the classical result. The − and + directions denote passage to the intracellular (*z* = 0) and extracellular (*z* = *L*) interfaces, respectively.

| Ion | max \|βΔW\| | δΔG<sub>−</sub> (meV) | δΔG<sub>+</sub> (meV) | Δτ<sub>−</sub> (%) | Δτ<sub>+</sub> (%) |
|---|---|---|---|---|---|
| Na<sup>+</sup> | 1.588 | −7.8 | −6.8 | −6.5 | −7.1 |
| Mg<sup>2+</sup> | 6.714 | −22.7 | −20.9 | −37.4 | −34.5 |
| Ca<sup>2+</sup> | 5.289 | −17.0 | −15.6 | −29.0 | −26.7 |

**Figure 6:**
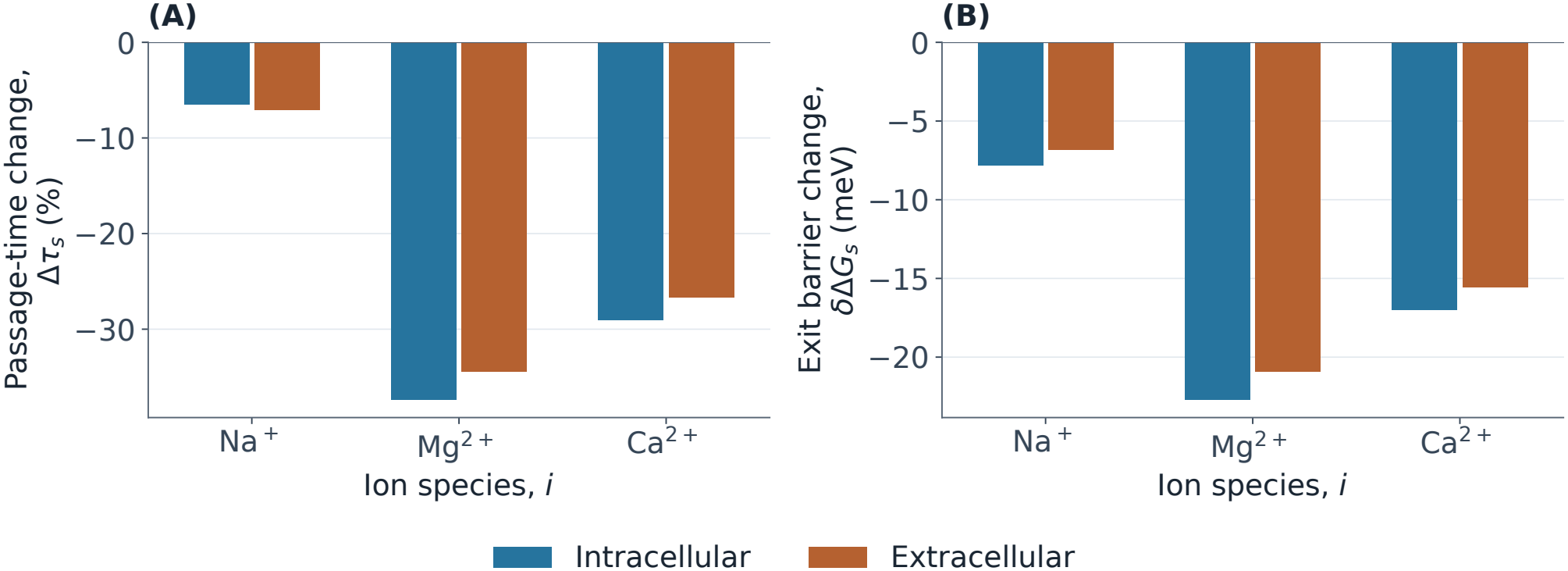
Quantum changes in directional escape for ion species *i* in the circular point-ion reference at 310 K. (A) Relative mean first-passage time change Δ*τ*_*s*_, expressed as a percentage. (B) Change in exit barrier *δ*Δ*G*_*s*_. At each classical starting well the direction *s* selects the intracellular or extracellular interface. The barrier is measured to the maximum on each profile.

The magnesium barrier decreases exceed the calcium decreases for both exit routes. Their ratio is below the inverse-mass ratio *m*_Ca_*/m*_Mg_. The barrier correction combines free-energy changes at positions where the local curvature and confinement differ. The passage time also depends on the widths of these regions because the integrals include positions near the basin minimum and barrier maximum [10, 37]. For magnesium passage to the intracellular interface, the logarithm of the time ratio, ln *R*_*τ, −*_, differs from the barrier change in thermal units, *βδ*Δ*G*_*−*_ . Factoring the passage integral (Eq. 9) separates the basin-width, exit-width, and nesting contributions. The basin and exit-width contributions are positive, and the nesting contribution is small in either direction. The buffer and reflecting-boundary comparisons appear in the Supplementary Materials. Most of the gap reflects the broader thermally occupied basin in the quantum profile.

At the wider entrance slice, a small part of the reference disk overlaps the prepared atomic spheres and carries almost all of the classical magnesium weight. Excluding the points inside these spheres raises the classical slice free energy (Supplementary Materials). The magnesium extremum lies on the preceding, narrower side of the widening. Its sensitivity to the wall follows from the change in the accessible positions of adjacent slices.

The sphere-excluding circle reduces the accessible radius (Fig. 7A). We use the same potential, temperature, ion mass, thermal grids, and passage boundaries as in the structural circle. The largest local correction is reduced in magnitude and remains near the intracellular widening (Fig. 7B). In this geometry, the quantum barriers and passage times decrease toward both interfaces (Fig. 7C, D). The classical intracellular passage time also decreases at the same diffusivity. The local peak and the relative changes in escape are therefore sensitive to different parts of the landscape. The centered disk tests sphere exclusion within the circular model. The irregular-wall comparisons retain off-axis space and resolve the effects of geometry, hydration, and dielectric charging (Fig. 4).

**Figure 7:**
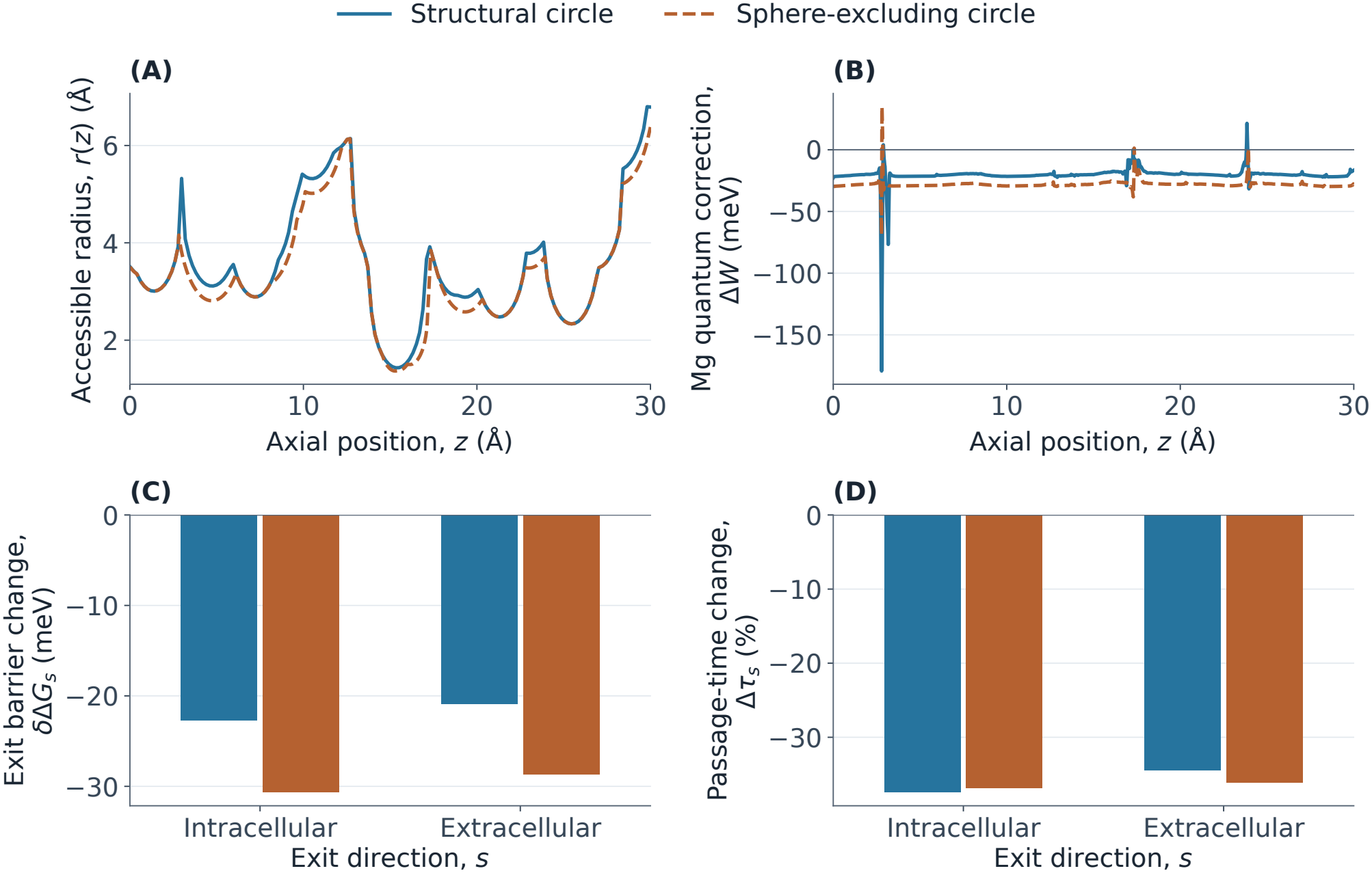
Effect of circular exclusion of prepared atomic spheres on magnesium at 310 K. (A) The structural and sphere-excluding pore radii *r*(*z*) and (B) the corresponding magnesium correction Δ*W* (*z*) along *z*. (C) Quantum exit-barrier shift *δ*Δ*G*_*s*_ and (D) relative mean first-passage time change Δ*τ*_*s*_ upon quantization in each circular geometry. The direction *s* selects the intracellular or extracellular interface.

The uniform 0.5 Å inset also restricts access in the occupied well. In the structural circle, the outer ring carries most of the classical magnesium weight at the entrance peak and at the basin. The inset attenuates the passage-time response toward both interfaces (Supplementary Materials). Together the two wall comparisons show how access near the occupied site shapes the escape response. The circular inset is a geometry control for the point ion. The finite-ion model also accounts for dielectric charging and restricted water coordination.

The charge controls help explain why the local effect can be large even when the thermal wavelength is short relative to the pore diameter. With the electrostatic energy set to zero, the maximum free-energy correction in thermal units, max |*β*Δ*W*|, is small for all three ions. The leading boundary term in the Dirichlet heat trace, the thermal sum over states whose wave functions vanish at the wall, estimates the local transverse confinement cost as Δ*F*_*⊥*_ */k*_B_*T*≃ *λ*_th,*i*_*/*(2*r*) [14]. At fixed radius this estimate scales as 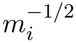. The total geometry-only correction also includes axial motion and is measured relative to the basin (Fig. 8A).

**Figure 8:**
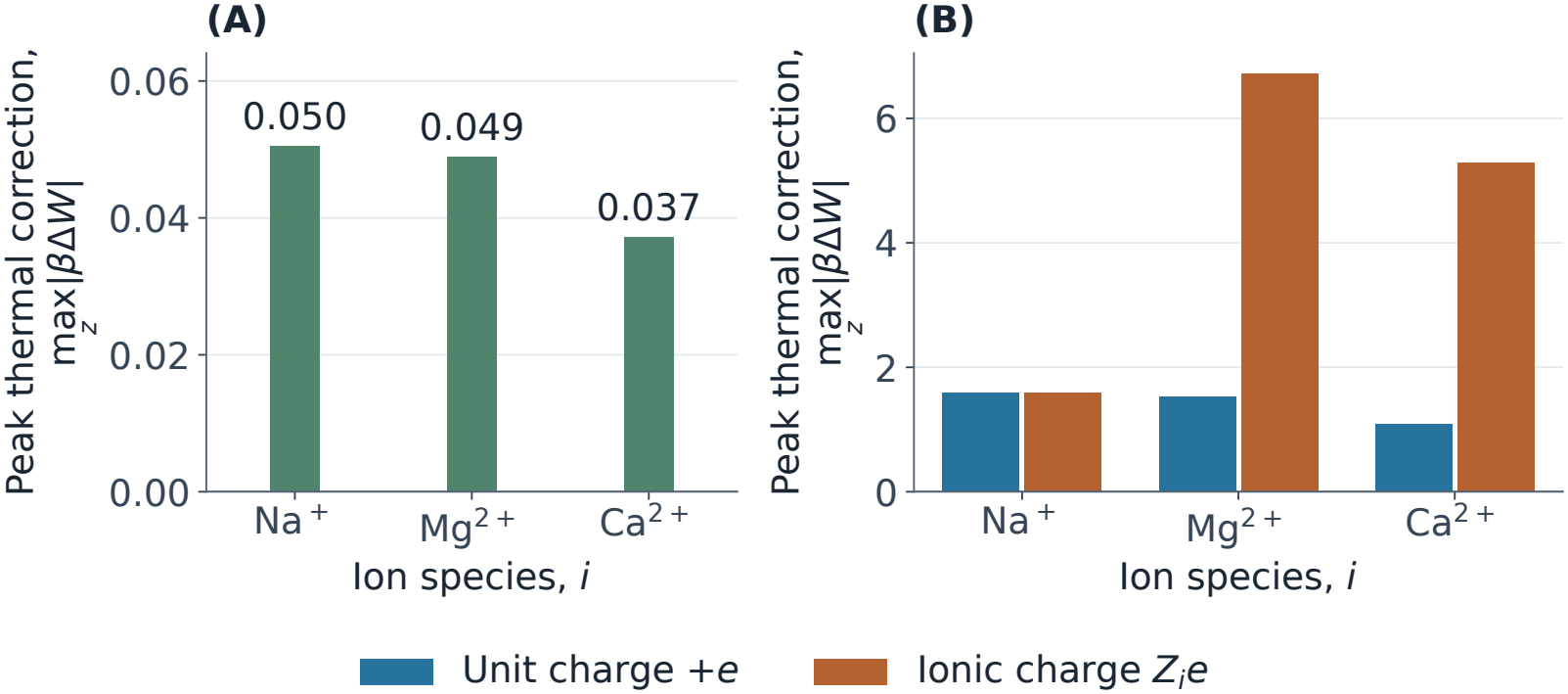
Circular point-ion controls at 310 K. The plotted quantity is the peak quantum correction in thermal units, max_*z*_ |*β*Δ*W*|, for ion species *i*. (A) Confinement alone with *U*_*i*_ = 0. (B) The comparison between unit charge (*U*_*i*_ = *eϕ*) and each ion’s physical charge (*U*_*i*_ = *Z*_*i*_*eϕ*). Both charge assignments give the same energy for sodium.

When all three ions have charge +*e*, sodium and magnesium give similar corrections and have similar nuclear masses. Raising the test charge to +2*e* makes favorable electrostatic positions more strongly preferred at equilibrium and increases the corrections for magnesium and calcium in this field. These two ions have identical classical position profiles because they share the domain and energy, and the mass factor from the momentum integral is constant along the pore. Magnesium’s larger quantum correction then reflects its lower mass (Fig. 8B).

Higher temperatures shorten the thermal de Broglie wavelength, the scale for quantum delocalization, and reduce the Boltzmann preference for low-energy positions. We vary temperature with the test-charge energy held fixed. The peak correction in thermal units decreases for sodium, magnesium, and calcium (Fig. 9A). For each ion, the larger of the two absolute percentage changes in passage time also decreases at every step in temperature (Fig. 9B).

**Figure 9:**
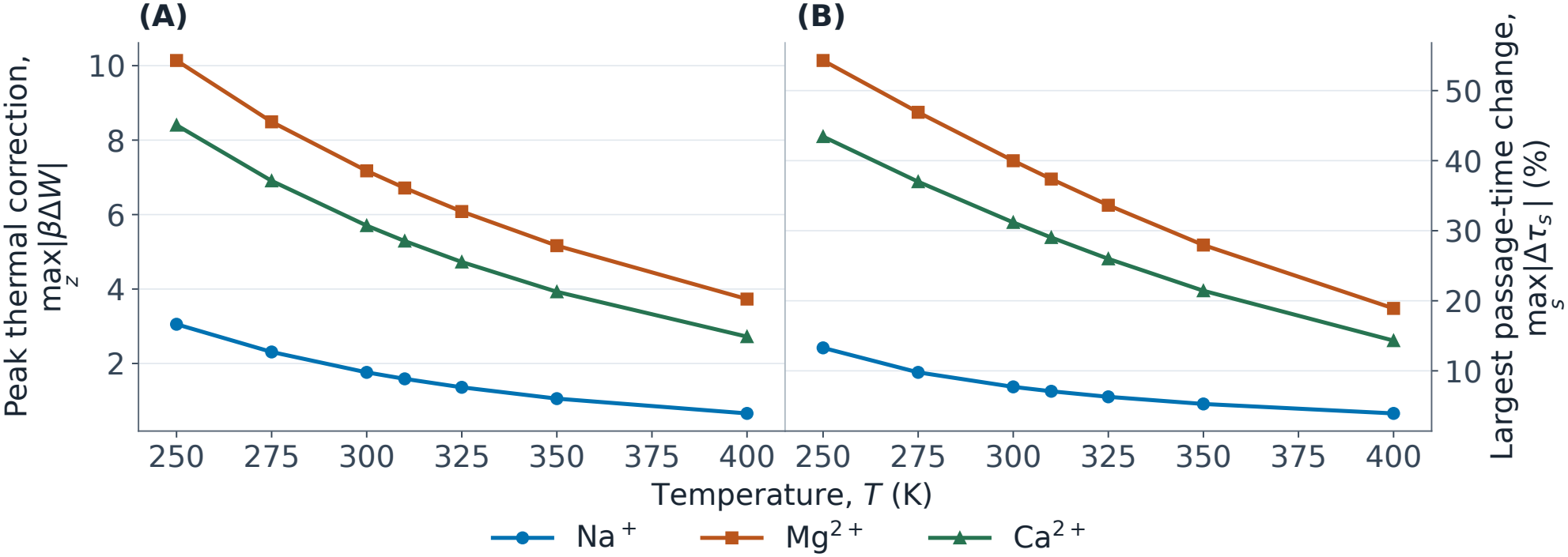
Temperature *T* dependence in the circular point-ion reference at fixed test-charge energy *U*_*i*_. (A) Peak correction in thermal units max_*z*_ |*β*Δ*W* |. (B) Largest absolute percentage change in mean first-passage time, max_*s*_ |Δ*τ*_*s*_|, where *s* selects the exit interface. The electrostatic energy is fixed at its *T*_PB_ = 310 K value.

Magnesium’s dimensionless peak decreases across the temperature sweep. For a fixed, smooth energy surface, the dimensionless curvature term in the Wigner–Kirkwood expansion scales as *β*^2^. The term from the squared gradient scales as *β*^3^. In the pore, temperature changes the transverse free energy itself, and both the smooth terms and the sharp entrance feature shape the peak response.

### 3.4 Pore fluctuations and kinetic coupling

The prescribed mode amplitude produces a geometry-dependent displacement of the entrance feature. The ensemble profile has a weaker negative extremum than the ensemble mean of the signed individual extrema (Fig. 10A). The intracellular time changes differ between the fast and slow limits. The whole-pore mode of the same amplitude produces a different displacement at the entrance because of the endpoint constraints (Fig. 10B).

**Figure 10:**
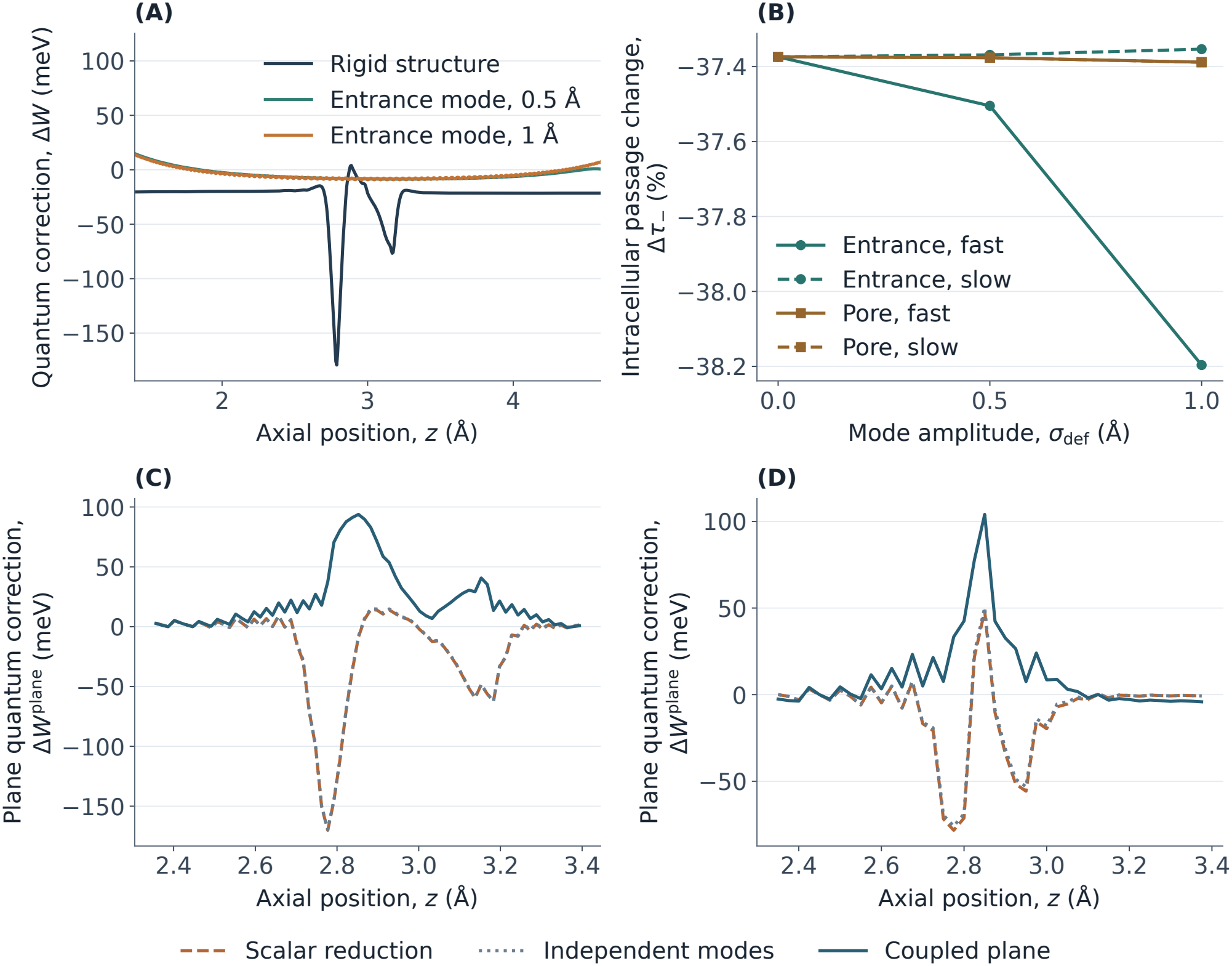
Pore fluctuations and coupling between axial and transverse motion in the point-ion reference model. (A) Δ*W* (*z*) for the rigid reference and fast-exchange entrance ensembles. (B) Intracellular passage-time change Δ*τ*_*−*_ versus deformation amplitude *σ*_def_ for two modes and both exchange limits. (C) Coupled and separated corrections Δ*W* ^plane^(*z*) on the refined entrance plane with the structural wall. (D) The same comparison with atomic-sphere exclusion. Panels C and D use a common axial reference near 2.45 Å within each comparison.

Near the widening on the refined disk plane, the scalar and independent-band corrections are negative. The coupled correction at the same sample is positive (Fig. 10C). For the atomic-union plane, the scalar and coupled corrections also have opposing signs at the scalar extremum (Fig. 10D). The opposing signs persist across the tested disk meshes and the orthogonal-plane test. The size of the effect depends on the boundary and potential. This localized plane calculation retains one transverse coordinate. It establishes a coupling effect for that problem and motivates a three-dimensional benchmark before assigning a corrected passage time.

### 3.5 Second-ion interactions and isotope effects

In the screened pair benchmark, classical repulsion adds an interaction free energy and separates the ions along the axis. The barrier comparison follows release from the occupied region within the Y1085–E1047 reference window. For a singly tracked ion in the pair, repulsion lowers the intracellular and extracellular barriers for sodium and the divalent ions. The divalent intracellular barriers remain large. The benchmark therefore identifies the scale on which a second ion can alter escape from a well in the fixed continuum field.

Quantum motion increases the pair interaction free energy for all three ions, with a larger increment for magnesium and calcium than for sodium. For a separate diagnostic, we hold the pair action at its root value as the single-ion potentials vary along the paths. Allowing the repulsion to vary along the quantum paths changes the increment only slightly. This comparison attributes most of the quantum increment to shifts in equilibrium occupation. For free identical ions at physical separations of at least 2 Å, the ratio of the exchange and direct thermal kernels is exp 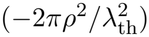 and is exponentially small for the three masses at 310 K. The thermal benchmark treats the axial coordinates as labeled and uses the prescribed repulsion. The slab-restricted paths provide a comparison of pair repulsion at fixed single-ion profiles.

On the fixed classical transverse profiles, the lighter isotope lowers both exit barriers for ^24^Mg/^26^Mg, ^40^Ca/^44^Ca, and ^6^Li/^7^Li. The changes in mean passage time remain small at common diffusivity. The larger local isotope differences occur at positions whose contributions to the passage integral are small. The leading 1*/m*_*i*_ scaling expresses the isotope difference as a fraction of the light-isotope correction. Barrier-only rate estimates depend on the size of the quantum barrier shift, while the calculated passage times also include the widths of the contributing regions.

At matched grids and boundaries, the extended calculation reproduces the four-site response. The extended matrix resolves the axial variation of dielectric charging and screened repulsion (Table 5). The interaction ranges use distinct probes with disjoint truncated Gaussian supports. Pairs with the same axial separation have different screened energies at different positions. The pair response therefore depends on axially varying dielectric and screening properties as well as on separation. The pair map describes on-axis linear response for the specified charge shape. The one-ion insertion landscape uses the spatial dielectric and hydration energy (Eq. 1). The thermal two-ion benchmark uses the prescribed screened repulsion, which identifies the effect of repulsion at fixed single-ion profiles. A joint two-ion landscape across the pore would need off-axis pair response and joint coordination of the two hydration shells.

**Table 5.** Axial response for 16 fixed divalent Gaussian probes. The mesh spacing is *h* and the lateral half-width is *b*. Pair energies *v*_*ij*_ use distinct sites with disjoint charge supports. The self-energy *S*_*σ*_ is relative to homogeneous water with the same mesh and box.

| *h* (Å) | *b* (Å) | *S*<sub>*σ*</sub> (meV) | *v*<sub>*ij*</sub> (meV) |
|---|---|---|---|
| 0.25 | 8 | 65–438 | 3–928 |
| 0.25 | 10 | 66–482 | 5–1011 |
| 0.125 | 8 | 64–430 | 3–914 |

## 4 Discussion

### 4.1 Origin of the local quantum correction

In the circular point-ion reference, the intracellular widening can expose a small set of off-axis positions whose favorable electrostatic energy dominates the transverse Boltzmann sum, the sum of equilibrium weights across the pore. This gives the effective axial free energy a steep feature even where the centerline potential varies smoothly. In an imaginary-time path description, paths rooted on the narrower side can sample that adjacent region and gain thermal weight [22]. These paths represent the spatial spread of a thermal quantum state [22]. This explains why the peak can occur on a slice that clears the prepared atomic spheres while its amplitude depends on near-wall access in a neighboring slice. A circular inset removes access uniformly around the axis. An atomic boundary preserves sphere-free off-axis space and changes which favorable positions are sampled.

An abrupt change in transverse access also changes the shape of the transverse states. The coupled diametric-plane calculation shows that this change can alter the local quantum correction when axial and transverse motion are treated together. The plane provides a test of the scalar reduction near the widening and resolves one transverse coordinate. Its result helps identify the entrance feature as a joint consequence of local geometry, electrostatics, and the kinetic coupling of the accessible states. As the pore opens, the quantum state can overlap a region that carries much more classical weight. The rapid gain in favorable positions explains why the axial response is most sensitive to the shape and hydration of the wall here.

### 4.2 Regional control of diffusive escape

The constriction affects escape when it lies on the route from the occupied well. An ion can also revisit sites behind its starting position before it reaches the target interface. The passage calculation gives greater weight to regions where the ion tends to reside. Dielectric charging and restricted water coordination both shift the relative depths of wells. They can therefore move the starting well to a position from which an ion crosses the constriction toward a different interface. In the finite-ion magnesium calculation, the time averaged over classical starting positions can be governed by a sparsely occupied well whose occupants take a long time to escape. This explains how starting from the global minimum and averaging over an equilibrium population can give different extracellular responses. A position-dependent diffusivity would further reweight the contributions of different pore regions [37, 38].

The entrance is a separate example of the difference between a local correction and its contribution to passage. Its small weights in the equilibrium and passage integrals explain why smoothing an entrance feature has little influence on release from the modeled basin. In the prescribed deformation ensemble, an average of conformation-specific passage times gives extra weight to slow conformations, while an equilibrium profile pooled under fast exchange combines their occupation weights. The exchange regime thus governs how an ensemble of open structures contributes to the measured release time.

### 4.3 Mass dependence and isotope effects

In the point-ion comparison, magnesium and calcium share the energy and domain at fixed valence. Their different nuclear masses therefore account for their different thermal delocalization. For the finite ions, differences in water coordination also change the energy landscape. Isotope substitution preserves charge and the classical equilibrium distribution at fixed interactions and provides a more direct probe of the mass dependence [39]. The smooth-potential mass factor gives a light-minus-heavy difference equal to 1 −*m/m*^*′*^ of the light-isotope correction for masses *m < m*^*′*^. The axial isotope calculation tests how this mass dependence propagates into equilibrium occupation and the escape integral, with the sodium landscape as a fixed monovalent reference for lithium. The full passage calculation also weights the widths of the well and exit region. A flux measurement would also include any isotope dependence of diffusivity [40].

### 4.4 Electrostatic response to ion occupancy

An inserted charge polarizes the surrounding medium and changes the statistical weights of configurations available to a second ion [7, 30, 31]. In the screened pair benchmark, repulsion shifts configurations away from simultaneous occupation of the deepest region. The resulting change in the marginal barrier connects ion occupancy to release from the open pore. In a channel with several potential binding positions, two ions at the same separation may experience different dielectric environments. The spatial response of the continuum field therefore gives a physical basis for position-dependent interactions and for asymmetric effects of occupancy on escape.

Protein coordination links occupancy to the chemistry of the constriction. A protein oxygen can replace a water ligand, and the rearrangement of an ion’s coordination environment can change the weights of neighboring binding configurations [41, 42]. The prescribed shell term captures the cost of losing water access at fixed coordination geometry. The constriction restricts the orientations in which coordinating waters can accompany the ion. The free-energy cost of restricted access therefore couples local pore shape to ion retention. A cavity can admit the ion center while limiting the orientations in which its waters fit together. The water-access cost can therefore shift the favored site even when the ion-center domain stays fixed. Explicit water and protein ligands would resolve how compensation by protein oxygens changes that cost and would permit a separate treatment of quantum motion in the ion and its surroundings [11, 15, 18].

### 4.5 Biological implications for TRPM7 permeation

Neutralizing E1047 reduces divalent permeation and establishes the influence of filter charge on the conduction of magnesium and calcium [3]. The calculations identify two routes by which a filter perturbation could change conduction. A shift in the preferred well changes the route that an occupied ion takes to reach an interface. A shift in the coordination cost at the constriction changes the free-energy barrier on that route. Those routes can contribute differently to ion binding and release, so single-channel current and open probability constrain different parts of the interpretation of a perturbation [1, 5]. A filter mutation that stabilizes an occupied site could increase its occupancy and extend the time each ion spends there. Measuring ion residence alongside current during openings could help identify changes in retention at the filter.

Relating release to the contribution of TRPM7 to mineral homeostasis requires the probabilities of ion entry, pore occupancy, and channel opening under the relevant cellular conditions [4, 5]. The estimates here isolate release within the naltriben-open conformation. They locate the chemical and geometric features that can couple ion coordination to escape in that state.

## 5 Conclusion

Dielectric self-energy and water coordination can change the classical well from which a divalent ion escapes. The location of that well governs which parts of the axial quantum correction enter the directional passage integral. The circular point-ion reference shows that a large entrance correction can have little influence on release from an interior well. In the finite-ion model, the extracellular response can reverse when the preferred well crosses the constriction. The fixed-geometry calculations establish how ion insertion and pore shape condition the kinetic effect of nuclear delocalization. In this setting, comparisons across isotopes can probe ion mass at fixed chemistry. Comparisons across different ion species also reflect their coordination and preferred sites. These effects connect the electrical and chemical properties of the open filter to how long an ion stays in the channel.

## Notes

### Competing Interest Statement

The authors have declared no competing interest.

