## Supplementary Materials for "Nuclear quantum effects on ion permeation through the open transient receptor potential melastatin 7 pore: A continuum-electrostatics estimate"

The open TRPM7 structure defines the accessible space and the electrostatic environment for the ion. The calculations below test how field preparation, electrostatic resolution, thermal sampling, and pore geometry affect three observables. We distinguish the local quantum free-energy peak, the quantum shift in each exit barrier, and the relative change in mean first-passage time (MFPT) to each interface. These quantities probe different parts of the free-energy landscape.

#### 1 Structure preparation and electrostatic model

We first place the structure in a coordinate system that can be used for both the field and the pore radius. We used biological assembly 1 of the naltriben-open mouse channel in Protein Data Bank (PDB) entry `pdb_00008si5` [1]. We retained the four protein chains of polymer entity 1 from the Protein Data Bank exchange macromolecular Crystallographic Information Framework (PDBx/mmCIF) file. We removed resolved calcium, water, ligands, and other nonprotein components and left gaps at unresolved residues. To define the direction of passage, we fitted the pore axis to the centers of the rings of corresponding $\alpha$-carbon atoms (C$\alpha$) in the four subunits. We used the orientation from the Orientations of Proteins in Membranes (OPM) database and projected the membrane boundary markers onto the fitted axis [2]. The intracellular and extracellular interfaces define $z = 0$ and $z = L$, respectively.

The structure sets the accessible radius along this axis. The I1093, Y1085, and E1047 reference planes mark the mean axial C$\alpha$ positions of the four subunits at 4.54, 13.59, and 22.98 Å, respectively. To set the circular wall for the thermal calculation, we found the radial distance to the closest heavy-atom van der Waals sphere among residues 839–1146 that intersects each plane. The structural sphere radii were 1.70 Å for carbon, 1.55 Å for nitrogen, 1.52 Å for oxygen, and 1.80 Å for sulfur [3]. These planes are spaced 0.1987 Å apart. Linear interpolation gives a continuous radius and allows its slope to change at the tabulated planes. This centered clearance uses a prescribed axis. A movable probe, as in the HOLE method, also permits lateral displacements [4].

We used PDB2PQR (3.7.1) to generate a PQR structure file containing atomic coordinates, charges, and radii [5–7]. We used the parameters for solvation energy (PARSE) for the atomic charges and radii [8]. We used PROPKA (3.5.1) to estimate the initial protonation states at pH 7.4 [9–11]. Preparation included steric clash removal, hydrogen addition, and hydrogen-bond optimization. We retained the chain identifiers and permitted coordinate changes during optimization. The initial protonation calculation reached its ten-iteration limit and assigned a deprotonated E1047 to one protomer. We defined the reference protonation microstate by assigning the deprotonated glutamate charges to all four E1047 residues. The prepared protein had a net charge of $+40e$ with $+10e$ per chain. The neutral histidine

*
†

at position 191 retained its chain-specific tautomer assignment. Alternative protonation and coordinate preparations were outside the scope of the sensitivity tests.

The Adaptive Poisson–Boltzmann Solver (APBS, 3.4.1) provided smoothed molecular-surface coefficients, water-reference dielectric maps at staggered grid edges, and a fixed-charge map [6, 12, 13]. The solvent probe radius was 1.4 Å, the surface spline window was 0.3 Å, and the surface discretization density was 10 points per square angstrom. We distributed fixed charges by quintic B-spline discretization (`spl4`) [14]. The relative permittivities of the protein, membrane, and pure water at 310 K were 4, 2, and 74.2433, respectively [15].

To exclude isolated cavities from the membrane, we selected six-connected regions within the slab that reached a lateral box face, were accessible to a 2.7 Å probe, and lay outside the hydrated pore. The hydrated pore was defined from the clearance to positive-radius PQR atomic spheres at each axial position. The clearance was sampled at intervals of approximately 0.05 Å and interpolated linearly to each grid. We converted the signed distance from the selected membrane region to a membrane fraction with a cubic smooth step of width 1.0 Å. The hydrated pore used the same width for radial and axial transitions. At each staggered grid edge, we derived a protein fraction from the reference reciprocal permittivity. The fraction of the edge outside the hydrated pore scaled the protein and membrane fractions. The membrane fraction occupied only the volume left after assigning protein, and the remaining volume was assigned to water. The local permittivity was the harmonic mean of the three permittivities, weighted by these fractions.

We used 150 mM each of $Na^+$ and $Cl^-$ in both reservoirs, with ion radii of 2.0 Å and zero imposed transmembrane voltage. The common 2.0 Å radius defined the base ion-accessibility map. We reduced this accessibility by the local membrane fraction and suppressed mobile ions inside the hydrated pore, extending the exclusion by 2.0 Å past each interface before the 1.0 Å axial transition. This map controls the mobile-ion term independently of the dielectric map.

For a symmetric monovalent electrolyte, the continuum Poisson–Boltzmann (PB) equation that we discretized is [6]

$$-\nabla \cdot [\varepsilon_0 \varepsilon_r(\mathbf{r}) \nabla \phi(\mathbf{r})] + 2 e n_{\mathrm{b}} \chi(\mathbf{r}) \sinh\left( \frac{e\phi(\mathbf{r})}{k_{\mathrm{B}} T_{\mathrm{PB}}} \right) = \rho_{\mathrm{f}}(\mathbf{r}), \tag{1}$$

where $\varepsilon_0$ is the vacuum permittivity, $\varepsilon_r$ is the relative permittivity, $n_{\mathrm{b}}$ the bulk number density of either ion species, $\chi$ the dimensionless ion accessibility, $\rho_{\mathrm{f}}$ the fixed charge density, and $T_{\mathrm{PB}}$ the temperature used in the Poisson–Boltzmann calculation. Each ionic species has a reservoir concentration of 150 mM.

#### 2 Electrostatic grid checks

In the figures, $z$ is axial position and $(x, y)$ are transverse coordinates. The original potential is written as $u = e\phi/(k_{\mathrm{B}} T_{\mathrm{PB}})$. A test case is indexed by $j$, and $\delta_j Q = Q_j - Q_{\mathrm{ref}}$ denotes the difference for any plotted quantity $Q$. The reference is the paired calculation specified for each test. For a set of sample positions $R$, $\|\delta_j u\|_{\infty,R}$ is the largest absolute potential difference over $R$. The bracket $\langle\cdot\rangle_{\mathrm{C}}$ denotes the weighted classical average in the stated reference domain. The local correction has peak magnitude $A_W = \max_z |\Delta W(z)|$. We write $h_z$ for axial slice spacing, $h$ for electrostatic grid spacing, and $h_{\mathrm{g}}$ for the geometric mean of a pair of grid spacings. The paired-field root-mean-square difference is $D_{\mathrm{RMS}}$, and the root-mean-square error against an exact solution is $E_{\mathrm{RMS}}$. Changes of a passage-time percentage are in percentage points (pp).

The charge of the entire protein helps set the potential near the pore. We solved the nonlinear equation first in a large box, then in a series of smaller boxes to resolve the potential near the pore at finer spacing. At each stage, we used nested finite-difference multigrid calculations (`mg-manual`). We supplied the precomputed coefficient maps through the APBS map interface [16]. We set the potential at the outer boundary to the single-sphere Debye–Hückel Dirichlet values. For each subsequent box, we interpolated the potential from the previous, larger box to specify values on its boundary. For local boxes whose faces cut through the structure, the coefficient grids extended beyond each face by at least the largest atomic radius plus three coefficient-grid spacings. The extended coefficient grids contained every node used to assign charge near the boundary.

The outer, whole-protein, and transmembrane boxes had side lengths of 320, 192, and 96 Å, respectively, and spacings of 2.0, 1.5, and 0.60 Å. The outer and whole-protein boxes were centered at $(0.0025, -0.0035, -23.199)$ Å. The remaining boxes were centered at $(0, 0, 15.0)$ Å, except in the grid translation test. The box that supplied boundary values near the pore measured $40.96 \times 40.96 \times 60$ Å$^3$ with $129 \times 129 \times 193$ nodes, giving spacings of $0.3200 \times 0.3200 \times 0.3125$ Å.

The two most local calculations used the same boundary potential on the $24 \times 24 \times 40$ Å$^3$ box. The coarse grid had $97 \times 97 \times 161$ nodes at 0.25 Å spacing. The fine grid had $129 \times 129 \times 193$ nodes with spacings of $0.1875 \times 0.1875 \times 0.2083$ Å. The nonlinear solver tolerance was $10^{-6}$ at each stage except on the fine grid, which used $10^{-8}$. We used trilinear interpolation of the fine grid to evaluate the potential where the thermal calculation samples the cross-section. We stored the potential as $u = e\phi/(k_B T_{PB})$, so the test-charge energy is $U_i = Z_i k_B T_{PB} u$.

We also compared linearized and nonlinear screening on the same 0.25 Å pore grid. Both solves used the same fixed charge, staggered permittivities, mobile-ion accessibility, and nonlinear parent potential as the boundary condition. The linear solve replaced $\sinh u$ by $u$ in the mobile-ion term. On the common 0.50 Å sample, which includes 61 centerline and 6,269 hydrated-core nodes, the root-mean-square potential difference was 0.489 in units of $k_B T_{PB}/e$ on the centerline and 0.526 in the hydrated core. The largest core difference was 1.12 in those units. The core root-mean-square difference corresponds to 28.1 meV for a divalent test charge. Fixing the same parent field isolates the local effect of the screening law. The thermal calculations use nonlinear electrostatics throughout the focusing sequence.

To test grid spacing, we compare the two solutions on the same domain. Three further tests translate the lattice by 0.125 Å in each direction, set the boundary values from a $30.72 \times 30.72 \times 50$ Å$^3$ pore parent domain with $97 \times 97 \times 161$ nodes, or enlarge the child domain to $32 \times 32 \times 48$ Å$^3$ at a grid spacing of 0.25 Å. The standard 0.25 Å child solution serves as the reference for each of the three tests. Each comparison gives a distinct axial profile of potential differences along the pore centerline (Fig. S1A). Differences in the potential away from the axis can affect the calculation because the classical ion distribution varies across each accessible cross-section. We calculated the root-mean-square (RMS) potential difference over each cross-section, using the classical Boltzmann weights of a divalent ion in the corresponding reference potential. The statistic thus gives greater influence to differences at locations that the ion is more likely to occupy in the reference calculation (Fig. S1B).

For the archived grid comparisons, we selected pore-interior points with water-like permittivity. Each selected point must lie at least 1 Å outside the PQR atomic spheres, and the permittivity in all three directions must reach at least 95% of the bulk-water value. We reconstructed this fixed set of grid comparison points from the structural pore radius, PQR atomic spheres, and staggered dielectric maps. For acceptance, at least 90% of the clearance-qualified points had to satisfy the permittivity criterion, with representation on every axial plane. The selected set contains 12,437 points distributed over all 121 comparison planes. The pore centerline also meets the dielectric criterion in every comparison plane. We recomputed the potential statistics from cubically interpolated values in the hydrated core, on the centerline, and for the three clearance thresholds of 0, 0.5, and 1 Å. The four comparisons meet the numerical thresholds in the hydrated core and along the centerline, including the 99th percentile (P99) of the absolute core differences (Table S1). The root-mean-square cutoff of $0.20\, k_B T_{PB}/e$ corresponds to 5.3 and 10.7 meV per monovalent and divalent charge, respectively. The downstream results quantify sensitivity at the scale of the quantum corrections. At the selected pore points, the largest difference between cubic and trilinear interpolation was 0.00515 $k_B T_{PB}/e$. The broader sample outside the PQR atomic spheres extends closer to the dielectric interface. The largest difference between paired potentials in that sample is 2.71 $k_B T_{PB}/e$ (Fig. S1C). We used the entire accessible cross-section for the thermal comparisons to include the sensitivity of states near the wall.

We independently solved the electrostatic problem on outer boxes with side lengths of 256 and 320 Å and a common spacing of 2.0 Å. We interpolated both potentials trilinearly to the 98,306 boundary nodes of the whole-protein child domain. The differences at these boundary nodes take both positive

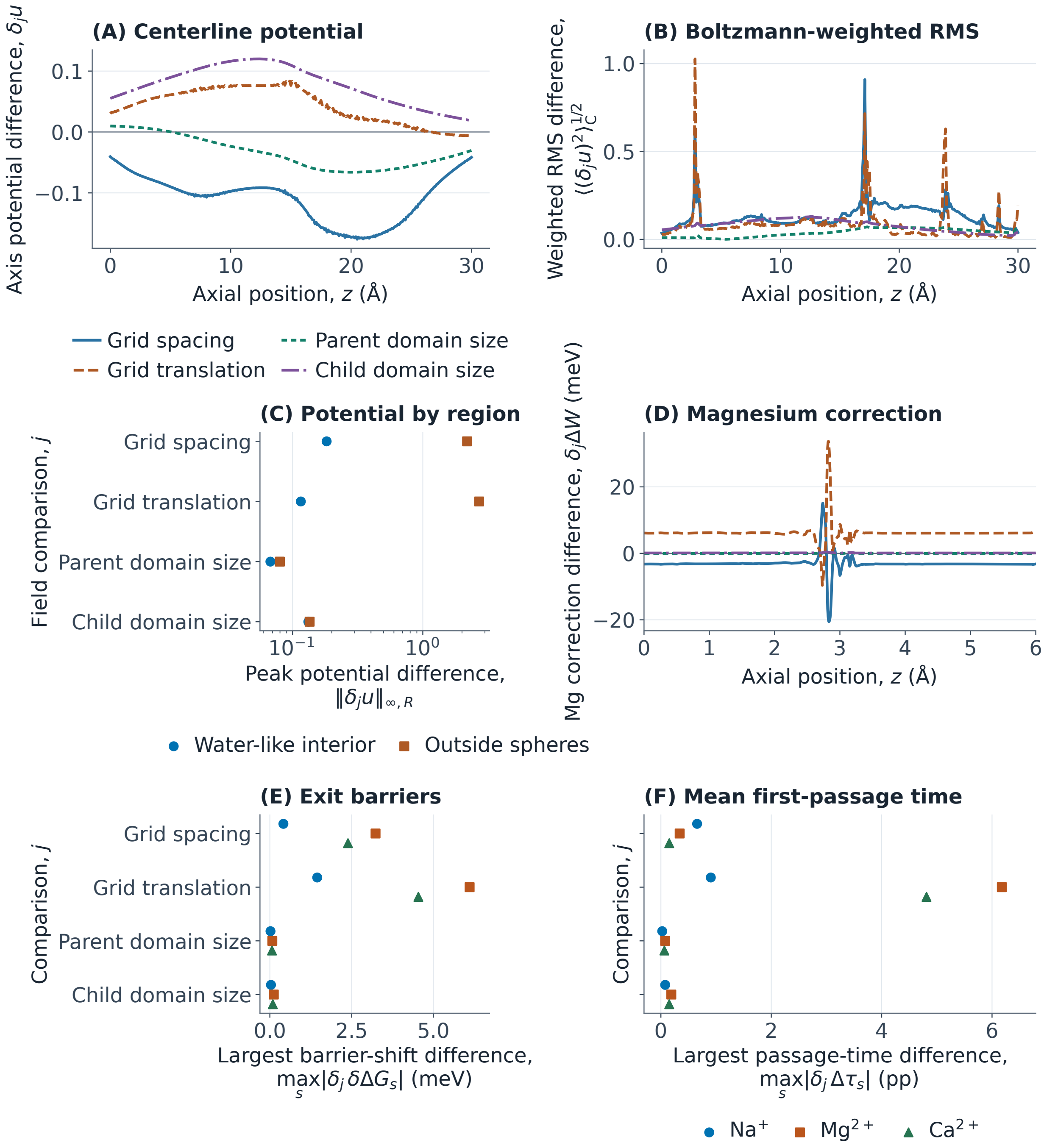


**Figure S1:** Electrostatic grid sensitivity at 310 K. (A) Signed dimensionless potential difference $\delta_j u(0,0,z)$ along the axis for comparison $j$. The spacing comparison subtracts the fine field from the coarse field. Translation and domain tests subtract the standard coarse field. (B) Cross-sectional RMS difference $\langle(\delta_j u)^2\rangle_{\mathrm{C}}^{1/2}$ with divalent classical weights. (C) Maximum absolute difference $\|\delta_j u\|_{\infty,R}$ in the water-like and sphere-free regions $R$. (D) Signed magnesium correction difference $\delta_j \Delta W$ along $z$. (E) Maximum directional exit-barrier difference $\max_s |\delta_j \delta\Delta G_s|$. (F) Maximum passage-time difference $\max_s |\delta_j \Delta\tau_s|$.

**Table S1:** Potential differences in the hydrated pore core and the recorded grid acceptance thresholds, expressed in $k_B T_{PB}/e$. The shape RMS is the root-mean-square difference after removal of the mean signed difference. The centerline maximum is evaluated separately on the 121 axial points.

| Comparison | Core RMS | Core P99 | Core max. | Shape RMS | Axis max. |
|---|---|---|---|---|---|
| Grid spacing | 0.100 | 0.176 | 0.183 | 0.038 | 0.175 |
| Half-cell grid translation | 0.054 | 0.089 | 0.116 | 0.035 | 0.077 |
| Parent domain size | 0.038 | 0.067 | 0.068 | 0.022 | 0.066 |
| Child domain size | 0.087 | 0.128 | 0.132 | 0.040 | 0.120 |
| Acceptance limit | 0.200 | 0.300 | 0.500 | 0.075 | 0.250 |

and negative values. When the permittivity, fixed charge, and ion-accessibility maps are identical, the monotonicity of the mobile-ion term makes the zeroth-order coefficient in the linear elliptic equation for the potential difference nonnegative. The maximum principle then bounds the difference throughout the interior by its minimum and maximum on the boundary [17]. The same bound applies at subsequent focusing stages if the child-domain coefficients are fixed and interpolation preserves the bounds on the boundary values. The resulting bound on the absolute potential difference is $3.025 \times 10^{-4}\, k_B T_{PB}/e$, which gives an upper bound of 0.0162 meV on the energy change for a divalent test charge before thermal integration. This result bounds the change in electrostatic potential due to the placement of the outer boundary.

We used the circular point-ion thermal calculation at 310 K for each focused potential map. The electrostatic lattice controls the resolution of near-wall energies. Independently, the axial slice grid places changes in accessible cross-section into the thermal trace. Both the change in grid spacing and the half-cell translation affect the local $Mg^{2+}$ correction near the entrance (Fig. S1D). Across all ions, the change in grid spacing alters the peak magnitude by up to 9.53 meV and the quantum correction to an exit barrier by up to 3.23 meV. Translating the grid changes the quantum correction to the exit barrier by at most 6.11 meV (Fig. S1E). The MFPT percentages differ from their respective reference results by up to 0.66 percentage points in the spacing test and 6.18 percentage points in the grid translation test (Fig. S1F). The prepared protein and hard-wall geometry are identical across the comparisons, and the imposed voltage is zero in every case. The two final grids use unequal refinement ratios along and across the axis and different stopping tolerances. Their differences quantify the sensitivity to those choices. Section 5 examines an isotropic refinement series with common solver tolerances and coefficients.

### 3 Resolution of the thermal calculation

For the circular point-ion reference, we resolve motion across the pore separately from motion along its axis. The two directions require different numerical treatments because the radius varies along the pore and each accessible cross-section is circular. The atomic-union and finite-ion settings appear in Section 6.

For the classical calculation across the pore, we evaluated the weighted area integral by 64-point Gauss–Legendre quadrature in the squared radial coordinate and by sampling 256 points in the periodic angular coordinate. For the quantum partition function, we discretized the transverse Hamiltonian on a Cartesian grid with a symmetric five-point operator. Where a grid edge met the circular Dirichlet wall, the intersection set the boundary diagonal weight. A minimum wall distance of one fifth of the grid spacing limited the spectral span of the discrete operator.

The quantum partition function sums the Boltzmann weights of the transverse energy levels. We evaluated the corresponding matrix trace deterministically with a Chebyshev polynomial expansion and graph-coloring probes. Chebyshev expansions provide a spectral representation of matrix functions [18]. We calibrated the trace by computing a second trace on the same grid with $U_i = 0$, using the continuum Bessel spectrum of the reference disk and restoring the continuum classical integral over the cross-section. With subscript $\ell$ denoting the lattice, the logarithm of the calibrated trace is $\ln Z^{Q}_{\perp,0} + \ln\langle e^{-\beta U_i}\rangle_\Omega + \ln[\mathrm{Tr}(e^{-\beta H_{\ell,i}})/\,\mathrm{Tr}(e^{-\beta H_{\ell,0}})] - \ln\langle e^{-\beta U_i}\rangle_\ell$. The continuum average weights the cross-section by area, and the lattice average weights the interior nodes equally. For a constant potential, the calibration recovers the continuum disk trace up to spectral truncation. With a spatially varying potential, the mesh-refinement test measures the sensitivity of the calibrated result. The polynomial and probe tolerances control the trace evaluation on a given lattice.

We evaluated the transverse correction at 489 axial positions, with axial intervals no larger than 0.115 Å and a transverse mesh spacing of 0.065 Å. The set of axial positions contains the tabulated radii and the planes of the APBS grids. We evaluated the classical integral at axial intervals no larger than 0.0275 Å. We interpolated the resulting classical and quantum transverse free energies separately onto the axial thermal grid with shape-preserving cubics. We discretized the axial Hamiltonian on a 0.00625 Å mesh aligned to both membrane interfaces and extended the effective axial potential at its endpoint value for 2 Å beyond each interface. The numerical wave-function derivative vanishes at the remote edges. The density is normalized on $[0, L]$, so the constant-potential extensions are outside the passage interval. We computed the diagonal of the thermal density matrix from the spectra of local tridiagonal matrices with overlap regions of 1.35 Å. At the outer endpoints of the extended interval, we imposed a zero-normal-derivative boundary condition. To test sensitivity to axial discretization and the numerical boundary treatment, we halved the axial spacing and increased the overlap and extension lengths.

At 310 K, we used the fine potential map to repeat the transverse calculations on uniform axial grids with nominal spacings of 0.25, 0.10, and 0.05 Å. To place both interfaces on the grid without exceeding these spacings, we used intervals of 0.24793, 0.09967, and 0.04992 Å, respectively. We computed the transverse free energies at these positions and interpolated between them with linear and shape-preserving cubic functions. The slice locations and interpolation together determine the free-energy slopes sampled by the axial Hamiltonian. With cubic interpolation, the local minimum of the $Mg^{2+}$ correction near the entrance becomes deeper as the sampling interval decreases (Fig. S2A). Even on the same set of axial planes, linear and cubic interpolation produce different slopes. We also resampled the resolved transverse free energies at the same nominal 0.25 and 0.10 Å spacings to assess how the results depend on the source of the transverse values (Fig. S2B).

The piecewise-linear interpolation of the radius gives slope changes at the original tabulation points. Including only the ten tabulated radius positions near the entrance on the 0.05 Å grid increases the $Mg^{2+}$ peak magnitude from 148.0 to 170.7 meV with cubic interpolation. When we also include the nine axial planes of the fine potential map in the same interval, the peak magnitude is 170.4 meV. Most of the increase thus comes from including the original radius positions (Fig. S2A). The ranges of $Mg^{2+}$ peak magnitudes are 46.7–72.5 meV at the nominal spacing of 0.25 Å and 123.8–145.7 meV at 0.10 Å. At the spacing of 0.05 Å, the range is 142.6–148.0 meV (Fig. S2C). The MFPT ratio depends on the full profiles. It changes less in response to sampling near the entrance than does the local extremum (Fig. S2D).

The remaining tests change the axial Hamiltonian grid spacing, the overlap and endpoint extensions, the transverse grid spacing, the axial sampling together with the transverse quadrature, the transverse quadrature alone, and the interpolation of the three-dimensional potential. The sensitivity data record the signed peak correction, both barrier corrections, and both passage-time percentages for each ion and test. Because these tests use the same potential map, they isolate sensitivity to the numerical representation of the thermal problem. Finer axial sampling and classical quadrature change the largest magnitude of the local correction by up to 5.44 meV (Fig. S2E). A finer transverse grid changes the MFPT percentages by up to 1.74 percentage points. Cubic interpolation of the three-dimensional potential changes the MFPT percentages by up to 0.47 percentage points (Fig. S2F). This interpolation test samples the full accessible cross-section, including the region outside the hydrated core.

On small test problems, we checked the transverse partition function against a direct sum over all eigenvalues of the discretized Hamiltonian and the axial density matrix against a dense matrix exponential. In both benchmarks, adding a constant energy offset multiplied the result by the expected Boltzmann factor. Across the reference temperature series and controls, the MFPT percentages from exact integration over each linear segment of the interpolated profiles differ from the trapezoidal estimates by at most 0.024 percentage points. For the reflecting-boundary test, we used exact segment integration at every endpoint position to isolate the effect of the boundary position from that of the quadrature method.

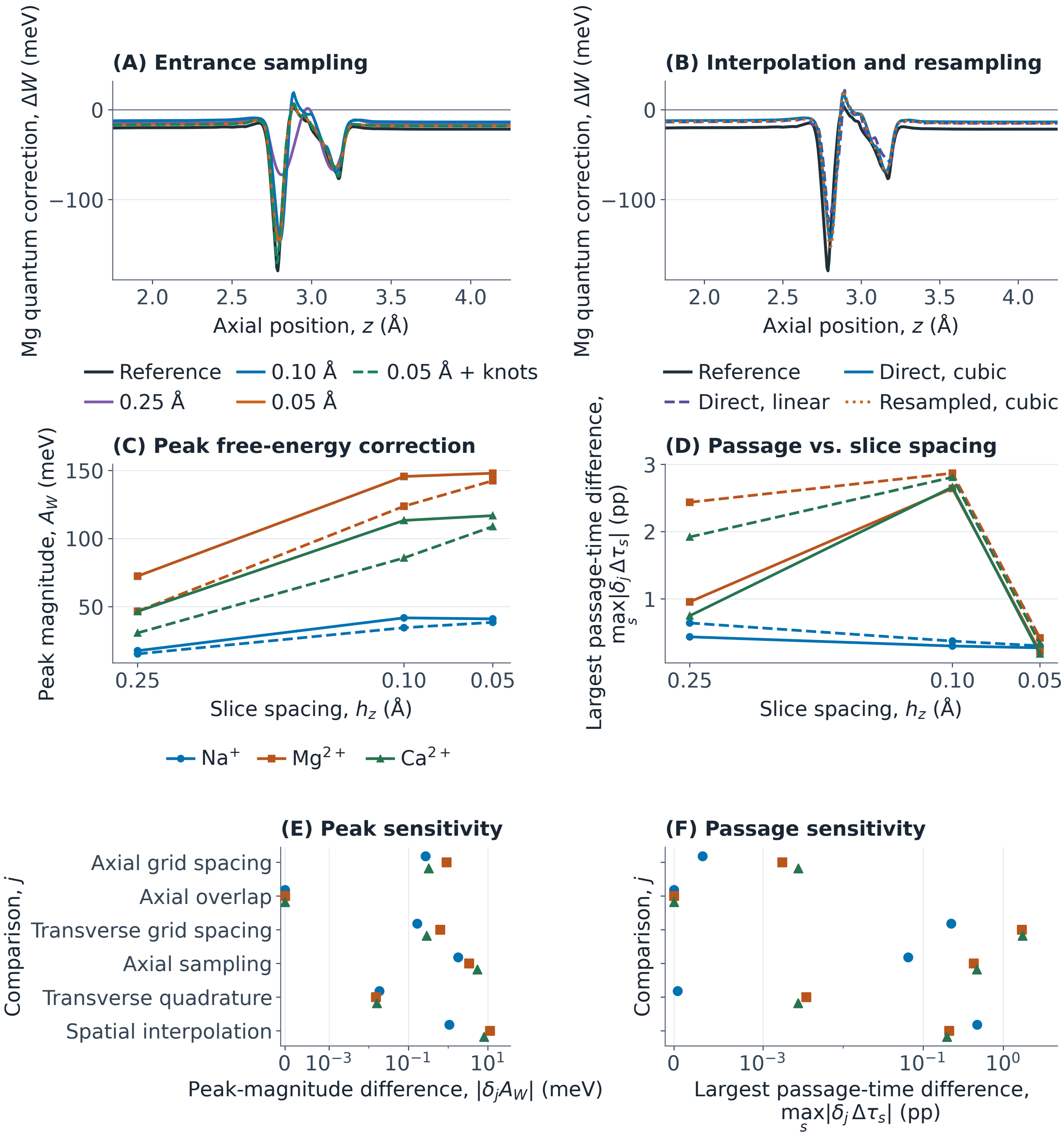


**Figure S2:** Axial sampling and thermal discretization at 310 K. (A) Magnesium correction $\Delta W(z)$ with reference and uniform slice sampling, and with radius knots added to the 0.05 Å grid. (B) $\Delta W(z)$ for linear and cubic interpolation on 0.10 Å slices and for cubic interpolation after resampling. (C) Peak magnitude $A_W = \max_z |\Delta W|$ and (D) maximum directional passage-time difference $\max_s |\delta_j \Delta\tau_s|$ as functions of nominal slice spacing $h_z$. Curves are solid for cubic and dashed for linear interpolation. (E) $|\delta_j A_W|$ and (F) $\max_s |\delta_j \Delta\tau_s|$ for numerical comparison $j$.

The tables use percentage points (pp) for differences between MFPT percentages.

**Table S2:** Changes in the thermal and transport results for matched comparisons of electrostatic potentials at 310 K. Each entry reports the largest absolute change among the three ions. For quantities defined in both directions, the maximum also spans the two target interfaces.

| Test | Free-energy changes (meV) Profile | Peak magnitude | Barrier correction | MFPT change (pp) |
|---|---|---|---|---|
| Grid spacing | 20.580 | 9.533 | 3.232 | 0.656 |
| Half-cell grid translation | 33.696 | 7.536 | 6.108 | 6.177 |
| Parent domain size | 0.254 | 0.254 | 0.075 | 0.077 |
| Child domain size | 0.400 | 0.400 | 0.120 | 0.189 |

**Table S3:** Effect of discretization and interpolation on the thermal and transport results at 310 K. Each entry reports the largest absolute change among the three ions. For directional quantities, the maximum also includes both target interfaces.

| Test | Free-energy changes (meV) Profile | Peak magnitude | Barrier correction | MFPT change (pp) |
|---|---|---|---|---|
| Axial step halved | 2.621 | 0.908 | 0.476 | 0.0028 |
| Axial overlap and extension | $< 0.0001$ | $< 0.0001$ | $< 0.0001$ | $< 0.0001$ |
| Transverse mesh refined | 5.940 | 0.628 | 1.346 | 1.739 |
| Axial sampling and quadrature | 9.707 | 5.436 | 0.076 | 0.468 |
| Transverse quadrature only | 0.206 | 0.019 | 0.021 | 0.0035 |
| Spatial interpolation | 13.507 | 11.292 | 2.697 | 0.472 |

### 4 Geometry and passage boundaries

The near-wall weight $w_{\text{wall}}(z)$ is the fraction of the classical transverse Boltzmann integral in the outer 0.5 Å of the circular cross-section. The reflecting-end displacement $\Delta z_{\text{ref}}$ is the distance through which the opposite boundary moves inward from its membrane interface.

We varied the radius with the electrostatic field held fixed to test how access to near-wall positions affects the magnesium correction. We sampled the linearly interpolated radius every 0.01 Å, extended it at constant values at both ends, and smoothed it with a Gaussian of standard deviation 0.25 Å. For the entrance-only test, we applied the full radius correction from 2.4 to 3.4 Å and tapered it to zero at 1.9 and 3.9 Å with raised-cosine weights. A separate test reduced the radius from $r(z)$ to $r(z) - 0.5$ Å throughout the pore, using the same potential and displacement for all three ions (Fig. S3A). We also calculated the share of the classical transverse Boltzmann weight contributed by the outer 0.5 Å of the reference cross-section for $Mg^{2+}$ (Fig. S3B). A large share means that a small change in radius can remove or admit positions that account for much of the equilibrium weight at that axial position.

The reference radius grows from 3.92 to 5.32 Å between $z_1 = 2.781$ Å and $z_2 = 2.980$ Å. The classical magnesium free energy falls by about 0.41 eV. The change in area contributes $-k_{\text{B}}T \ln[A(z_2)/A(z_1)] \simeq -16$ meV, and the change in the mean Boltzmann weight accounts for the remainder. At the peak of the magnesium correction, the outer 0.5 Å carries 98.2% of the classical transverse weight. Points less than 1 Å from the prepared atomic surfaces carry 99.0% of that weight.

We also compared the structural thermal wall with the atomic spheres used in the electrostatic calculation. We tested each transverse quadrature point for inclusion in any positive-radius sphere of the prepared charge-and-radius model. We integrated the area and the classical Boltzmann weight separately inside and outside this union. We repeated the test with radial and angular orders of $100 \times 400$, $256 \times 1024$, and $512 \times 2048$. At the wider entrance slice ($z = 2.980$ Å), the two highest orders give sphere-exclusion free-energy differences within 0.36 meV of one another. The sampled points at the magnesium peak

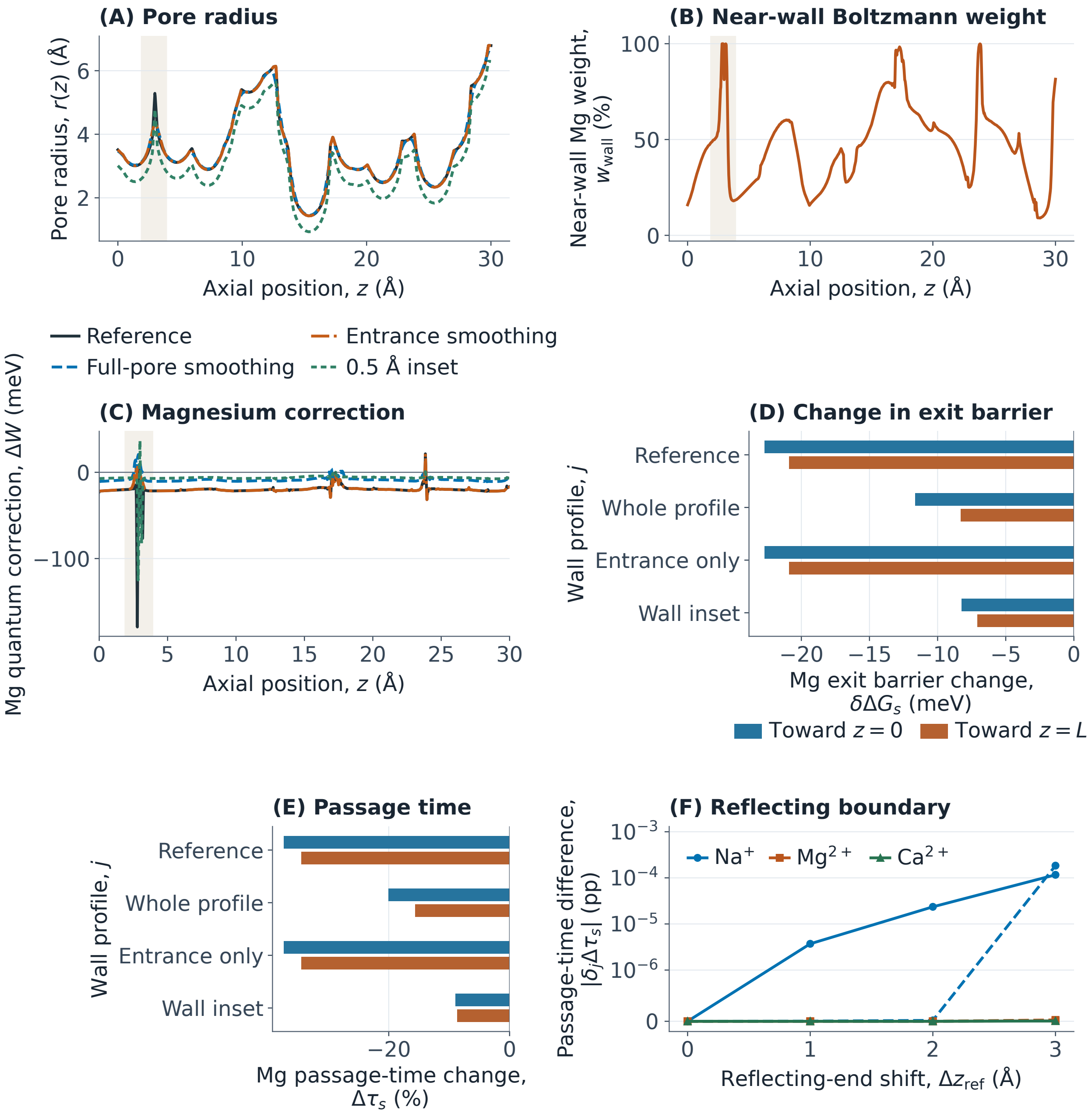


**Figure S3:** Geometry and reflecting-boundary sensitivity at 310 K. Along axial position $z$, (A) compares the radius $r(z)$ for each wall profile $j$, (B) gives the magnesium near-wall weight $w_{\text{wall}}$ within 0.5 Å of the boundary, and (C) gives the quantum correction $\Delta W$. Shading marks the entrance smoothing interval. (D) Exit-barrier change $\delta\Delta G_s$ and (E) passage-time percentage change $\Delta\tau_s$ for each geometry. (F) Passage-time difference $|\delta_j \Delta\tau_s|$ relative to reflection at the membrane interface for inward shift $\Delta z_{\text{ref}}$. Solid and dashed curves indicate target interfaces at 0 and $L$, respectively.

($z = 2.788$ Å) lie outside the spheres. At the basin and the two classical barrier maxima, this change is below 0.06 meV at the sampled positions. These pointwise comparisons diagnose how overlap changes the classical integrals. The main-text geometry comparison propagates sphere exclusion through the magnesium thermal and passage calculations.

To assess the numerical sensitivity, we halved the axial Hamiltonian step in the sphere-excluding circle and inserted a midpoint between each pair of classical quadrature planes while retaining the same transverse quantum traces. The peak changed by 3.71 meV, the barrier corrections by up to 1.13 meV, and the passage-time percentages by up to 0.03 percentage points.

After we smooth the radius only near the entrance, the $Mg^{2+}$ correction of greatest magnitude is -31.7 meV, and its extremum occurs at 23.96 Å, outside the smoothed region (Fig. S3C). The classical basin minimum stays at its reference position under entrance-only smoothing. The larger radius perturbations change the basin or exit-barrier regions and give bigger changes in the quantum barrier correction (Fig. S3D). These changes in the free-energy profiles affect the MFPT estimates. Smoothing the radius only near the entrance changes the magnesium MFPT percentage by at most 0.00063 percentage points relative to the reference. Across the three ions, sodium gives the largest such change, 0.079 percentage points (Fig. S3E). The interval from 1.9 to 3.9 Å contains about $6 \times 10^{-9}$ of the axial equilibrium probability in each of the reference magnesium profiles.

In the uniform-inset calculation, the removed 0.5 Å ring carries about 98.3% of the classical magnesium weight at the basin and 15.9% at the intracellular barrier. Excluding a fraction $f$ of the Boltzmann weight raises the classical free energy by $-k_{\mathrm{B}}T \ln(1-f)$. The increase is about 109 meV in the basin and 5 meV at the intracellular interface. The classical intracellular barrier decreases by about 105 meV, and the magnitude of its quantum correction decreases from 22.7 to 8.2 meV. The reference and inset results are shown with the whole-pore and entrance-only smoothing results (Fig. S3D, E).

We asked how much of a passage estimate depends on positions inside the smoothed interval. The mean first-passage time contains two nested integrals [19]. The outer position $x$ lies between the basin and the absorbing interface. The inner position $y$ lies between $x$ and the reflecting interface. Each pair carries weight $\exp(\beta[W_X(x) - W_X(y)])$, so pairs that combine high outer free energy with low inner free energy contribute most. We grouped position pairs by membership in the entrance interval. The groups contain pairs with $x$ inside, pairs with only $y$ inside, and pairs with both positions outside. We inserted the interval endpoints into the piecewise-linear axial profiles and integrated the exponential weights on each segment. The fractions from the first two groups together give the share of the passage integral associated with either position inside the interval. The reference magnesium fractions sum to at most 0.117% across both profiles and directions. For passage to the extracellular side, every outer position lies between the basin and that interface, so the entrance contributes only through the inner integral (Table S4).

**Table S4:** Fractions of the reference mean first-passage integrals associated with the entrance interval from 1.9 to 3.9 Å at 310 K. The two columns for each direction are disjoint. The inner-only fraction includes pairs whose outer position lies outside the interval.

| | | Intracellular | | Extracellular | |
|---|---|---|---|---|---|
| Ion | Profile | Outer | Inner only | Outer | Inner only |
| $Na^{+}$ | Classical | $3.23 \times 10^{-2}$ | $9.73 \times 10^{-6}$ | 0 | $1.01 \times 10^{-5}$ |
| $Na^{+}$ | Quantum | $3.25 \times 10^{-2}$ | $7.91 \times 10^{-6}$ | 0 | $8.19 \times 10^{-6}$ |
| $Mg^{2+}$ | Classical | $1.10 \times 10^{-3}$ | $6.32 \times 10^{-9}$ | 0 | $6.33 \times 10^{-9}$ |
| $Mg^{2+}$ | Quantum | $1.17 \times 10^{-3}$ | $6.17 \times 10^{-9}$ | 0 | $6.18 \times 10^{-9}$ |
| $Ca^{2+}$ | Classical | $1.10 \times 10^{-3}$ | $6.32 \times 10^{-9}$ | 0 | $6.33 \times 10^{-9}$ |
| $Ca^{2+}$ | Quantum | $1.16 \times 10^{-3}$ | $6.38 \times 10^{-9}$ | 0 | $6.39 \times 10^{-9}$ |

To test the reflecting boundary, we moved it inward by 1, 2, or 3 Å and calculated passage times on the shortened interval with the same absorbing interface and axial free-energy profiles. Across all three ions and both directions, the MFPT percentage differs by at most 0.000184 percentage points from the value with reflection at the membrane interface (Fig. S3F). Every shifted reflecting boundary lies within the membrane slab. A bulk-to-bulk calculation would require a finely resolved potential and solvent-accessible geometry through both vestibules.

#### 5 Matched-grid refinement of the electrostatic field

We tested how the spacing of the electrostatic grid affects the results for a fixed description of the protein and membrane. We held the parent potential, physical box, ion accessibility, directional permittivities, and volumetric charge density fixed across the sequence. We reconstructed each coefficient as the trilinear interpolant of its original 0.25 Å map. We used trilinear interpolation of the pore-parent potential to set Dirichlet values on all faces of the $24 \times 24 \times 40$ Å$^3$ box. We used the same spacing in all three coordinate directions, with values of 0.50, 0.25, and 0.125 Å. We reserved a 1/6 Å grid to test the dependence on spacing inferred from the other grids. This construction isolates the effect of discretizing a common continuous electrostatic problem. The separate focused-field comparisons also capture changes associated with generating the coefficient maps on different grids.

We independently discretized the nonlinear Poisson–Boltzmann equation (Eq. 1) with centered face fluxes for the dielectric term and nodal values for the fixed-charge and mobile-ion terms. We evaluated the three permittivities at the corresponding face centers. At each interior node, we divided the absolute residual by the corresponding diagonal entry of the Jacobian. The solver stopped when the maximum of these ratios fell below $10^{-10}$ in units of the dimensionless potential $u = e\phi/(k_B T_{PB})$. The 0.25 Å solve reproduced the original field with a root-mean-square difference of $1.324 \times 10^{-4}$ $k_B T_{PB}/e$ over the box nodes. Tightening the finest-grid criterion to $10^{-12}$ changed the potential by at most $6.978 \times 10^{-11}$ $k_B T_{PB}/e$. This check places the iterative change below the differences among grids.

We compared potential values on common nodes at 0.50 Å intervals, so the error comparison requires no spatial interpolation of the solved fields. The centerline sample includes 61 positions. Of the 10,733 nodes within the accessible cross-sections, 6,269 satisfy the hydrated-core criteria of at least 1 Å clearance from the atomic spheres and at least 95% of bulk-water permittivity in all three directions. We denote the root-mean-square differences between the coarse and medium grids and between the medium and fine grids by $D_{CM}$ and $D_{MF}$, respectively. The effective order $p_{\text{eff}} = \log_2(D_{CM}/D_{MF})$ describes how strongly the two differences decrease. An asymptotic model $u_h = u_* + ah^p$ also requires the signed difference fields to be proportional and to predict the difference at the held-out spacing [20]. We tested this by multiplying the medium-minus-fine difference field by $(h_M^{p_{\text{eff}}} - h_V^{p_{\text{eff}}})/(h_M^{p_{\text{eff}}} - h_F^{p_{\text{eff}}})$, where $h_V = 1/6$ Å, and comparing it with the measured medium-minus-intermediate difference.

On the centerline, the medium-minus-fine and medium-minus-held-out differences are both smaller than the coarse-minus-medium difference and vary differently along the axis (Fig. S4A). In the hydrated core, the root-mean-square difference decreases from 4.81 to 0.0683 $k_B T_{PB}/e$ across the two halving pairs (Fig. S4B). The fitted model predicts a core difference of 0.0635 $k_B T_{PB}/e$ for the held-out pair, which gives 0.500 $k_B T_{PB}/e$ when solved directly. The prediction error has the same scale as the measured difference (Table S5). We therefore use the effective orders to characterize the halving comparisons and report the observed medium-to-fine differences to describe the sensitivity of the pore field. On a unit cube with relative permittivity 2, a unit coefficient for the nonlinear term, and exact potential $u = 0.4 \sin(\pi x) \sin(\pi y) \sin(\pi z)$, the solver gave successive root-mean-square orders of 1.94 and 1.97 on 17, 33, and 65 nodes per axis (Fig. S4C). This manufactured test verifies the expected second-order behavior for smooth inputs.

The fixed-charge data explain much of the large coarse-grid difference. Trapezoidal integration of the sampled charge over the box gives -8.87 and -4.97 $e$ at 0.50 and 0.25 Å, respectively. The 0.125 Å grid gives -4.97 $e$. Its nodes subdivide each trilinear cell of the original charge map, and the trapezoidal integral is exact for that interpolant on both grids. The 1/6 Å sample gives -5.26 $e$. When we replaced point sampling by exact averages of the same charge interpolant over the 0.50 Å dual cells, the coarse-to-medium core difference decreased to 0.044 $k_B T_{PB}/e$. The charge-averaging comparison identifies the contribution of coarse-grid charge sampling to the reduction between the halving pairs. Across the full accessible region, the medium-to-fine maximum difference is 1.70 $k_B T_{PB}/e$, which also identifies a larger residual sensitivity

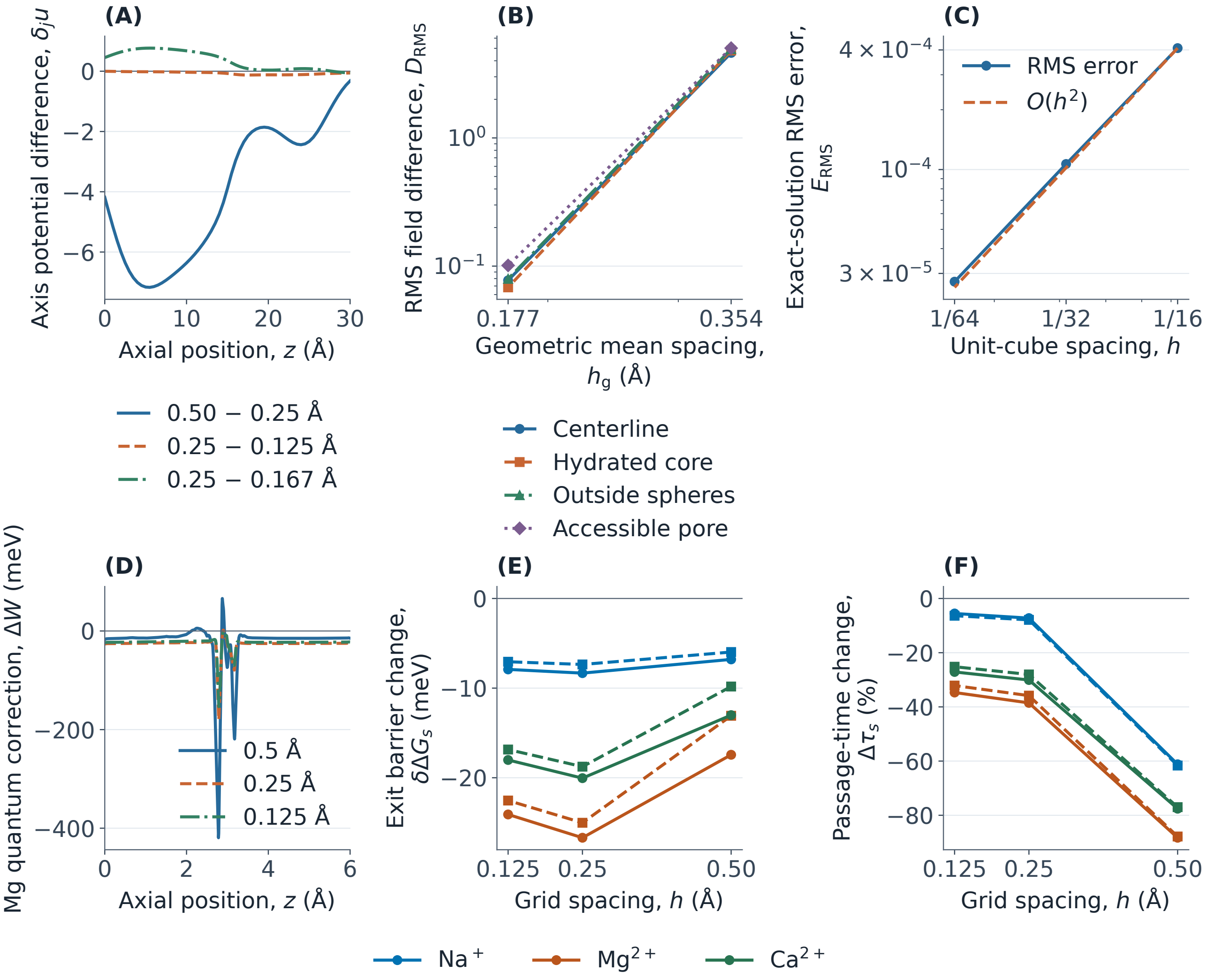


**Figure S4:** Matched-grid refinement at fixed field coefficients and parent boundary values. (A) Signed axial potential differences $\delta_j u(0,0,z)$ for adjacent and held-out grid pairs. (B) Root-mean-square field differences $D_{\mathrm{RMS}}$ against the geometric mean spacing $h_{\mathrm{g}}$ of each pair. (C) Root-mean-square error $E_{\mathrm{RMS}}$ versus grid spacing $h$ for the exact smooth solution on a unit cube. (D) Entrance-region magnesium correction $\Delta W(z)$. (E) Barrier correction $\delta\Delta G_s$ and (F) passage-time change $\Delta\tau_s$ as functions of the electrostatic mesh spacing $h$. The thermal panels use the circular point-ion reference. Solid lines indicate the intracellular target interface and dashed lines the extracellular target interface.

outside the hydrated core.

We used the three halving fields in the circular point-ion thermal and passage models at 310 K with identical hard-wall geometry and the reference axial Hamiltonian mesh. The 505 trace-correction planes and 1,291 classical integration planes included the tabulated radius positions and the finest electrostatic grid planes. The magnesium correction near the intracellular entrance is compared on the same axial interval (Fig. S4D). Across all three ions, the medium-to-fine change in the signed local extremum is at most 26.96 meV. The corresponding changes in the quantum barrier corrections reach 2.60 meV (Fig. S4E). The barrier corrections change nonmonotonically over the three grids. For magnesium, the medium and fine grids give passage-time changes of −38.52% and −34.69% toward the intracellular interface, and −35.85% and −32.13% toward the extracellular interface. The largest change in a first-passage percentage across the two finer grids is 3.83 percentage points (Fig. S4F). At both finer spacings, quantization lowers the two exit barriers and shortens the modeled passage times for all three ions. We report the directly measured medium-to-fine changes as resolution indicators for the fixed coefficients and protonation microstate.

**Table S5:** Field differences on common nodes. Each difference is a root-mean-square potential difference in $k_B T_{PB}/e$. Cosine similarity is the normalized dot product of the signed difference vectors for the two halving pairs. The held-out column gives the ratio of prediction error to the measured difference for the 1/6 Å grid.

| Region | $D_{CM}$ | $D_{MF}$ | $p_{eff}$ | Cosine similarity | Held-out ratio |
|---|---|---|---|---|---|
| Centerline | 4.605 | 0.077 | 5.894 | 0.513 | 1.059 |
| Hydrated core | 4.806 | 0.068 | 6.137 | 0.503 | 1.050 |
| Outside atomic spheres | 4.931 | 0.080 | 5.954 | 0.440 | 1.048 |
| Accessible cross-section | 5.019 | 0.101 | 5.632 | 0.334 | 1.040 |

### 6 Computational settings and quantitative comparisons

The following numerical settings and quantitative results specify the calculations discussed in the main text. The descriptions retain the physical references, comparison conditions and numerical values for each model.

#### 6.1 Computational sampling and resolution

For the magnesium geometry control, we construct a centered circular wall that clears every positive-radius PARSE sphere in the electrostatic model. We combine the structural radius knots with a uniform mesh spaced by at most 0.01 Å to enforce clearance between samples. For each interval between adjacent knots, a sphere has its largest cross-section on the plane closest to its center. We subtract the radius of that cross-section from the distance between the center and the pore axis. The smallest of these clearances across the intersecting spheres sets the upper bound on a constant radius that avoids sphere overlap throughout the interval. We cap both endpoints by that radius, with a numerical clearance of $10^{-8}$ Å, so linear interpolation also clears the spheres between knots. A further reduction where needed bounds the magnitude of the slope by that of the steepest segment of the structural profile. The resulting disk can be smaller than the full irregular sphere-free cross-section. The irregular-domain calculation retains the remaining accessible space around the axis.

The shortest clearance from a center to the atomic-sphere surfaces is a 1-Lipschitz function of position. We bounded its maximum over each structural disk using a square grid spaced by 0.035 Å. The maximum at the nodes inside the disk is a lower bound. The maximum at all nodes within one half-diagonal of the disk plus one half-diagonal is an upper bound.

We sampled the regularized dielectric Green function in four boxes. The first three have lateral half-widths of 8, 10, and 12 Å and extend from $-4$ to 36 Å along the pore axis. The fourth retains the 12 Å half-width and moves the axial faces to $-7.5$ and 37.5 Å. Each source carried charge $+2e$ in a Gaussian of standard deviation 0.38 Å, truncated at three standard deviations and normalized on the mesh. We solved the dielectric equation with zero Dirichlet values on all six faces. The same source, mesh, and boundary conditions define the homogeneous-water charging energy. We invert the homogeneous-water finite-difference operator in a separable discrete sine basis. A direct sparse solve on a small three-dimensional box verifies the evaluation. The background molecular potential is averaged over the truncated Gaussian for the thermal calculation. The dielectric charging calculation concerns polarization of the fixed medium. The nonlinear Poisson–Boltzmann background retains reservoir screening.

We use the variational energies in main-text Eq. 2 to supply the short-range spatial variation. At each calibration probe we form $\theta = (S_\sigma - S_L)/(S_U - S_L)$ and clip it to $[0, 1]$. A local thin-plate spline interpolates $\theta$ using 35 neighboring training values and coordinates scaled by factors of 1, 1, and 0.62 along $x$, $y$, and $z$. The primary spline smoothing parameter is $10^{-3}$. We also use $2.5 \times 10^{-2}$ to test sensitivity to the smoothness of the interpolant. The thermal solver samples the resulting field $S_L + \theta(S_U - S_L)$ by trilinear interpolation of its values on the 0.25 Å mesh.

We approximate the rotation integral using 256 scrambled Sobol rotations. A Cartesian field with 0.20 Å spacing represents the minimum sphere clearance at its nodes. We interpolate the clearance at each rotated oxygen position. Interpolation and rotational quadrature were tested at 14 representative ion centers using direct sphere distances and independent 1024-orientation calculations. We denote the corresponding free-energy errors by $e_{\mathrm{grid}}$ and $e_{\mathrm{orient}}$. The grid error is the steric-grid estimate minus the direct-distance estimate at the common 256-rotation rule. The orientation error is the 256-rotation direct estimate minus the mean of the 1024-rotation direct estimates. The mean accessible water-site count is $\langle n_{\mathrm{open},i} \rangle$. The signed held-out self-energy error is $e_S = S_\sigma^{\mathrm{model}} - S_\sigma^{\mathrm{direct}}$. The maximum changes are 9.3 and 16.3 meV, respectively. A second full shell map uses a 0.125 Å clearance grid with 512 orientations and is propagated through the thermal calculation. In a matched independent-ligand control, we first average the single-site access factor over all rotations and then take its $N_i$th power. This keeps the same steric criterion, single-site access, penalty, and dielectric field.

For the separate control with independent ligand directions, let $f_i(\mathbf{r})$ be the fraction of 110 approximately uniform antipodal directions for which an oxygen center, placed at the representative first-shell distance from $\mathbf{r}$, lies outside the prepared atomic spheres. We evaluate this fraction on a 0.25 Å mesh and interpolate to the thermal positions (main-text Fig. 2D).

For the atomic-union and finite-ion calculations, each thermal plane retains the part of the structural disk outside the union of all positive-radius spheres of the prepared charge-and-radius model (main-text Fig. 2A, B). An ion-center clearance $a_{\mathrm{c}}$ is implemented by expanding each three-dimensional sphere radius by $a_{\mathrm{c}}$ before intersecting it with the plane. The five-point transverse kinetic stencil sets the wave function to zero at the first encountered boundary along an edge. It severs a link if a sphere intersects the edge between accessible nodes. For boundary distances below one fifth of the mesh spacing we use the same cutoff convention as for the original disk. For classical integration we subtract the exact excluded intervals on each azimuthal ray and apply Gauss–Legendre quadrature in squared radius on each remaining interval. The primary comparisons use 32 radial nodes per interval and 320 azimuths. At six representative planes a 96-by-1280 check changes the classical integrals, expressed as free energies, by at most 0.453 meV for the atomic-union calculation and 0.002 meV for the revised calculation. The transverse quantum trace uses the deterministic Chebyshev solver. Its local ratio to the classical lattice sum is calibrated by the infinite free square-lattice trace per node before applying the continuum classical integral. This calibration uses the same discrete kinetic operator for all irregular domains and preserves a comparison across meshes.

We estimate the first and second derivatives of the effective axial potential $V(z) = F_{\perp,i}^{\mathrm{Q}}(z)$ at interior axial positions from a local quartic fit over 0.8 Å.

To estimate how a second ion of the same species affects escape, we combine two copies of the classical transverse free-energy profile with a screened Coulomb repulsion $v = Z_i^2 e^2 \exp(-\rho/\lambda_{\mathrm{D}})/(4\pi\varepsilon_0\varepsilon\rho)$. The effective separation is $\rho = [(z_1 - z_2)^2 + a^2]^{1/2}$, with transverse offset $a = 2$ Å and screening length $\lambda_{\mathrm{D}} = 8.01$ Å. We evaluate this prescribed interaction at 310 K with relative permittivities $\varepsilon = 20$, 40, and 78.5. The screening length is the bulk Debye value for 150 mM electrolyte at $\varepsilon = 78.5$ and remains fixed across the three permittivities. This comparison varies the strength of the prescribed pair repulsion at a common screening range. The axial quadrature uses a 0.0125 Å interval and a 0.025 Å comparison. The classical partition function for two indistinguishable ions is $Z_2 = \frac{1}{2} Z_1^2 \langle \exp(-\beta v) \rangle_{\mathrm{C}}$. Here $Z_1$ is the single-ion partition function, and the average uses two independent draws from the classical single-ion Boltzmann distribution. The interaction raises the pair free energy by $\Delta F_{\mathrm{pair}} = -k_{\mathrm{B}} T \ln \langle \exp(-\beta v) \rangle_{\mathrm{C}}$ relative to the same two-ion system with $v = 0$. Integrating over one ion's position gives the marginal free-energy profile whose barriers describe escape for its partner. The reported pair barriers use the local marginal minimum within the Y1085–E1047 reference window, consistent with the circular single-ion comparison.

We repeated the two-ion benchmark on the directly calculated classical transverse profiles. Both ions follow independent axial imaginary-time Brownian bridges in the kinetic reference; their potential along a path is the sum of the two single-ion profiles and the screened repulsion used in the classical two-ion benchmark. For each thermal trace, we drew the path roots from the corresponding classical Boltzmann distribution on a midpoint grid. The quantum-to-classical partition-function ratio, after removal of the free kinetic prefactor, is the classical-root average of the bridge weight $\exp[-\beta(\overline{U} - U_{\mathrm{root}})]$, where $\overline{U}$ is the discretized imaginary-time average and paths with an axial bead outside the slab receive zero weight [21]. In the ratio of the interacting trace to the square of the single-ion trace, the kinetic prefactors cancel. We performed eight independent scrambled Sobol integrations with $2^{14}$ bridges each and compared 32 and 64 time slices at root spacings of 0.025 and 0.0125 Å. We used 128 time slices as an additional check for magnesium at the finer root spacing.

We solved the coupled problem over 1.35–4.40 Å and examined the interior interval 2.35–3.40 Å. The plane quantum corrections are referenced to the grid point nearest 2.45 Å.

For the axis-calibrated control, we obtained $\theta(z)$ at 16 axial sites by matching independent dielectric-only finite-difference solves at 0.25 Å spacing and interpolated it with a shape-preserving cubic.

We estimated the smooth electrostatic contribution from the squared-gradient term of the local Wigner–Kirkwood expansion [21–23] at 1,001 centerline positions from 0.4 Å inside the intracellular interface to 0.4 Å inside the extracellular interface.

The mass parameters use tabulated atomic masses as the ion-mass approximation. We used tabulated isotope masses [24] and repeated the calculation with axial spacings of 0.0125 and 0.00625 Å. The isotope passage ratios use analytic integration of the piecewise-linear axial profiles, with identical basin positions and integration rules at the two masses.

Of the 409 direct probe positions in each box, 343 supply the fit at 49 axial planes. At each plane we used the centerline and two radial fractions, 0.52 and 0.96, along each of three azimuthal rays within one quadrant. The radial unit is the distance from the axis to the first excluded position at 1.15 Å sphere clearance or the structural disk boundary. The training set uses the fourfold rotations of these positions. An additional 48 probes at distinct azimuths and depths, 13 near low-energy positions of the axis-calibrated model, and five probes at rotated counterparts of training sites were reserved for comparison. All Gaussian supports clear the prepared atomic spheres.

#### 6.2 Dielectric and hydration calibration

The dielectric and shell calibration resolves the signed error at held-out probes (Fig. S5A), mesh and domain changes at selected probes (Fig. S5B), the mean accessible water-site count along the axis (Fig. S5C), and the separate steric-grid and orientation-quadrature errors (Fig. S5D).

#### 6.3 Water coordination and occupied wells

At the narrowest tested plane ($z = 15.497$ Å), the maximal ion-center clearance lies between 1.442 and 1.466 Å (main-text Fig. 2C). This clearance lies below both representative bulk ion–oxygen distances. A spherical envelope that extends to the first-shell oxygen radius therefore has no continuous path through the full modeled slab for either divalent ion. A discrete shell can rotate and retain a subset of its waters as it enters a constriction.

At the centerline of the narrowest structural section ($z = 15.497$ Å), the mean counts of geometrically accessible sites are 0.62 for magnesium and 1.09 for calcium with the 1.40 Å water probe. The correlated free energies there are 465 and 646 meV under the primary penalty. Average site counts use the sharp distance criterion, and the energy calculation uses the smooth access factor. Both 1024-orientation samples give zero fully accessible orientations at this site for either geometry. This observation concerns the centerline and the specified coordination geometries.

At the point-ion magnesium basin plane ($z = 17.281$ Å), the refined probe benchmark gives 430 meV on the pore axis and 743 meV at an interior off-axis position on the 0.125 Å mesh. The largest box used for the spatial calculation gives 508 meV on the axis at the same plane. These charging energies exceed the 21–23 meV quantum barrier shifts of the bare magnesium reference. Their spatial variation changes which positions contribute to the transverse Boltzmann weights.

The primary dielectric box has lateral half-width $b = 12$ Å and extends from $-7.5$ to 37.5 Å along the axis. The primary passage values use the finest transverse spacing, $h_\perp = 0.050$ Å. The deepest classical wells lie at 22.75 Å for magnesium and 11.00 Å for calcium. Their intracellular and extracellular classical barriers are 698 and 290 meV for magnesium and 671 and 669 meV for calcium (main-text Fig. 3B). The corresponding passage-time changes from those wells are 0.68% and $-0.66$% for magnesium and $-0.10$% and 16.12% for calcium. The extracellular route from the calcium well crosses the constriction, where the

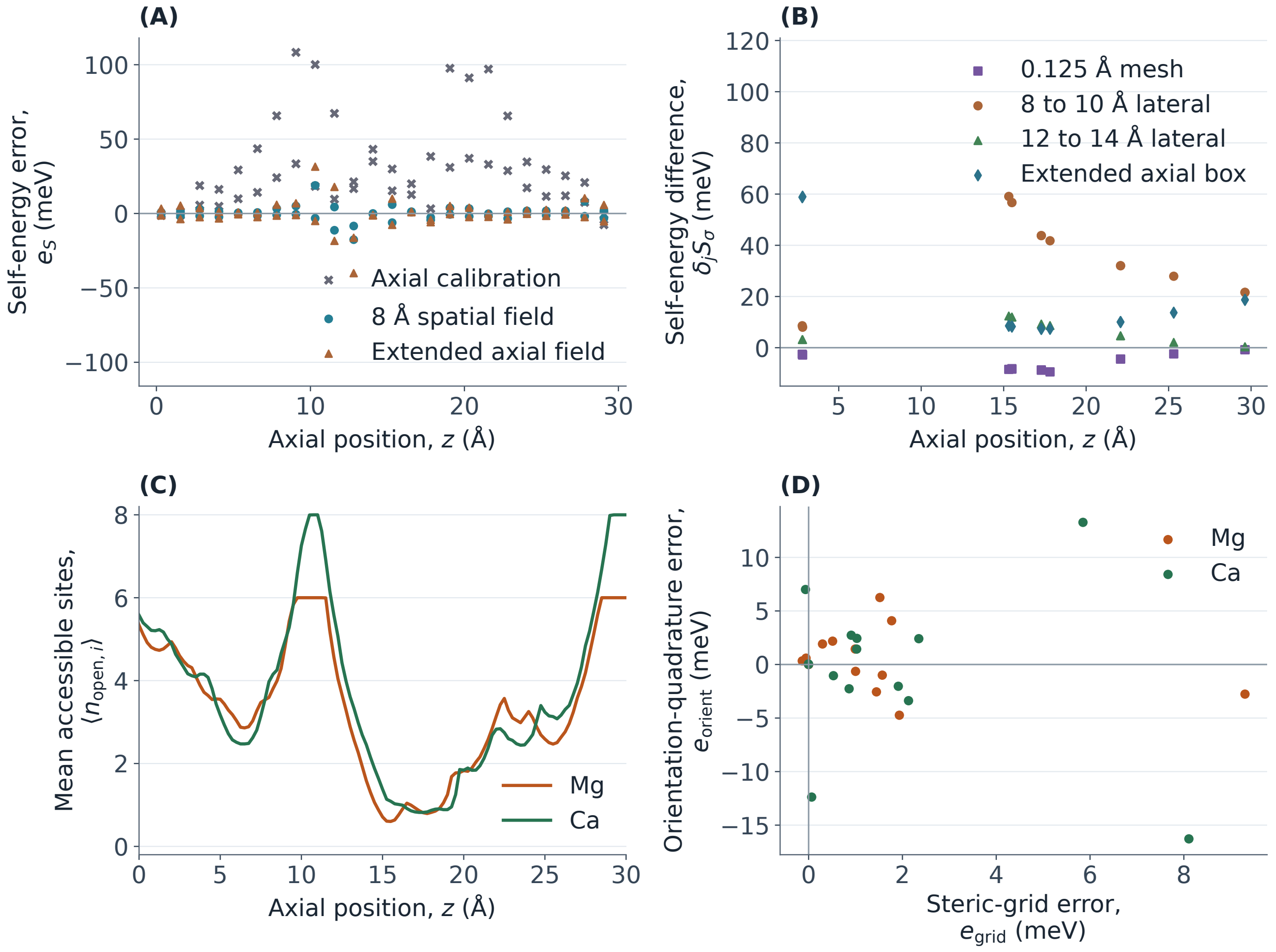


**Figure S5:** Numerical checks of dielectric insertion and shell access. (A) Signed self-energy error $e_S$ along $z$ at 48 held-out probes. Axis-calibrated and 8 Å spatial estimates use the same boundaries. The extended axial field uses its own direct probes. (B) Self-energy difference $\delta_j S_\sigma$ for the indicated mesh and box comparisons at 13 probes. The 0.125 Å mesh and 10 Å box are compared with the 8 Å calculation. The 14 Å and extended boxes are compared with the standard 12 Å calculation. (C) Mean accessible water-site count $\langle n_{\mathrm{open},i} \rangle$ along the axis. (D) Steric-grid and orientation-quadrature errors $e_{\mathrm{grid}}$ and $e_{\mathrm{orient}}$.

quantum free-energy profile raises the exit barrier by 3.30 meV. The preferred magnesium well lies beyond the constriction on the extracellular side (main-text Fig. 3C).

The deepest magnesium well in the 8 Å box lies on the intracellular side of the constriction. At 12 Å lateral half-width it lies on the extracellular side. At common transverse spacing, extending the axis of the 12 Å box changes the directional passage effects by up to 6.27 percentage points for magnesium and 0.02 percentage points for calcium (main-text Fig. 3D).

At 0.085 Å transverse spacing in the extended box, the matched independent-ligand calcium control favors the well at 22.08 Å and gives an extracellular passage change of −0.45%. A common shell rotation places the well at 11.00 Å and increases the extracellular passage time by 13.14%. The comparison uses the same dielectric field, ligand count, access criterion, and obstruction penalty. Within the correlated model, reducing the penalty from $4k_{\mathrm{B}}T$ to $2k_{\mathrm{B}}T$ moves the calcium well downstream and gives an extracellular passage change of −0.28% (main-text Table 2).

To resolve regional retention, we also start passage from local classical wells in the same insertion landscape. In the extended axial box the Y1085–E1047 minima are at 22.75 Å for magnesium and 22.75 Å for calcium. Their energies above the respective global minima are 0.0 and 42.4 meV. Starting from the window minimum changes the magnesium times by 0.68% and −0.66%, and the calcium times by 1.83% and −0.01%. The lowest magnesium well between $z = 6$ and 13.59 Å is at 11.25 Å, 69.2 meV above the global minimum. From that upstream well, the extracellular passage change is 23.35%.

We also averaged the two pointwise passage profiles over a shared classical equilibrium starting distribution. This gives percentage changes of 0.63% and 23.33% for magnesium and 0.35% and 16.12% for calcium. For magnesium, positions with $z \leq 15.50$ Å carry 5.2% of the classical weight and 99.89% of the averaged classical extracellular time. The long extracellular escape time from the upstream side accounts for its large contribution to the average.

For intracellular passage of magnesium, the logarithm of the time ratio is 0.007. Its contributions are -0.030 from the barrier change, 0.011 from the basin weight, 0.027 from the exit width, and -0.001 from the nested integration limits. The positive contributions from the basin and exit weights exceed the contribution of the barrier decrease and leave the overall time ratio above one.

##### 6.4 Atomic access and dielectric sensitivity

Seven off-axis sites with at least 1.15 Å clearance and seven with smaller clearance were withheld from calibration. The root-mean-square errors were 98 and 1209 meV, respectively (main-text Fig. 4A, B). The two largest discrepancies occurred at low-clearance sites. At the basin plane an interior probe gives 743 meV on the 0.125 Å mesh. A nearby wall probe gives 4234 meV at that spacing, and 91.4% of its Gaussian weight lies outside the atomic spheres. The centerline value at the structural throat is 378 meV on the same refined mesh. These comparisons quantify the loss of accuracy as the Gaussian approaches the protein boundary. The guarded control restricts centers to at least 1.15 Å clearance. Its electrostatic uncertainties still exceed individual quantum free-energy changes at some sites.

Cutting out the atomic spheres reduces the magnitude of the local magnesium peak to 133.5 meV. Its passage-time changes remain −37.36% and −34.35% toward the intracellular and extracellular interfaces. A further 0.5 Å clearance around the spheres gives a different perturbation from a uniform 0.5 Å inset of the circular wall. Adding the reaction field at fixed 0.5 Å atomic clearance moves the classical magnesium basin from 17.28 to 22.08 Å and reduces the classical intracellular barrier from 1204.95 to 723.24 meV. With independent shell directions and 1.15 Å clearance, calcium gives passage-time changes of 0.03% and −0.35% (main-text Fig. 4C, D).

The axis-calibrated estimate raises low-energy positions in the original basin and alters the classical barriers. We compare several controls, including the variational energies propagated independently, the Gaussian background average, and the shell-access penalty (main-text Table 3). On the finer transverse mesh, the axis-calibrated magnesium control with Gaussian background changes the predicted time by 0.06% and −0.63%. The lower and upper variational fields give different changes, and the intracellular change can have either sign. The two trial fields give model sensitivity cases for passage because the passage integral depends on energy differences along the entire profile. In the upper-energy case, an additional classical well lies at 11.25 Å and is 65.31 meV below the minimum in the reference basin window. Starting at this global minimum gives changes of −0.09% and 6.28%. The axis-calibrated comparisons place the global classical minima inside the reference basin window. The passage comparison depends on spatial differences in energy, so the bounds on charging energy define sensitivity cases for escape.

In the 8 Å half-width box, the spatial estimate reduces the root-mean-square discrepancy at the 48 off-axis held-out sites from 41.6 to 5.0 meV, with a maximum discrepancy of 18.9 meV. The corresponding root-mean-square errors are 1.3 and 2.7 meV for the low-energy and rotated probes. These comparisons use the interpolated field supplied to the thermal calculation. At 13 selected sites, refining the dielectric mesh to 0.125 Å changes the direct self-energies by at most 9.4 meV. Increasing the half-width to 10 Å changes those energies by up to 59.1 meV. The span of the changes across those sites is 51.1 meV and measures the part that varies with position. Extending the 12 Å box to a lateral half-width of 14 Å gives changes spanning 12.0 meV. Extending its axial range gives changes spanning 51.652 meV (Fig. S5A, B).

The standard and axially extended 12 Å boxes give root-mean-square errors of 9.1 and 9.4 meV across the same 48 held-out positions, and 1.8 and 1.9 meV at the 13 low-energy validation positions. We separately selected 20 low-energy ion centers in each 12 Å box from the full $Mg^{2+}$ and $Ca^{2+}$ insertion energies. The spatial estimates there have root-mean-square discrepancies of 5.5 and 5.6 meV, with maximum discrepancies of 11.2 and 11.6 meV. Each full spatial field was propagated through the thermal and passage calculations. These interpolation comparisons share the dielectric boundary within each box. The independent comparisons across boxes measure the response to changing that boundary.

In the extended axial box, reducing the transverse spacing from 0.085 to 0.050 Å changes the passage effects by up to 0.17 percentage points for magnesium and 2.97 for calcium. The refined shell grid and orientation rule at 0.085 Å change them by up to 0.02 and 0.50 percentage points. At the same transverse spacing, changing the lateral half-width from 8 to 12 Å shifts them by up to 20.46 and 0.28 percentage points, and extending the axial box shifts them by up to 6.27 and 0.02 percentage points. For the 8 Å magnesium case at $h_\perp = 0.065$ Å, reducing the maximum axial sampling interval from 0.115 to 0.055 Å changes the passage percentages by up to 0.10 percentage points.

Across the three ligand penalties in the extended axial box at 0.085 Å, the intracellular changes range from $-0.35\%$ to 21.47% for magnesium. The calcium extracellular changes range from $-0.28\%$ to 14.74%. At the reference penalty, reducing the water probe to 0.70 Å moves the lowest calcium well downstream and gives an extracellular change of $-0.32\%$. The seven-ligand geometry with the 1.40 Å probe retains the intracellular-side calcium well and gives 12.74% (main-text Tables 1 and 2).

#### 6.5 Point-ion thermodynamics and passage

A widening gives the ion a larger area and can expose new electrostatic environments. Both effects change the transverse partition function, so the preferred axial position depends on the whole cross-section. At 310 K, the classical free-energy minimum is near $z = 17.28$ Å, on the intracellular side of the centerline potential minimum. The positions differ because the transverse Boltzmann integral depends on the area of each slice and the mean Boltzmann factor over that area.

From the interior minimum, an ion encounters a free-energy rise on the way to either membrane interface. The classical barriers toward the intracellular and extracellular interfaces are approximately 590 and 397 meV for sodium (22 and 15 $k_\mathrm{B}T$) and 1252 and 877 meV for either divalent ion (47 and 33 $k_\mathrm{B}T$). The highest point along the intracellular route lies at the interface for all three ions. On the extracellular route it lies at the interface for sodium and near $z = 29.63$ Å for magnesium and calcium. The classical and quantum profiles place the interior minimum and exit maxima in the same regions of the membrane slab (main-text Fig. 5A). These are confined escape barriers of the fixed test-charge model. A bulk-to-bulk permeation profile would include the energy of ion insertion, the vestibules, and occupancy by mobile ions [25–27]. The barrier heights represent a large thermal penalty for an ion to leave its preferred site in the bare-charge model. Dielectric charging and water coordination change the landscape on which that retention occurs.

The local effect of quantization is greatest near the intracellular entrance, where the free energy changes over a short distance. At 310 K the thermal energy $k_\mathrm{B}T$ is 26.71 meV. Near $z \simeq 2.79$ Å, the quantum-minus-classical free-energy differences are $-42.4$ meV for sodium, $-179.3$ meV for magnesium, and $-141.3$ meV for calcium. For magnesium, motion along the pore contributes $-184.3$ meV, and motion across the pore offsets part of that change with a contribution of $+5.0$ meV. Calcium likewise gives an axial contribution of $-144.4$ meV and a transverse contribution of $+3.1$ meV. For sodium, the axial contribution is $-42.4$ meV and the transverse contribution is about $-0.1$ meV. At that entrance peak, the smooth-potential estimates for the axial component are $-5.8$, $-21.7$, and $-14.9$ meV for sodium, magnesium, and calcium, respectively (main-text Eq. 11, main-text Fig. 5B). For magnesium, the negative entrance extremum is followed by a positive local maximum of 4.1 meV at $z = 2.89$ Å. Smaller variations occur farther down the pore between 24 and 28 Å. Smoothing the radius near the entrance moves the largest magnesium correction to $z = 23.96$ Å. The sharp widening therefore controls where the largest local correction occurs, while the slow steps of escape lie elsewhere in the pore (Section 4).

The entrance widening exposes substantially lower-energy positions across neighboring slices. From $z = 2.781$ to 2.980 Å, the radius increases from 3.92 to 5.32 Å and the classical magnesium free energy falls by about 0.41 eV. The increase in area alone accounts for about 16 meV. Favorable positions close to the protein account for most of the lowering. Delocalization across a sharp change in the positions accessible to the ion explains the much larger axial correction.

Centered differences in all three Cartesian directions gave maximum magnitudes of 0.033, 0.123, and 0.075 meV for sodium, magnesium, and calcium at 310 K. Increasing the difference half-step from 0.10 to 0.40 Å changed the magnesium maximum by less than 0.001 meV. This is a gradient-term diagnostic on the pore axis. The effective axial potential also depends on the area and energy distribution across the section, including regions where the permittivity changes. The centerline diagnostic therefore quantifies a different contribution from the axial Wigner–Kirkwood estimate (main-text Eq. 11). A scale estimate illustrates the difference. In the leading local expression for magnesium, a squared-gradient term of magnitude 184 meV would require a slope of about 4.3 eV per angstrom. A curvature term of the same signed magnitude would require about −344 eV per square angstrom. These are one-term scales calculated by inserting the size of the entrance axial correction into the smooth expansion, before choosing a reference. The axial correction itself is basin-referenced and resolves an abrupt change in accessible configurations. The one-term estimates give a scale for comparison with a smooth landscape.

In the circular point-ion reference, quantum delocalization shortens the modeled time to reach either membrane interface. Magnesium shows the largest effect on passage, with changes of −37.4% toward the intracellular side and −34.5% toward the extracellular side (main-text Fig. 6A). Its exit barriers change by −22.7 and −20.9 meV, much less than the largest local free-energy change at the entrance (main-text Fig. 6B, main-text Table 4). Along the intracellular route, the free-energy difference derived from the normalized densities, $-k_{\mathrm{B}}T\ln(P_{\mathrm{Q}}/P_{\mathrm{C}})$, is +9.0 meV at the basin minimum and −13.7 meV at the barrier maximum. The change in the barrier height is the difference between these two values, −22.7 meV.

The magnesium barrier decreases are 1.33 and 1.34 times the calcium decreases for the intracellular and extracellular routes, respectively. A barrier correction proportional to the inverse of ion mass would give $m_{\mathrm{Ca}}/m_{\mathrm{Mg}} = 1.65$ in both directions. The barrier correction combines free-energy changes at positions where the local curvature and confinement differ. The passage time also depends on the widths of these regions because the integrals include positions near the basin minimum and barrier maximum [19, 28]. For magnesium passage to the intracellular interface, the logarithm of the time ratio, $\ln R_{\tau,-} = -0.47$, differs from the barrier change in thermal units, $\beta\delta\Delta G_{-} = -0.85$, by about 0.38. Factoring the passage integral (main-text Eq. 9), with analytic integration of every piecewise-linear segment for both widths and the passage time, gives log contributions of +0.338 and +0.044 from the basin and exit widths. The extracellular exit-width contribution is +0.022. The magnitude of the nesting contribution is below $10^{-10}$ in both directions. Sections 3 and 4 give the buffer and reflecting-boundary comparisons. Most of the gap reflects the broader thermally occupied basin in the quantum profile.

At $z = 2.980$ Å, the prepared atomic spheres occupy 14.2% of the reference disk and contain 99.9% of the classical magnesium weight. Excluding the points inside these spheres raises the slice free energy by 186 meV (Section 4). The magnesium extremum lies on the preceding, narrower side of the widening. Its sensitivity to the wall follows from the change in the accessible positions of adjacent slices.

The sphere-excluding circle reduces the radius by up to 1.51 Å and reaches a minimum of 1.37 Å (main-text Fig. 7A). We use the same potential, temperature, ion mass, thermal grids, and passage boundaries as in the structural circle. The largest correction by magnitude is −69.0 meV at $z = 2.78$ Å (main-text Fig. 7B). In this geometry the barrier changes are −30.62 and −28.70 meV, and the passage-time changes are −36.83% and −36.15% toward the intracellular and extracellular interfaces, respectively (main-text Fig. 7C, D). The classical intracellular passage time decreases by a factor of 3.1 at the same diffusivity. The local peak and the relative changes in escape are therefore sensitive to different parts of the landscape. The centered disk tests sphere exclusion within the circular model. The irregular-wall comparisons retain off-axis space and resolve the effects of geometry, hydration, and dielectric charging (main-text Fig. 4).

The uniform 0.5 Å inset also restricts access in the occupied well. In the structural circle, the outer 0.5 Å carries 98.2% of the classical magnesium weight at the entrance peak and 98.3% at the basin. The inset gives passage-time changes of −8.9% and −8.6% toward the intracellular and extracellular interfaces, respectively (Section 4). Together the two wall comparisons show how access near the occupied site shapes the escape response. The circular inset is a geometry control for the point ion. The finite-ion model also accounts for dielectric charging and restricted water coordination.

The charge controls help explain why the local effect can be large even when the thermal wavelength is short relative to the pore diameter. With the electrostatic energy set to zero, the maximum free-energy correction in thermal units, $\max|\beta\Delta W|$, equals 0.050, 0.049, and 0.037 for sodium, magnesium, and calcium, respectively. The leading boundary term in the Dirichlet heat trace, the thermal sum over states whose wave functions vanish at the wall, estimates the local transverse confinement cost as $\Delta F_\perp/k_\mathrm{B}T \simeq \lambda_{\mathrm{th},i}/(2r)$ [29]. At fixed radius this estimate scales as $m_i^{-1/2}$. For magnesium it gives about 0.03–0.07 across radii of 1.4–3.0 Å. The total geometry-only correction also includes axial motion and is measured relative to the basin (main-text Fig. 8A).

When all three ions have charge $+e$, their peak magnitudes in thermal units are 1.588, 1.535, and 1.094, respectively. Sodium and magnesium give similar corrections and differ by about 6% in mass. Raising the test charge to $+2e$ makes favorable electrostatic positions more strongly preferred at equilibrium and increases the corrections for magnesium and calcium in this field. These two ions have identical classical position profiles because they share the domain and energy, and the mass factor from the momentum integral is constant along the pore. Magnesium's larger quantum correction then reflects its lower mass (main-text Fig. 8B).

Higher temperatures shorten the thermal de Broglie wavelength, the scale for quantum delocalization, and reduce the Boltzmann preference for low-energy positions. We vary temperature from 250 to 400 K with the test-charge energy held fixed. The peak correction in thermal units decreases from 3.054 to 0.659 for sodium, from 10.139 to 3.730 for magnesium, and from 8.402 to 2.721 for calcium (main-text Fig. 9A). For each ion, the larger of the two absolute percentage changes in passage time also decreases at every step in temperature (main-text Fig. 9B).

Magnesium's dimensionless peak decreases by a factor of 2.72. For a fixed, smooth energy surface, the dimensionless curvature term in the Wigner–Kirkwood expansion scales as $\beta^2$, which gives a factor of 2.56 over this temperature range. The term from the squared gradient scales as $\beta^3$. In the pore, temperature changes the transverse free energy itself, and both the smooth terms and the sharp entrance feature shape the peak response.

##### 6.6 Pore motion, coupled motion and pair comparisons

For a mode amplitude RMS of 1 Å, the entrance feature has an actual RMS displacement of 0.84 Å. The ensemble correction of greatest magnitude is −144.84 meV, and the ensemble mean of the signed individual extrema is −206.00 meV. The intracellular time changes are −38.20% and −37.35% in the fast and slow limits. The whole-pore mode of the same amplitude produces a different displacement at the entrance because of the endpoint constraints. Increasing the amplitude integration from 81 to 161 nodes changes the passage percentages by up to 0.001 percentage points and the ensemble extrema by up to 2.7 meV. A profile extremum carries greater quadrature sensitivity than the passage integral (main-text Fig. 10A, B).

At $z = 2.78$ Å on the refined disk plane, the scalar and independent-band corrections are −170.1 and −169.6 meV. The coupled correction at the same sample is 37.4 meV. For the atomic-union plane, at the scalar extremum ($z = 2.78$ Å), the scalar and coupled corrections are −78.3 and 33.4 meV. The opposing signs persist across three disk meshes and the orthogonal-plane test. The size of the effect depends on the boundary and potential. This localized plane calculation retains one transverse coordinate. It establishes a coupling effect for that problem and motivates a three-dimensional benchmark before assigning a corrected passage time.

In the screened pair benchmark, the classical interaction free energy is 44–112 meV for sodium and 159–299 meV for the divalent ions. The mean absolute axial separations are 3.2–6.0 and 5.9–6.6 Å, respectively. For a singly tracked ion in the pair, repulsion lowers the intracellular barrier by 45–116 meV for sodium and by 171–297 meV for the divalent ions. The corresponding extracellular reductions are 42–104 and 163–266 meV. The divalent intracellular barriers still lie between 0.95 and 1.08 eV. The benchmark therefore identifies the scale on which a second ion can alter escape from a well in the fixed continuum field. The reported barriers start at the local marginal minimum within the Y1085–E1047 window. For the divalent pairs, the global marginal minimum lies at 23.84 Å, beyond that window, and is 24–30 meV lower. From this global minimum the intracellular barriers are 0.98–1.10 eV and the extracellular barriers are 0.64–0.74 eV. The companion global-reference values are retained with the pair data.

Quantum motion increases the pair interaction free energy by 0.025–0.134 meV for sodium, 4.57–6.15 meV for magnesium, and 3.08–4.17 meV for calcium. The standard errors of the finest estimates, computed across the eight independent scrambles, range from 0.006 to 0.033 meV for the quantum interaction increment. These errors quantify integration uncertainty at fixed time slicing. At the common root spacing of 0.0125 Å, the magnesium estimates from 64 and 128 time slices differ by at most 0.05 meV. For a separate diagnostic, we hold the pair action at its root value as the single-ion potentials vary along the paths. Allowing the repulsion to vary along the quantum paths changes the increment by at most 0.05 meV. This comparison attributes most of the quantum increment to shifts in equilibrium occupation. For free identical ions at physical separations of at least 2 Å, the ratio of the exchange and direct thermal kernels is $\exp(-2\pi\rho^2/\lambda_{\mathrm{th}}^2)$ and lies below $10^{-250}$ for the three masses at 310 K. The thermal benchmark treats the axial coordinates as labeled and uses the prescribed repulsion. The slab-restricted paths provide a comparison of pair repulsion at fixed single-ion profiles.

On the fixed classical transverse profiles, the light-minus-heavy isotope shifts in the intracellular and extracellular barriers are −0.35 and −0.34 meV for $^{24}$Mg/$^{26}$Mg, −0.33 and −0.31 meV for $^{40}$Ca/$^{44}$Ca, and −0.46 and −0.41 meV for $^{6}$Li/$^{7}$Li. The largest change in the mean passage time is about 0.25% at common diffusivity. The larger local isotope differences, 6.6–9.8 meV, occur at positions whose contributions to the passage integral are small. For the three isotope pairs the leading $1/m_i$ scaling gives fractions of about 8%, 9%, and 14% of the light-isotope correction. For magnesium, a quantum barrier shift of 4 meV corresponds to a barrier-only isotope rate difference of about 1.2% at 310 K. A shift of 0.3 meV gives about 0.09%.

All three response calculations use zero perturbation at the faces of the same box. The box extends axially from −4 to 36 Å. On a 0.125 Å mesh with lateral half-width 8 Å, the dielectric self-energies at $z = 2.980$, 17.281, 22.980, and 29.630 Å are 150, 430, 277, and 76 meV, respectively, for the specified Gaussian probe. The screened pair energies for the six distinct pairs range from 7 to 520 meV. At a lateral half-width of 8 Å, refining the mesh from 0.25 to 0.125 Å changes the largest self-energy by about 9 meV. At 0.25 Å spacing, increasing the lateral half-width from 8 to 10 Å changes it by 44 meV. The maximum pair reciprocity error is below $10^{-7}$ meV.

At matched grids and boundaries, the 16-site calculation reproduces the four-site values within $10^{-7}$ meV. The extended matrix resolves the axial variation of dielectric charging and screened repulsion (main-text Table 5). We used the 110 distinct axial pairs separated by at least 2.28 Å for the interaction ranges, so their truncated Gaussian supports are disjoint. At the finer spacing, the five pairs separated by 2.50 Å span 340–656 meV. The pair response therefore depends on axially varying dielectric and screening properties as well as on separation. Reducing the mesh spacing at fixed box size changes the self-energies by up to 9 meV and the nonoverlapping pair energies by up to 15 meV. Moving the lateral faces at fixed mesh spacing changes those energies by up to 80 and 163 meV, respectively. Reciprocity errors across the three matrices are below $10^{-7}$ meV. The pair map describes on-axis linear response for the specified charge shape. The one-ion insertion landscape uses the spatial dielectric and hydration energy (main-text Eq. 1). The thermal two-ion benchmark uses the prescribed screened repulsion, which identifies the effect of repulsion at fixed single-ion profiles. A joint two-ion landscape across the pore would need off-axis pair response and joint coordination of the two hydration shells.

[2] M. A. Lomize, I. D. Pogozheva, H. Joo, H. I. Mosberg, and A. L. Lomize, OPM database and PPM

web server: resources for positioning of proteins in membranes, *Nucleic Acids Res.* **40**, D370–D376 (2012). doi:10.1093/nar/gkr703.
[3] A. Bondi, van der Waals volumes and radii, *J. Phys. Chem.* **68**, 441–451 (1964). doi:10.1021/j100785a001.
[4] O. S. Smart, J. G. Neduvelil, X. Wang, B. A. Wallace, and M. S. P. Sansom, HOLE: a program for the analysis of the pore dimensions of ion channel structural models, *J. Mol. Graphics* **14**, 354–360, 376 (1996). doi:10.1016/S0263-7855(97)00009-X.
[5] T. J. Dolinsky, P. Czodrowski, H. Li, J. E. Nielsen, J. H. Jensen, G. Klebe, and N. A. Baker, PDB2PQR: expanding and upgrading automated preparation of biomolecular structures for molecular simulations, *Nucleic Acids Res.* **35**, W522–W525 (2007). doi:10.1093/nar/gkm276.
[6] E. Jurrus, D. Engel, K. Star, et al., Improvements to the APBS biomolecular solvation software suite, *Protein Sci.* **27**, 112–128 (2018). doi:10.1002/pro.3280.
[7] Electrostatics Project, *PDB2PQR 3.7.1 release notes.* PDB2PQR release history.
[8] D. Sitkoff, K. A. Sharp, and B. Honig, Accurate calculation of hydration free energies using macroscopic solvent models, *J. Phys. Chem.* **98**, 1978–1988 (1994). doi:10.1021/j100058a043.
[9] M. H. M. Olsson, C. R. Søndergaard, M. Rostkowski, and J. H. Jensen, PROPKA3: Consistent treatment of internal and surface residues in empirical p$K_\mathrm{a}$ predictions, *J. Chem. Theory Comput.* **7**, 525–537 (2011). doi:10.1021/ct100578z.
[10] C. R. Søndergaard, M. H. M. Olsson, M. Rostkowski, and J. H. Jensen, Improved treatment of ligands and coupling effects in empirical calculation and rationalization of p$K_\mathrm{a}$ values, *J. Chem. Theory Comput.* **7**, 2284–2295 (2011). doi:10.1021/ct200133y.
[11] PROPKA developers, *PROPKA 3.5.1 release notes.* PROPKA releases.
[12] N. A. Baker, D. Sept, S. Joseph, M. J. Holst, and J. A. McCammon, Electrostatics of nanosystems: application to microtubules and the ribosome, *Proc. Natl. Acad. Sci. U.S.A.* **98**, 10037–10041 (2001). doi:10.1073/pnas.181342398.
[13] Electrostatics Project, *APBS 3.4.1 documentation.* APBS manual.
[14] Adaptive Poisson–Boltzmann Solver documentation, Charge discretization for multigrid calculations, APBS charge-mapping options.
[15] International Association for the Properties of Water and Steam, *Release on the Static Dielectric Constant of Ordinary Water Substance for Temperatures from 238 K to 873 K and Pressures up to 1000 MPa*, IAPWS R8-97 (1997). IAPWS R8-97.
[16] Adaptive Poisson–Boltzmann Solver documentation, Precomputed coefficient maps. APBS map interface.
[17] D. Gilbarg and N. S. Trudinger, *Elliptic Partial Differential Equations of Second Order*, 2nd ed. (Springer, Berlin, 2001), Chap. 3. doi:10.1007/978-3-642-61798-0.
[18] A. Weiße, G. Wellein, A. Alvermann, and H. Fehske, The kernel polynomial method, *Rev. Mod. Phys.* **78**, 275–306 (2006). doi:10.1103/RevModPhys.78.275.
[19] A. Szabo, K. Schulten, and Z. Schulten, First passage time approach to diffusion controlled reactions, *J. Chem. Phys.* **72**, 4350–4357 (1980). doi:10.1063/1.439715.
[20] I. B. Celik, U. Ghia, P. J. Roache, C. J. Freitas, H. Coleman, and P. E. Raad, Procedure for estimation and reporting of uncertainty due to discretization in CFD applications, *J. Fluids Eng.* **130**, 078001 (2008). doi:10.1115/1.2960953.
[21] P. Jizba and V. Zatloukal, Path-integral approach to the Wigner–Kirkwood expansion, *Phys. Rev. E* **89**, 012135 (2014). doi:10.1103/PhysRevE.89.012135.
[22] E. Wigner, On the quantum correction for thermodynamic equilibrium, *Phys. Rev.* **40**, 749–759 (1932). doi:10.1103/PhysRev.40.749.
[23] J. G. Kirkwood, Quantum statistics of almost classical assemblies, *Phys. Rev.* **44**, 31–37 (1933). doi:10.1103/PhysRev.44.31.
[24] National Institute of Standards and Technology, Atomic weights and isotopic compositions. NIST Physical Reference Data.
[25] B. Roux, T. Allen, S. Bernèche, and W. Im, Theoretical and computational models of biological ion

channels, *Q. Rev. Biophys.* **37**, 15–103 (2004). doi:10.1017/S0033583504003968.

[26] A. Parsegian, Energy of an ion crossing a low dielectric membrane: solutions to four relevant electrostatic problems, *Nature* **221**, 844–846 (1969). doi:10.1038/221844a0.

[27] B. Corry, S. Kuyucak, and S.-H. Chung, Dielectric self-energy in Poisson–Boltzmann and Poisson–Nernst–Planck models of ion channels, *Biophys. J.* **84**, 3594–3606 (2003). doi:10.1016/S0006-3495(03)75091-7.

[28] P. Hänggi, P. Talkner, and M. Borkovec, Reaction-rate theory: fifty years after Kramers, *Rev. Mod. Phys.* **62**, 251–341 (1990). doi:10.1103/RevModPhys.62.251.

[29] M. Kac, Can one hear the shape of a drum?, *Am. Math. Mon.* **73**, 1–23 (1966). doi:10.2307/2313748.
